# A general principle of neuronal evolution reveals a human-accelerated neuron type potentially underlying the high prevalence of autism in humans

**DOI:** 10.1101/2024.08.02.606407

**Authors:** Alexander L. Starr, Hunter B. Fraser

## Abstract

The remarkable ability of a single genome sequence to encode a diverse collection of distinct cell types, including the thousands of cell types found in the mammalian brain, is a key characteristic of multicellular life. While it has been observed that some cell types are far more evolutionarily conserved than others, the factors driving these differences in evolutionary rate remain unknown. Here, we hypothesized that highly abundant neuronal cell types may be under greater selective constraint than rarer neuronal types, leading to variation in their rates of evolution. To test this, we leveraged recently published cross-species single-nucleus RNA-sequencing datasets from three distinct regions of the mammalian neocortex. We found a strikingly consistent relationship where more abundant neuronal subtypes show greater gene expression conservation between species, which replicated across three independent datasets covering >10^6^ neurons from six species. Based on this principle, we discovered that the most abundant type of neocortical neurons—layer 2/3 intratelencephalic excitatory neurons—has evolved exceptionally quickly in the human lineage compared to other apes. Surprisingly, this accelerated evolution was accompanied by the dramatic down-regulation of autism-associated genes, which was likely driven by polygenic positive selection specific to the human lineage. In sum, we introduce a general principle governing neuronal evolution and suggest that the exceptionally high prevalence of autism in humans may be a direct result of natural selection for lower expression of a suite of genes that conferred a fitness benefit to our ancestors while also rendering an abundant class of neurons more sensitive to perturbation.

## Introduction

With the advent of single cell RNA-sequencing (scRNA-seq), it became possible to systematically delineate molecularly defined cell types across the brain^1,2^. As more large-scale datasets were published, it quickly became clear that the mammalian brain contains a staggering array of neuronal cell types, with recent whole-brain studies identifying nearly as many neuronal types as there are protein-coding genes in the genome^1–3^. In addition, cross-species atlases in the neocortex revealed that most cortical neuronal types are highly conserved in primates and rodents, with very few neocortical neuronal types being specific to primates and none being entirely specific to humans^4–8^. This suggests that divergence involving homologous cell types—such as their patterns of gene expression, relative proportions, and connectivity— may play a central role in establishing uniquely human cognition.

Two decades before the generation of these cross-species cell type atlases, the first whole-genome sequences of eukaryotes were published, enabling genome-wide studies of evolution for the first time^9^. One of the first questions to be addressed in the nascent field of evolutionary genomics was why some proteins are highly conserved throughout the tree of life, whereas others evolve so quickly as to be almost unrecognizable as orthologs even over relatively short divergence times^10–13^. A protein’s expression level emerged as the strongest and most universal predictor of its evolutionary rate, with highly expressed proteins accumulating fewer protein-coding changes due to greater constraint^10,14–16^.

In contrast to tens of thousands of publications about the evolutionary rates of proteins^17^, the evolutionary rates of cell types, another key building block of multicellular life, have received relatively little attention^18^. Just as different proteins make up every cell, different cell types make up every multicellular organism. Furthermore, just as protein evolutionary rates are measured by the total rate of change of their amino acids, the evolutionary rates of cell types—which are typically defined by their patterns of gene expression—can be measured by divergence in genome-wide gene expression^4–8^. For example, it is well-established that gene expression in neurons is more conserved between humans and mice than gene expression in glial cell types such as astrocytes, oligodendrocytes, and microglia^19^. Previous analogies between genes and neural cell types have been fruitful for understanding the evolution of novel cell types^6,20–23^, providing an encouraging precedent for our analogy.

One area that has been explored more thoroughly is the association of specific cell types with human diseases and disorders^24^. For example, integration of gene-trait associations with cell type-specific expression profiles has revealed that microglia likely play a central role in Alzheimer’s disease^25,26^. Similar analyses have also revealed that layer 2/3 intratelencephalic excitatory (L2/3 IT neurons)—which enable communication between neocortical areas^27^ and are thought to be important for uniquely human cognitive abilities^27,28^—likely play a particularly important role in autism spectrum disorder (ASD) and schizophrenia (SCZ)^29–36^, together with deep layer IT neurons^36–38^. ASD and SCZ are neurodevelopmental disorders with different but overlapping characteristics, including major effects on social behavior^39–41^. Interestingly, individuals with ASD are more likely to be diagnosed with SCZ than individuals without an ASD diagnosis^39,42–44^. Furthermore, there is a strong overlap in the genes that have been implicated in both disorders^36,39^.

From an evolutionary perspective, it has been proposed that ASD and SCZ may be unique to humans^45–47^. This is primarily based on two main lines of reasoning. First, ASD- and SCZ-associated behaviors that could reasonably be observed in non-human primates (e.g. SCZ-associated psychosis) have been observed either infrequently or not at all in non-human primates^46^. However, ASD-like behavior has been observed in non-human primates^48^ and the difficulties inherent to cross-species behavioral comparisons combined with relatively low sample sizes make it difficult to compare the prevalence of these behaviors in human and non-human primate populations. Second, core ASD- and SCZ-associated behavioral differences involve cognitive traits that are either unique to or greatly expanded in humans (e.g. speech production and comprehension or theory of mind)^49–53^. As a result, certain aspects of ASD and SCZ are inherently unique to humans.

While comparing interindividual behavioral differences across species remains challenging, recent molecular and connectomic evidence lend credence to the idea that the incidence of ASD and SCZ increased during human evolution. For example, large-scale sequencing studies in both ASD and SCZ cohorts have identified an excess of genetic variants in human accelerated regions (HARs)—genomic elements that were largely conserved throughout mammalian evolution but evolved rapidly in the human lineage^54–56^. Furthermore, transcriptomic studies have identified a human-specific shift in the expression of some synaptic genes during development that is disrupted in ASD^57^. In addition, connectomic studies have shown that human-chimpanzee divergence in brain connectivity overlaps strongly with differences between humans with and without SCZ^58^. Overall, evidence suggests that ASD and SCZ may be particularly prevalent in humans, but the factors underlying this increased prevalence remain unknown. Positive selection—also known as adaptive evolution—of brain-related traits in the human lineage has been proposed to underlie this increase^45–47,59,60^. Although this idea is supported by the links between HARs (many of which are thought to have been positively selected^56^) and ASD and SCZ, there is no direct evidence for positive selection on the expression of genes linked to ASD and SCZ.

Here, we set out to test whether the inverse relationship between abundance and evolutionary rates—which has been well-established for proteins^10,14–16^—might also hold for cell types. We found a robust negative correlation between cell type proportion and evolutionary divergence in the neocortex, suggesting that this relationship holds at multiple levels of biological organization. Based on this, we identify unexpectedly rapid evolution of L2/3 IT neurons and strong evidence for polygenic positive selection for reduced expression of ASD-linked genes in the human lineage, suggesting that positive selection may have increased the prevalence of ASD in modern humans.

## Results

### Cell type proportion as a general factor governing the rate of neuronal evolution

Based on the gene-cell type analogy outlined above, we hypothesized that a change in gene expression in a more abundant cell type may tend to have more negative fitness effects than the same change in a less abundant cell type (Figure 1A). If this were the case, this would lead to greater selective constraint, and thus slower divergence, of global gene expression in more abundant cell types.

**Figure 1:**
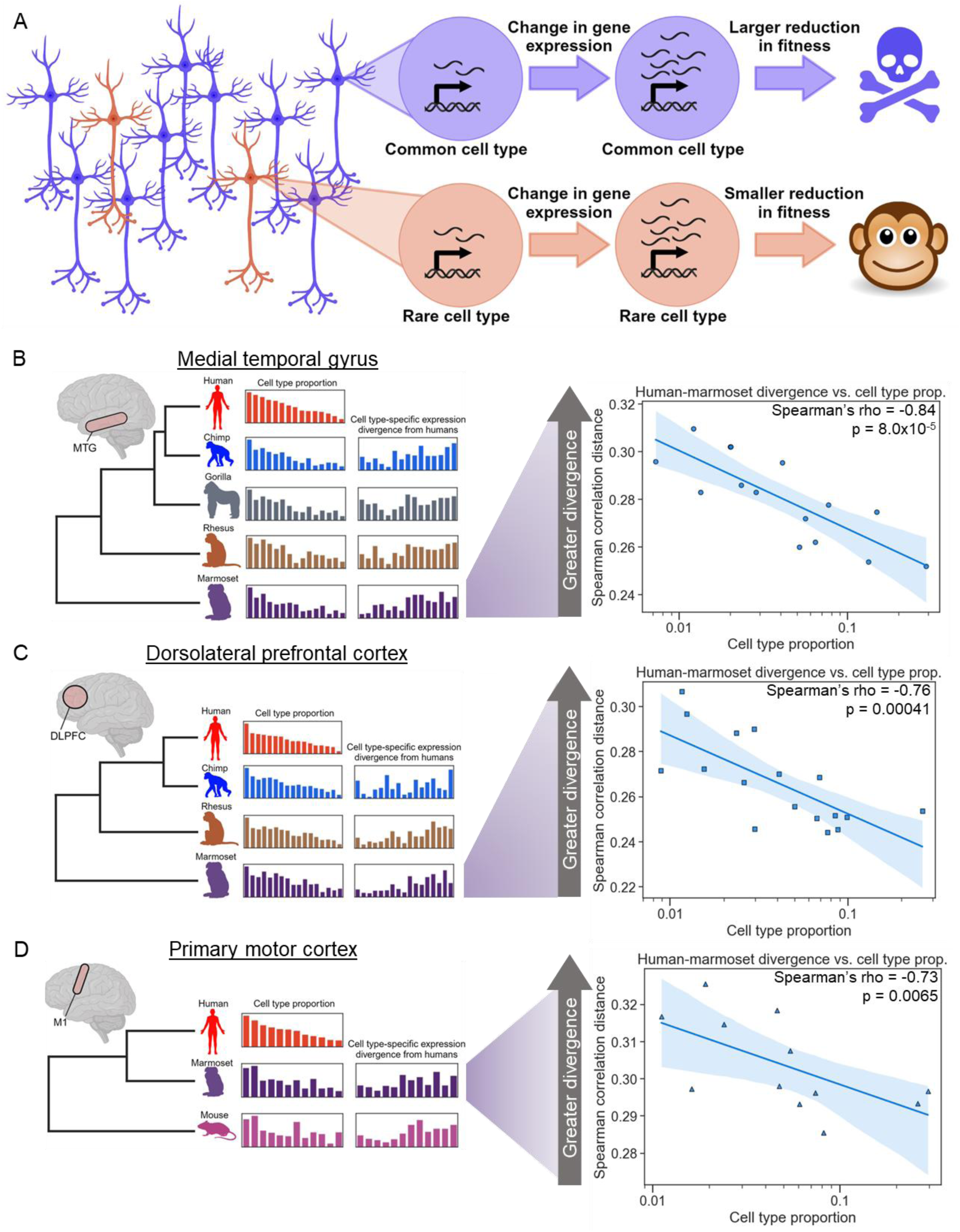
More common neuronal cell types evolve more slowly than rare types. **A)** Rationale for hypothesis that more common neuronal types might evolve more slowly than rarer types. A gene expression change in a common cell type has a large negative effect on fitness whereas the same change in a rarer cell type has a smaller effect. **B) Left:** outline of our data analysis strategy. SnRNA-seq from the MTG of five species (14 subclasses of neuron) was used to estimate each cell type’s proportion and pairwise divergence between species. **Right:** plot showing the correlation between neuronal subclass proportion (log_10_ scale on the x-axis) and subclass-specific divergence between human and marmoset in the MTG. A representative iteration from 100 independent down-samplings is shown. The Spearman’s rho and p-value shown are the median across 100 independent down-samplings (see Methods for details). The line and shaded region are the line of best fit from a linear regression and 95% confidence interval respectively. **C)** Same as (B) but snRNA-seq from the DLPFC (17 subclasses of neuron) of four species was analyzed. **D)** Same as (B) but snRNA-seq from M1 (12 subclasses of neuron) of three species was analyzed.

Testing this hypothesis requires comparing two quantities: cell type proportions and the evolutionary divergence in genome-wide gene expression levels between orthologous cell types across species. Importantly, both quantities can be estimated from the same single-nucleus RNA-seq (snRNA-seq) data, facilitating comparison between them. To ensure sufficient statistical power, we searched the literature for published snRNA-seq data sets that fulfilled a stringent pair of criteria. First, they must have multiple species profiled in the same study using the same snRNA-seq protocols for each species within a study. Second, they must contain at least 10 orthologous cell types having 250 or more cells per species (not including immune cells, as these do not have stable cell type proportions). We identified three studies fulfilling these criteria, focused on three distinct regions of the mammalian neocortex: medial temporal gyrus (MTG), dorsolateral prefrontal cortex (DLPFC), and primary motor cortex (M1)^5,7,8^. All three studies included samples from 3-5 species, including human and marmoset, with 300,000 – 500,000 neuronal nuclei profiled per study^5,7,8^. These nuclei were clustered into between 12 – 17 neuronal subclasses (with at least 250 cells per species) in each study, which we then used for our analyses^5,7,8^. Throughout, we use the term cell type for the general concept of different types of cells and as an umbrella term for both subclasses and subtypes, use the term subclass for the traditional classification of neuronal types found in the neocortex, and reserve the term subtype for more fine-grained clustering of cells.

To test our hypothesis, we began by comparing human and marmoset (the only pair of species present in all three datasets) in the MTG, which had the greatest sequencing depth. We first estimated gene expression divergence for each of 14 subclasses using the Spearman correlation distance (1 – Spearman’s rho) between the pseudobulked expression of each species for each neuron subclass, restricting to one-to-one orthologous genes (see Methods). We observed a surprisingly strong negative correlation between subclass proportion and gene expression divergence (Spearman’s rho = -0.84, p = 8.0×10^-5^, Figure 1B), indicating that more abundant neuronal subclasses showed greater conservation of genome-wide gene expression. To ensure that estimates of cell type-specific expression divergence were not biased by cell type proportion itself, we analyzed the same number of cells and total reads for each cell type in each species. Specifically, for all analyses we report the median rho and p-values from 100 independent down-samplings of cells and pseudobulked counts without replacement (see Methods).

We next asked whether the same pattern was present in the other cortical regions. We observed a similar strong negative correlation in the two other independently generated datasets (Spearman’s rho = -0.76, p = 0.00041 in the DLPFC, Figure 1C; Spearman’s rho = - 0.73, p = 0.0065 in the M1, Figure 1D). This replication suggests that the relationship we observed holds true across the primate neocortex. In addition, the fact that methodological details and biological samples differ across these studies lends additional robustness to any patterns shared by all three.

To explore the generality of this result in additional species, we repeated this analysis between every pair of species in each dataset. We observed similarly strong negative correlations across all pairwise comparisons (Supplemental Figures 1-3), with the interesting exception of comparisons between humans and non-human great apes, where a weaker negative correlation was observed (discussed below). Furthermore, we observed strong negative correlations within excitatory or inhibitory subclasses in all three brain regions (Figure 2 and Supplemental Figures 4-9, although this correlation does not reach statistical significance for inhibitory neurons in M1, potentially due to having only five subclasses in that dataset). In addition, we tested all possible combinations of a wide variety of filtering parameters, analysis decisions, and distance metrics, finding that this negative correlation was generally robust to any reasonable choice of parameters we made (Supplemental Table 1).

**Figure 2:**
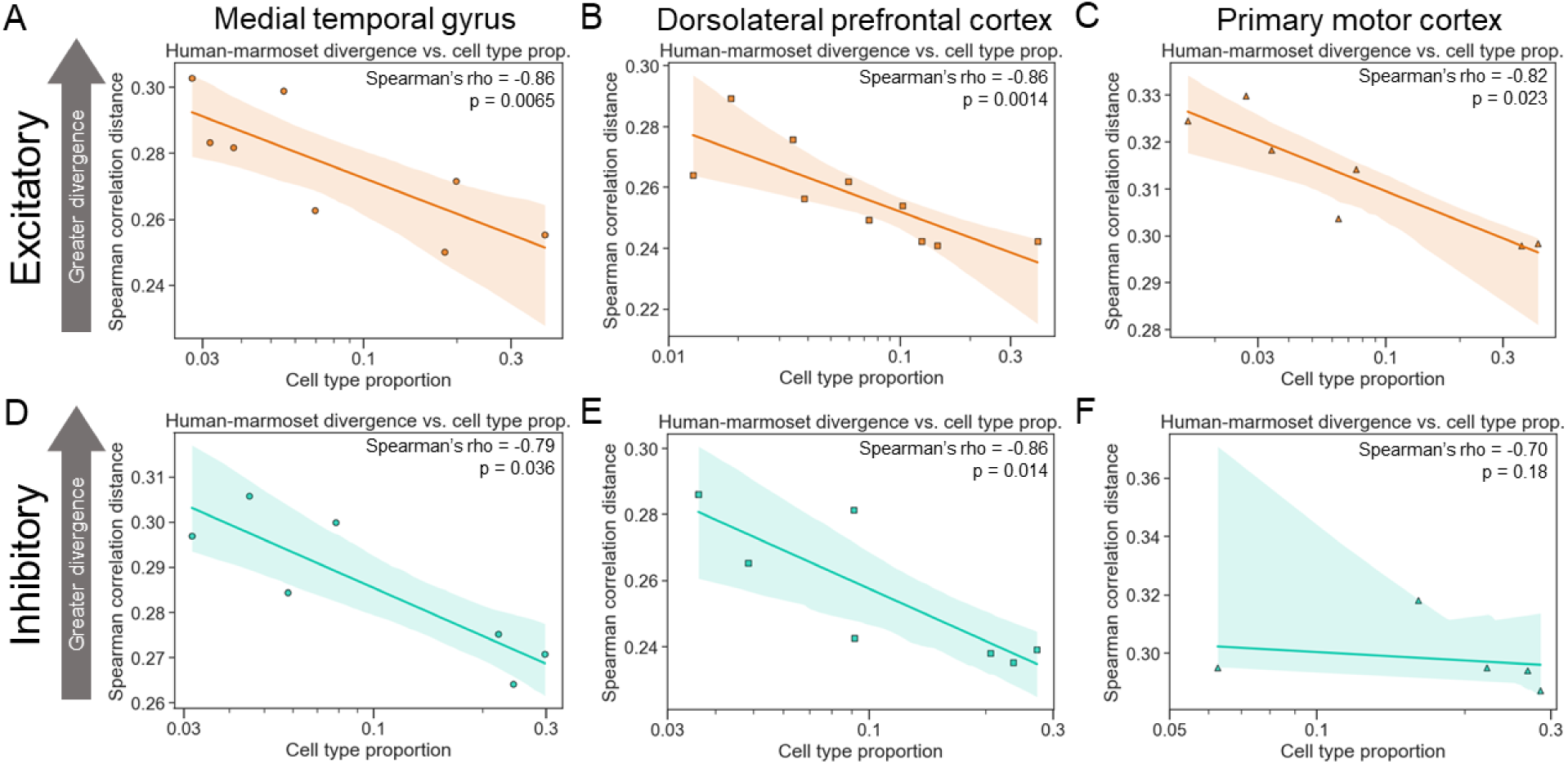
More common neuronal cell types evolve more slowly than rare types within excitatory and inhibitory classes. **A)** Plot showing the correlation between neuronal subclass proportion (log_10_ scale on the x-axis) and subclass-specific divergence between human and marmoset for excitatory neurons in the MTG. A representative iteration from 100 independent down-samplings is shown. The Spearman’s rho and p-value shown are the median across 100 independent down-samplings (see Methods for details). The line and shaded region are the line of best fit from a linear regression and 95% confidence interval respectively. **B)** Same as in (A) but for the DLPFC data. **C)** Same as in (A) but for the M1 data. **D)** Same as in (A) but for inhibitory neurons. **E)** Same as in (B) but for inhibitory neurons. **F)** Same as in (C) but for inhibitory neurons.

Next, we investigated this relationship at the level of neuronal subtypes, a finer-grained clustering with ∼4-fold more cell subtypes than subclasses. We found strong negative correlations between subtype proportion and expression divergence when using all neurons (Figure 3A-C, Supplemental Figures 10-12) or only excitatory neurons (Figure 3D-F, Supplemental Figures 13-15). When restricting our analysis to inhibitory neurons, this correlation was statistically significant in the MTG and in two of three comparisons (mouse-marmoset and human-mouse) in the M1, but not in DLPFC (Figure 3G-I, Supplemental Figures 16-18). This may reflect the lower read depth (average of 180,054 counts used for DLPFC, compared to 254,703 for M1 and 325,422 for MTG) or lower numbers of cells per subtype in the DLPFC data compared to the other datasets, as we observed a much stronger negative correlation (Spearman’s rho = -0.50, p = 0.057) when restricting to subtypes with at least 500 cells in the DLPFC data (Supplemental Figure 19). Overall, our results suggest that there is a strong, robust negative correlation between expression divergence and cell type proportion for neocortical neurons.

**Figure 3:**
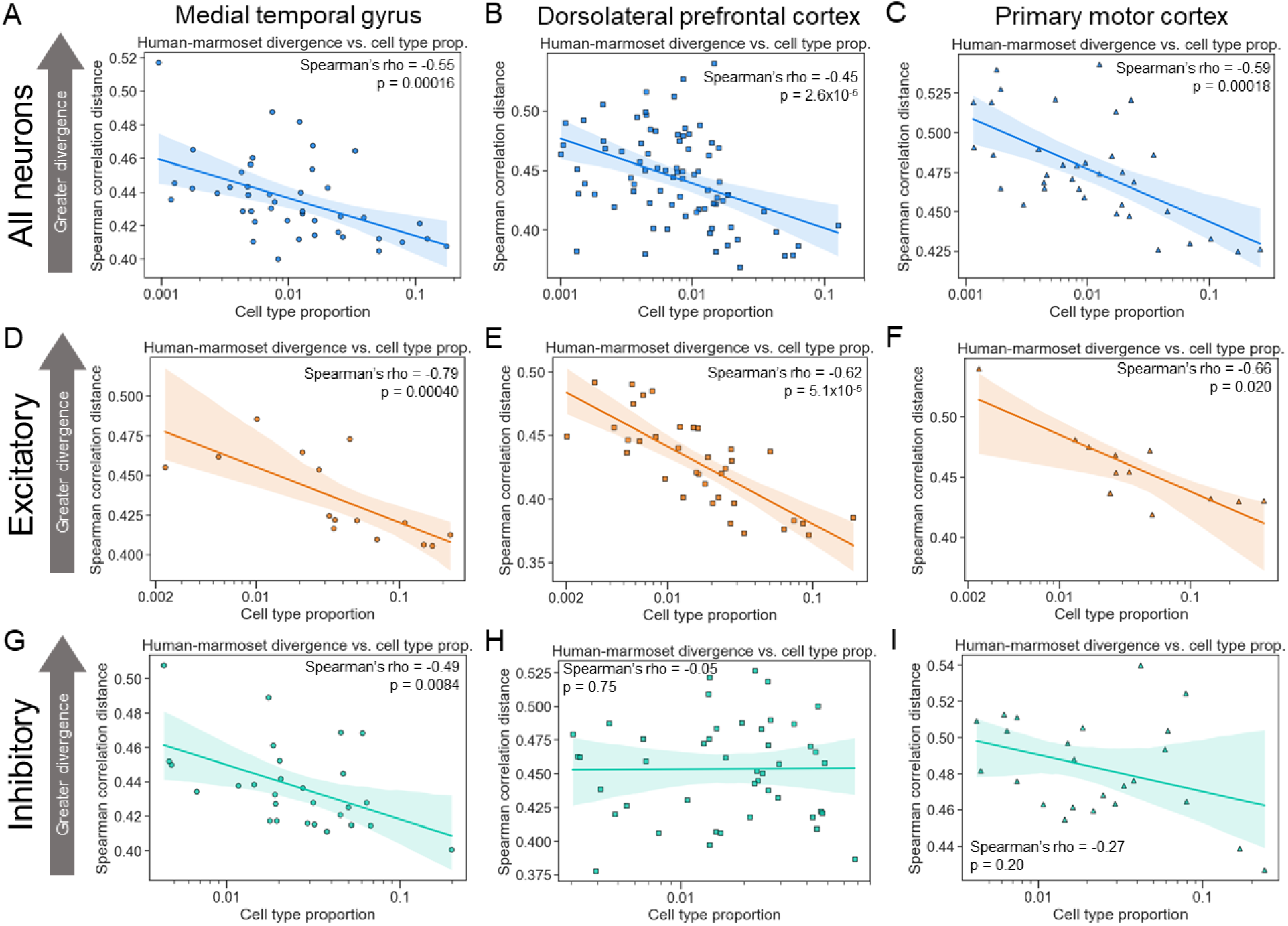
More common neuronal cell types evolve more slowly than rarer types at the subtype level. **A)** Plot showing the correlation between neuronal subtype proportion (log_10_ scale on the x-axis) and subtype-specific divergence between human and marmoset in the MTG. A representative iteration from 100 independent down-samplings is shown. The Spearman’s rho and p-value shown are the median across 100 independent down-samplings (see Methods for details). The line and shaded region are the line of best fit from a linear regression and 95% confidence interval respectively. **B)** Same as in (A) but for the DLPFC data. **C)** Same as in (A) but for the M1 data. **D)** Same as in (A) but for excitatory neurons. **E)** Same as in (B) but for excitatory neurons. **F)** Same as in (C) but for excitatory neurons. **G)** Same as in (A) but for inhibitory neurons. **H)** Same as in (B) but for inhibitory neurons. **I)** Same as in (C) but for inhibitory neurons.

Finally, we investigated the properties of the genes driving the negative correlation we observed. First, we stratified genes into three equally sized bins by their expression level and recomputed correlations in each bin. Interestingly, while we observed strong correlations for highly and moderately expressed genes, there was no significant correlation when restricting to lowly expressed genes (Figure 4A, Supplemental Figures 20-22, Supplemental Table 2). Next, we stratified genes based on evolutionary constraint on expression level or cell type-specificity of expression (using s_het_^61^ and the Tau metric^62^ respectively, Supplemental Tables 3 and 4).

**Figure 4:**
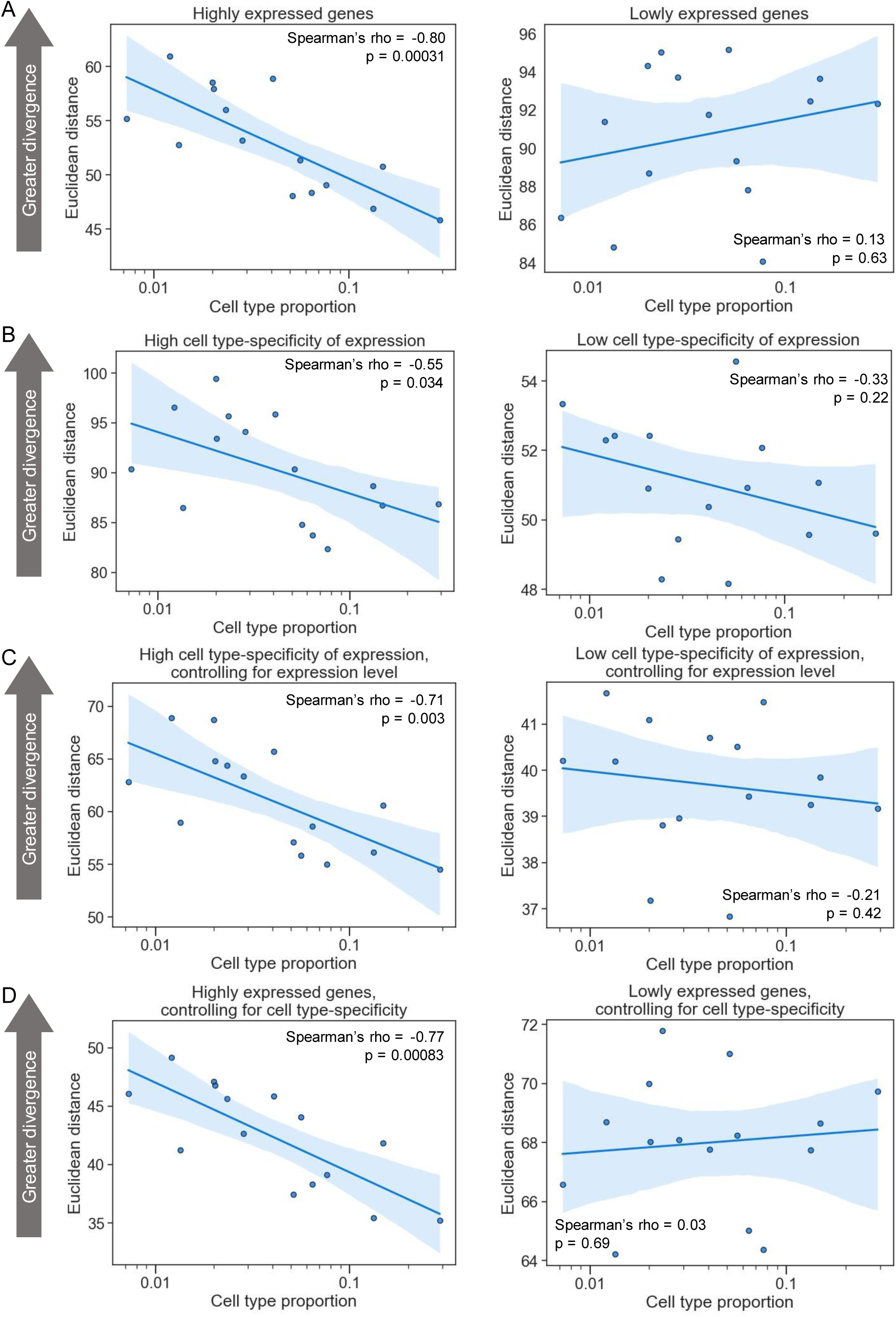
More highly expressed, cell type-specific genes drive the negative correlation between cell type proportion and evolutionary divergence. **A) Left:** Plot showing the correlation between neuronal subtype proportion (log_10_ scale on the x-axis) and subtype-specific divergence for highly expressed genes between human and marmoset in the MTG. A representative iteration from 100 independent down-samplings is shown. The Spearman’s rho and p-value shown are the median across 100 independent down-samplings (see methods for details). The line and shaded region are the line of best fit from a linear regression and 95% confidence interval respectively. **Right:** Same as the left but for lowly expressed genes. **B) Left:** Same as in (A) but for genes with more cell type-specific expression. **Right:** Same as left but for genes with less cell type-specific expression. **C)** Same as in (B) but controlling for expression level (see Methods). **D)** Same as in (A) but controlling for cell type-specificity of expression (see Methods).

While there was no difference in correlation when stratifying by constraint on expression (Supplemental Figures 23-25, Supplemental Table 3), we observed a much stronger negative correlation between cell type proportion and expression divergence for more cell type-specifically expressed genes (Figure 4B, Supplemental Figures 26-28, Supplemental Table 4). Since expression level is also associated with cell-type specificity, we tested whether these two properties were contributing independently to the negative correlations by stratifying genes by one of them while simultaneously controlling for the other. We found that both properties retained their predictive power even when controlling for the other (Figure 4C-D, Supplemental Figures 29-34, Supplemental Tables 2 and 4), suggesting independent contributions. We note that whether the weaker correlations we observed for lowly expressed genes were due to a true lack of association or simply less accurate expression level measurements remains an open question that will require larger datasets to explore. Overall, our results suggest that more highly expressed, cell type-specific genes are primarily driving the negative correlation between cell type proportion and gene expression divergence.

### Rapid evolution of layer 2/3 intratelencephalic neurons in the human lineage

Having identified this strong relationship between cell type proportion and evolutionary divergence, we reasoned that cell types with much faster divergence in the human lineage than expected based on their abundance may have been subject to atypical selective forces.

To identify subclasses showing the most dramatic lineage-specific shifts in selection, we decomposed human-chimpanzee MTG expression divergence into its two components, divergence on the human branch and divergence on the chimpanzee branch. Applying the concept of parsimony—explaining the data with as few evolutionary transitions as possible— allows an outgroup species such as gorilla to polarize changes and assign them to either the human or chimpanzee branch (see Methods). In the chimpanzee lineage, there was a strong negative correlation between divergence and subclass proportion (Figure 5A, Spearman’s rho = -0.77, p = 0.00076), similar to the correlations between other primate species (Figure 1A, Supplemental Figure 1). However, we observed a much weaker negative correlation in the human lineage (Figure 5B, Spearman’s rho = -0.19, p = 0.49). The clearest outlier weakening the correlation was L2/3 IT neurons, the most abundant neuronal subclass, which diverged much faster than expected based on its proportion. This was also true to a lesser extent for the next two most abundant subclasses, L4 IT and L5 IT neurons. Indeed, removing these three subclasses substantially strengthened the negative correlation between subclass proportion and human-specific divergence (Figure 5B; Spearman’s rho = -0.59, p = 0.041), making it indistinguishable from the corresponding chimpanzee-specific correlation (Figure 5A, blue points; Spearman’s rho = -0.58, p = 0.048). Quantifying the magnitude of human acceleration for every subclass confirmed that L2/3 IT neurons underwent the greatest acceleration, followed by L4 and L5 IT neurons (Figure 5C).

**Figure 5:**
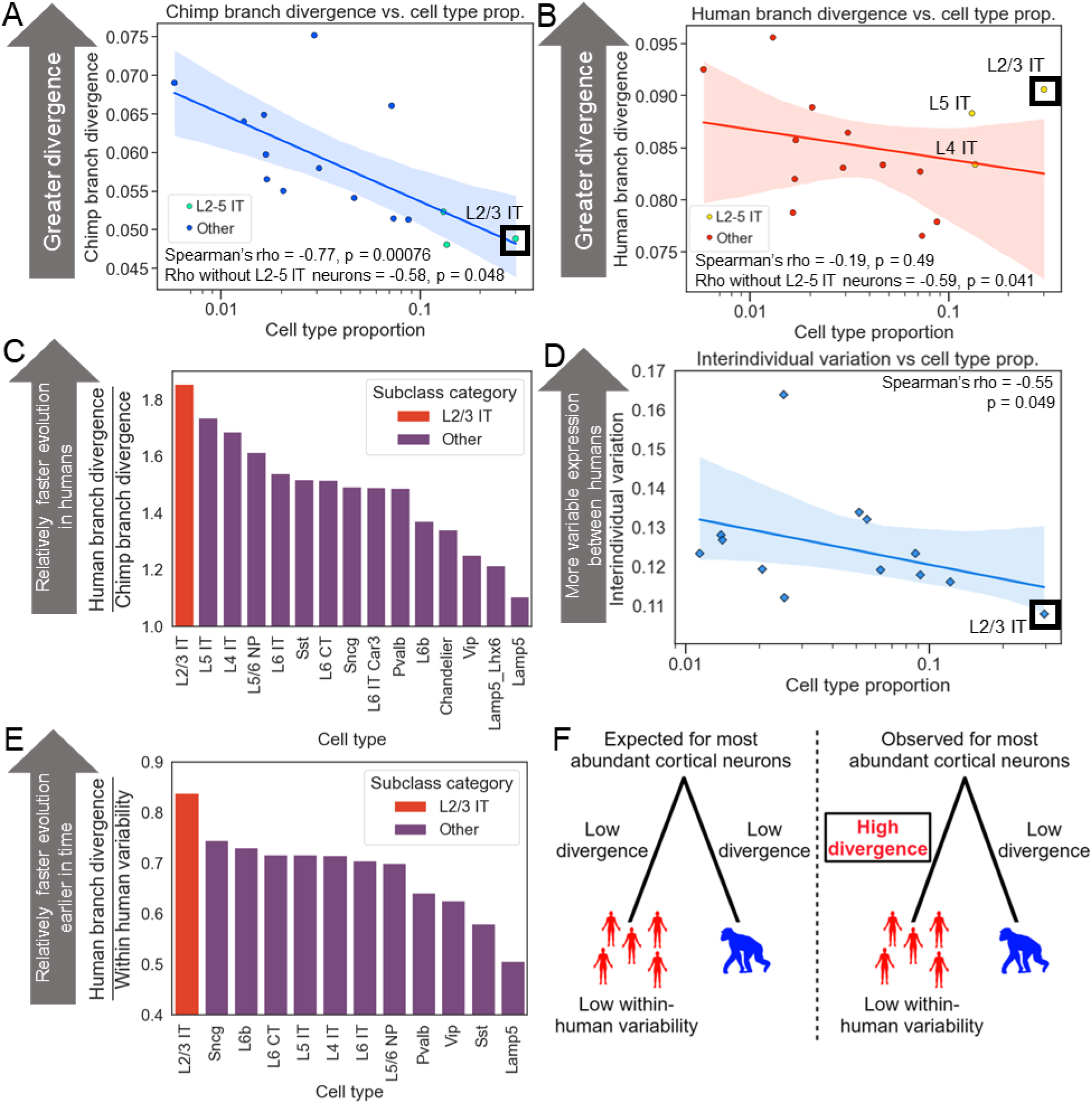
Accelerated evolution of L2/3 IT neurons in the human lineage. **A)** Plot showing the correlation between neuronal subclass proportion (log_10_ scale on the x-axis) and subclass-specific divergence on the chimpanzee branch in the MTG. Chimpanzee branch divergence was computed for each of 100 down-samplings and the mean across those down-samplings is shown. The line and shaded region are the line of best fit from a linear regression and 95% confidence interval respectively. Teal points indicate L2-5 IT neurons. **B)** Same as in (A) but for human branch divergence. Yellow points indicate L2-5 IT neurons. **C)** Barplot showing the human branch divergence divided by the chimpanzee branch divergence for each subclass. **D)** Plot showing the correlation between neuronal subclass proportion (log_10_ scale on the x-axis) and subclass-specific interindividual variation across DLPFC samples from 25 human individuals. A representative iteration from 100 independent down-samplings is shown. The Spearman’s rho and p-value shown are the median across 100 independent down-samplings (see Methods for details). The line and shaded region are the line of best fit from a linear regression and 95% confidence interval respectively. **E)** Barplot showing the human branch divergence divided by the within-human variability for each subclass. **F)** Conceptual model for accelerated evolution of L2/3 IT neurons in the human lineage.

Accelerated evolution can involve either positive selection favoring gene expression changes that increased fitness, or relaxed selective constraint in which random mutations are allowed to accumulate over time because they have little or no effect on fitness^56^. Although both positive selection and relaxed constraint can lead to similar patterns of lineage-specific acceleration, they imply very different underlying factors: positive selection is the force underlying nearly all evolutionary adaptation, while relaxed constraint is simply the weakening or absence of natural selection which can lead to the passive deterioration of genes and their regulatory elements via mutation accumulation.

To distinguish whether positive selection or relaxed constraint was more likely to underlie the human-specific acceleration of IT neurons, we investigated the interindividual variability in expression of each neuronal subclass in the human population^63^. If IT neurons evolved under reduced constraint in the human lineage then we would expect them to have more variable expression among humans, leading to a weaker negative correlation between subclass proportion and interindividual variability. Instead, we observed a strong negative correlation between subclass proportion and interindividual variability in gene expression, with L2/3 IT neurons having the lowest variability of any subclass among humans (Figure 5D, Spearman’s rho = -0.55, p = 0.049). Consistent with this, L2/3 IT neurons had the largest human branch divergence relative to their expression variability in modern humans (Figure 5E). Overall, these results suggest that the rapid gene expression evolution of L2/3 IT neurons in the human lineage was unlikely to be due to relaxed constraint, and instead more likely the result of positive selection (Figure 5F), though we cannot formally rule out other possible scenarios (see Discussion). In addition, it suggests that the relationship between cell type proportion and expression divergence holds within species as well as between species.

### Lower expression of ASD-linked genes in humans compared to chimpanzees

Our finding of human-specific accelerated evolution of L2/3 IT neurons raised the question of what phenotypes may be most affected by this. To explore this, we tested gene sets with strong evidence of linkage to specific human traits for bias toward higher or lower expression in humans relative to chimpanzees in L2/3 IT neurons. These gene sets were derived from two sources: the Human Phenotype Ontology (HPO)^64^, a broad database covering hundreds of human traits, and SFARI, an ASD-specific database. Although ASD is often influenced by common genetic variants of small effect, which can be identified by GWAS, it can also be caused by single large effect variants typically causing loss-of-function of a core^65^ ASD gene. The SFARI database is the most comprehensive collection of these core genes^66^; we refer to SFARI genes with a score of 1 as “high-confidence ASD-linked” and all SFARI genes, regardless of score, as “ASD-linked”.

Strikingly, we found that high-confidence ASD-linked genes showed a stronger directionality bias in L2/3 IT neurons than any of the 359 HPO gene sets tested (4.0-fold enrichment for lower expression in human MTG and 4.3-fold enrichment in DLPFC; p < 10^-7^ for each; Figure 6A, Supplemental Figure 35A). Although some HPO gene sets were also enriched, this was mostly a result of pleiotropic ASD-linked genes being present in multiple gene sets (Supplemental Figure 35B-C). This strong and specific enrichment for lower expression of high-confidence ASD-linked genes in human L2/3 IT neurons was intriguing, considering the known role of these neurons in ASD^30–34^.

**Figure 6:**
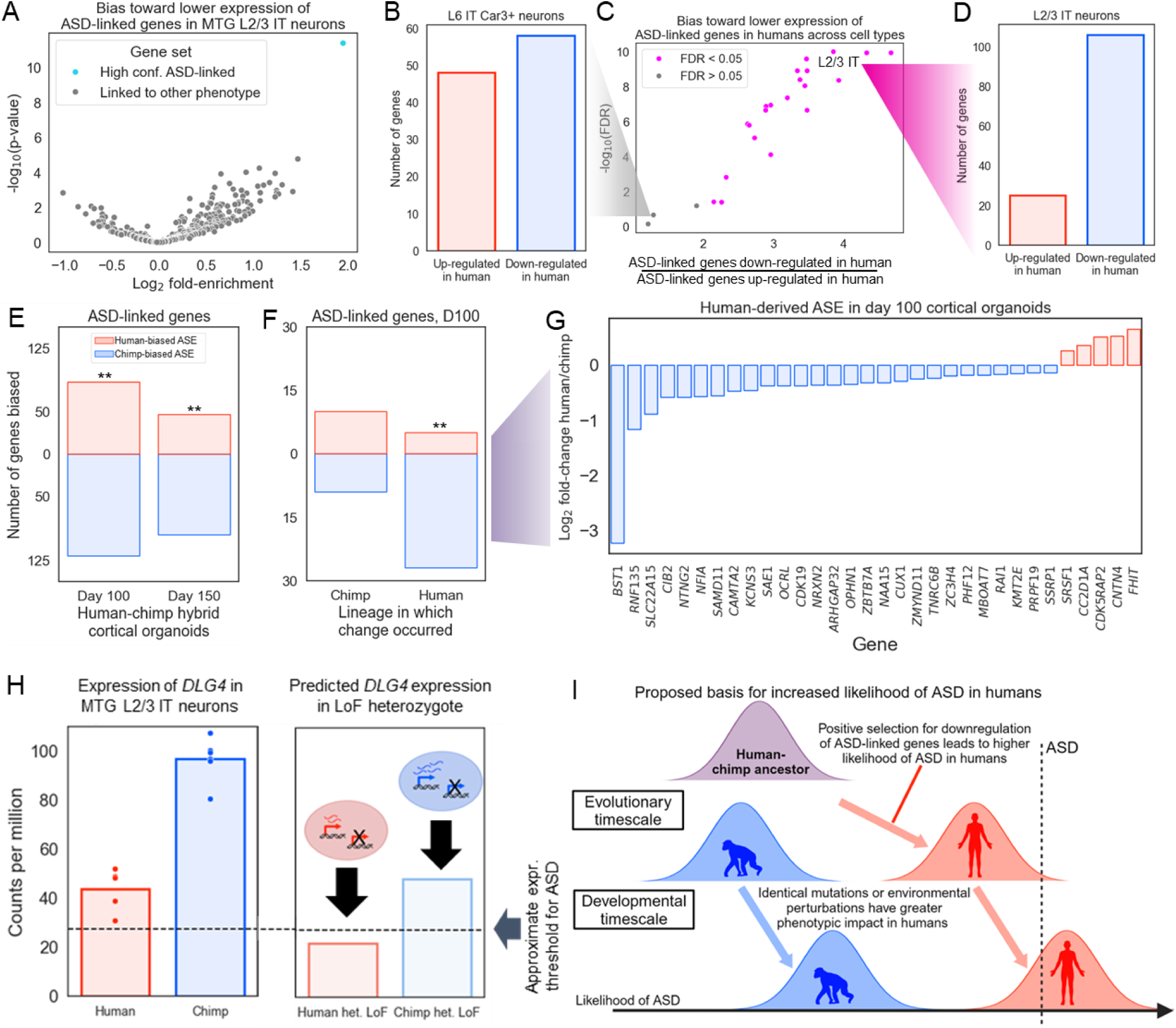
Positive selection for down-regulation of ASD-linked genes in the human lineage. **A)** Volcano plot showing the log_2_ fold-enrichment for down-regulation in humans (x-axis) and the -log_10_ binomial p-value (y-axis). SFARI high-confidence ASD-linked genes are shown in blue, all other categories of genes (taken from the Human Phenotype Onotology) are shown in grey. Data is from MTG L2/3 IT neurons. **B)** Barplot showing the number of high-confidence ASD-linked genes that are up-regulated vs. down-regulated in human relative to chimpanzee in MTG L6 IT Car3+ neurons. **C)** Plot showing the fold-enrichment for down-regulation in human MTG (x-axis) and the -log_10_ binomial FDR (y-axis). Subclasses with FDR < 0.05 are shown in magenta; only subclasses with at least 500 human vs. chimpanzee differentially expressed genes in each direction are shown. **D)** Barplot showing the number of high-confidence ASD-linked genes that are up-regulated vs. down-regulated in human relative to chimp in MTG L2/3 IT neurons. **E)** Barplot showing the number of differentially expressed ASD-linked genes with higher allele-specific expression from the human allele (red) and higher expression from the chimpanzee allele (blue) in cortical organoids. ** indicates binomial p < 0.01. **F)** Barplot showing the number of differentially expressed ASD-linked genes with higher allele-specific expression from the human allele (red) and higher expression from the chimpanzee allele (blue) in day 100 cortical organoids for human-derived and chimpanzee-derived genes separately. ** indicates binomial p < 0.01. **G)** Plot showing the log_2_ allele-specific expression ratios of differentially expressed, human-derived, ASD-linked genes in day 100 cortical organoids. **H)** Left: Expression of *DLG4* in MTG L2/3 IT neurons. Right: Predicted expression of *DLG4* if one copy of the gene were non-functional. **I)** Conceptual model for how positive selection for down-regulation of ASD-linked genes led to higher likelihood of ASD in humans compared to chimpanzees.

We then asked whether this lower expression of high-confidence ASD-linked genes was shared in other neuronal types beyond L2/3 IT. We found that some types of neurons had no significant directionality bias (Figure 6A), while many subclasses shared a bias towards lower expression of high-confidence ASD-linked genes in humans compared to chimpanzees (Fig 6B). In both the DLPFC and the MTG datasets we observed the most significant trend towards lower human expression of these genes in L2/3 IT neurons (Figure 6B-C, Supplemental Figure 36A-C).

This excess of high-confidence ASD-linked genes with lower expression in humans is consistent with either down-regulation in the human lineage, up-regulation in the chimpanzee lineage, or a combination of both. To distinguish between these possibilities, we used gorilla as an outgroup to assign each gene’s expression divergence in the MTG to either the human or chimpanzee lineage.

Comparing the expression of high-confidence ASD-linked genes in all three species revealed that gorilla gene expression is significantly closer to chimpanzee, suggesting that there has been greater divergence in the human lineage (Supplemental Figure 37A). Consistent with this, a significantly larger number of high-confidence ASD-linked genes’ expression diverged on the human branch than expected by chance in L2/3 IT neurons (Supplemental Figure 37B). In addition, human L2/3 IT neurons have overall lower expression of these genes as compared to all four NHPs in the dataset (Supplemental Figure 37C, Supplemental Table 5). Overall, these results suggest a consistent pattern of human-specific down-regulation of ASD-associated genes in a neuronal cell type with a key role in ASD.

### Polygenic positive selection for down-regulation of ASD-linked genes in the human lineage

This human-specific down-regulation of high-confidence ASD-linked genes is striking and, based on the highly constrained expression of these genes, likely functionally significant. However, as with the accelerated evolution of L2/3 IT neurons discussed above (Figure 5), the question of whether lineage-specific selection was responsible is key to understanding the factors that drove this divergence in the human lineage. Other potential explanations fall into two main categories. One is genetic changes that were not driven by selection, such as mutations that had little effect on fitness but became established in the human lineage through genetic drift. The other is non-genetic differences in the individuals sampled for these data sets; factors such as diet, environmental exposures, and age can impact gene expression but cannot be controlled in any comparison of tissue samples between humans and other species.

In order to definitively implicate lineage-specific selection, two steps are necessary. First, all non-genetic causes must be ruled out. Although this is not possible with tissue samples, it can be achieved *in vitro*. Human and chimpanzee induced pluripotent stem cells (iPSCs) can be fused to generate hybrid tetraploid iPSCs, which can then be differentiated into relevant cell types or organoids^67,68^. In each hybrid cell, the human and chimpanzee genomes share precisely the same intracellular and extracellular environment. As a result, any difference in the relative expression levels of the human and chimpanzee alleles for the same gene—known as allele-specific expression (ASE)—reflects *cis*-regulatory changes between the two alleles. Both environmental and experimental sources of variability (including batch effects) are perfectly controlled in the hybrid system, since all comparisons are between alleles that share an identical environment and are present in the same experimental samples^67,68^.

The second step necessary to infer lineage-specific selection is to test, and reject, a statistical “null model” of neutral evolution for the genetic component of divergence^69^. The simplest and most robust pattern predicted under neutral evolution of gene expression is the expectation that in a comparison between two species, genetic variants causing expression divergence will be just as likely to lead to higher expression in one species as in the other^70^. For example, in a set of 20 functionally related genes, neutral evolution leads to a similar pattern as a series of 20 coin flips—an expectation of ∼10 genes more highly expressed in one species and ∼10 in the other, with deviation from this average following the binomial distribution^70^. In contrast, natural selection that favors lower expression of these genes in one lineage will lead to a pattern of biased expression, with most of the 20 genes expressed lower in that lineage^70^. This framework, which has been applied extensively to gene expression and other quantitative traits^67–69,71,72^, is known as the sign test. Because the ASE of each gene in hybrid cells is generally independent of that of other genes, facilitating statistical analysis, hybrid ASE is ideally suited for detecting selection with the sign test whereas data from non-hybrids cannot be used in this manner.

To apply this test for lineage-specific selection, we focused on a previously published RNA-seq dataset from human-chimpanzee hybrid cortical organoids^67^. These organoids—which include glutamatergic and GABAergic neurons, astrocytes, and neural precursor cells—were sampled in a bulk RNA-seq time series of development *in vitro*^67^; we focus on the two timepoints, day 100 and day 150, with the highest proportion of neurons. As described above, a significant bias in the directionality of ASE for any predefined set of genes can reject the null hypothesis of neutral evolution, and instead suggests lineage-specific selection. Applying this test to ASD-linked genes, we found a strong bias toward lower expression from the human allele in cortical organoids at two different stages of development (2.0-fold enrichment at day 100 of organoid development; binomial p = 0.003; Figure 6E). The bias toward lower expression from human alleles was even stronger when using only high-confidence ASD-linked genes (2.5 fold-enrichment; binomial p = 0.01 at day 100; Supplemental Figure 38). This ASE bias is inconsistent with neutral evolution, and strongly implies the action of lineage-specific selection on the expression of ASD-linked genes.

To determine the lineage (human or chimpanzee) on which the ASD-linked gene expression changes occurred, for genes with matching directionality in the L2/3 IT and organoid data we once again polarized gene expression divergence in the MTG into human-derived and chimpanzee-derived categories using gorilla as an outgroup. Out of 17 chimpanzee-derived genes, there was no directionality bias in the organoid ASE data at either day 100 or day 150 (9 out of 19 with lower expression from the human allele at day 100, Figure 6F-G, Supplemental Figure 39), consistent with neutral evolution. However, out of 32 human-derived genes, 27 had lower expression from the human allele (Fisher’s exact test p = 0.010 at day 100, odds ratio = 6.0; p = 0.010, odds ratio = 8.9 at day 150; Figure 6F-G, Supplemental Figure 39;). This trend is even stronger when using a more relaxed false discovery rate (FDR) cutoff of 0.1 (34 down-regulated in human vs 5 up-regulated; Fisher’s exact test p = 0.0043, odds ratio = 5.9; p = 0.0017, odds ratio = 12.5 at day 150). Overall, this strongly suggests that many ASD-linked genes were down-regulated specifically in the human lineage.

This coordinated down-regulation of 34 ASD-linked genes could conceivably be due to either positive selection or loss of constraint, as both of these types of lineage-specific selection could lead to down-regulation^70,72^. To determine if ASD-linked genes might be evolving under relaxed constraint in humans, we tested several predictions of the relaxed constraint model. First, genes evolving under relaxed constraint might be expected to have accumulated more substitutions affecting protein sequence and/or gene expression in the human lineage. However, we found no difference in protein sequence constraint (measured by dN/dS^73^) or the number of mutations near the transcription start site (TSS) between humans and chimpanzees (after correcting for genome-wide differences between the two lineages, p = 0.42 for dN/dS, p = 0.24 for mutations near TSS, paired t-test, Supplemental Figure 40A-B). In addition, the expression of genes evolving under relaxed constraint in humans would likely be more variable across human individuals compared to chimpanzee individuals. However, we found the opposite for ASD-linked genes—slightly less variability in expression in humans (p = 0.08 for DLPFC, p = 2.5×10^-5^ for MTG, paired t-test, Supplemental Figure 40C-D), suggesting that the expression of ASD-linked genes may actually be under stronger constraint in humans compared to chimpanzees.

Consistent with this, the vast majority of ASD-linked genes have strongly constrained expression in humans as measured by loss-of-function intolerance (82% of ASD-linked genes have probability of loss of function intolerance^74^ > 0.9 compared to 17% genome-wide; similarly, 82% of ASD-linked genes have a fitness effect of heterozygous loss of function^61^ [s_het_] > 0.1, compared to 18% genome-wide).

Next, we explored whether particular subsets of ASD-linked genes had a stronger bias toward down-regulation than other ASD-linked genes. ASD-linked genes tend to encode proteins that localize to the synapse, encode transcription factors (TFs) or chromatin remodelers (CRs), and/or be haploinsufficient^75^. When splitting ASD-linked genes into these three partially overlapping categories, we found comparable human down-regulation in all groups (Supplemental Figure 41). For example, 83% of ASD-linked haploinsufficient genes were down-regulated, which is similar to the 75% of ASD-linked non-haploinsufficient genes that were down-regulated (Supplemental Figure 41A). This suggests that ASD-linked genes in general, rather than one of these specific subcategories, are biased toward down-regulation. Finally, we tested whether synaptic genes, TFs/CRs, or haploinsufficient genes in general tend to be down-regulated in the human lineage. We found that all three categories tend to have lower expression in humans compared to chimpanzee L2/3 IT neurons (Supplemental Figure 42). Overall, this suggests that the down-regulation of ASD-linked genes we observed may be part of a larger trend extending to other genes with similar properties as ASD-linked genes, consistent with previous work on human-specific synaptic gene expression^5^.

Although we cannot rule out any possibility of relaxed constraint at some point in the past, these results favor a model in which polygenic positive selection acted to decrease expression of ASD-linked genes in some types of human neocortical neurons, including L2/3 IT neurons (Figure 6B). As loss of function underlies increased probability of ASD diagnosis for the vast majority of these genes^75^, this suggests that down-regulation of ASD-linked genes may have increased ASD prevalence by bringing humans closer to a hypothetical “ASD expression threshold” below which ASD characteristics manifest. As an example, *DLG4*, which encodes the key synaptic protein PSD-95 and for which loss of one copy causes ASD^76^, has 2.5-fold lower expression in humans compared to chimpanzees (Figure 6H). Consistent with this, it also has 2.5-fold lower protein abundance in the postsynaptic density in humans compared to rhesus macaques, and 3.4-fold lower protein abundance in humans compared to mice^77^ (human vs. rhesus t-test p = 0.0028, human vs. mouse t-test p = 0.00014, Supplemental Figure 43). While this human-specific down-regulation (Supplemental Table 5) that led to the current human baseline expression level of *DLG4* is not sufficient to cause ASD, further down-regulation via loss of a single copy may push humans below the ASD expression threshold whereas loss of a single copy in chimpanzees would maintain expression above this threshold (Figure 6H).

Although these genes are linked to ASD primarily due to their monogenic effects, the majority of ASD cases are thought to be caused by many small genetic and environmental perturbations collectively pushing individuals past some threshold^78^. We propose that the down-regulation of ASD-linked genes in humans increased the likelihood of ASD in the human lineage such that small perturbations on a developmental timescale are sufficient to cause ASD characteristics in humans but not chimpanzees (Figure 6I).

### Down-regulation of schizophrenia-linked genes in humans

Having observed a consistent pattern of human-specific down-regulation for ASD-linked genes, we then tested whether genes linked to schizophrenia (SCZ)^79^, another human-specific neuropsychiatric disorder, show a similar bias. We found an 8-fold enrichment for human down-regulation of SCZ-linked genes in DLPFC L2/3 IT neurons (Supplemental Figure 44A-B).

Although this is even stronger than the ASD bias, it only reaches an FDR < 0.05 in three MTG subclasses, such as Lamp5 and Pax6 inhibitory neurons, due to much lower statistical power (31 SCZ-linked genes vs. 233 high-confidence ASD-linked). Consistent with the known genetic overlap between ASD and SCZ, six of the SCZ-linked genes are also implicated in ASD, making it difficult to disentangle the signal from ASD and SCZ. Furthermore, although there are very few SCZ-linked genes with significant ASE in the hybrid cortical organoid data, among all SCZ-linked genes regardless of significance there is some bias toward human down-regulation, only reaching significance at day 150 (1.5 fold-enrichment, binomial test p = 0.28 at day 100; 2.6 fold-enrichment, binomial test p = 0.025 at day 150, Supplemental Figure 44C). We interpret these results as preliminary evidence that SCZ-linked genes may have also been subject to selection for down-regulation in the human lineage, though further work will be required to confirm this.

## Discussion

Building on an analogy between genes and cell types, we have identified a general principle underlying the rate of evolution of different neuronal types in the mammalian neocortex. We found a strong negative correlation between the abundance of each neuronal cell type and the rate at which its gene expression levels diverge across six mammalian species and three independent datasets^5,7,8^. Interestingly, this correlation remained very strong when collectively analyzing inhibitory and excitatory neurons, despite their very different developmental origins and functions^80,81^.

Based on this initial discovery, we found that L2/3 IT neurons evolved unexpectedly quickly in the human lineage compared to other apes. This accelerated evolution included the disproportionate down-regulation of genes associated with autism spectrum disorder and schizophrenia, two neurological disorders closely linked to L2/3 IT neurons that are common in humans but rare in other apes. Finally, we found that this down-regulation, present both in adult neurons and in organoid models of the developing brain, was likely due to polygenic positive selection on *cis*-regulation. These results differ from, but do not contradict, previous findings that a group of synapse genes show human-specific up-regulation during early development that is disrupted in people with ASD^57^. Overall, our analysis suggests that natural selection on gene expression may have increased the prevalence of ASD, and perhaps also SCZ, in humans (Fig 6H).

Although it has been widely hypothesized that natural selection for human-specific traits has increased human disease risk^46,47,82–84^, unambiguous evidence for this has been lacking. While there is strong evidence linking natural selection on within-human genetic variation to disease risk (e.g. sickle cell disease^85^), it has proven far more challenging to find similar examples involving genetic variants shared by all humans. There are human-chimpanzee differences that have been linked to interspecies differences in disease risk (e.g. human-specific pseudogenization of the *CMAH* gene, which is thought to have shaped human susceptibility to infectious diseases^84,86,87^), but there is no evidence for positive selection on these interspecies genetic differences. In addition, while there are many examples of positive selection on human-chimpanzee differences^67,68,73,88–90^, these changes have no clear link to the likelihood of diseases or disorders in humans. Finally, although the enrichment for ASD-linked variants within HARs^54,55^ is suggestive of a role for human-chimpanzee differences in HARs (many of which are thought to be positively selected^56^) in increasing the likelihood of ASD in humans, a connection between those human-chimpanzee differences and ASD has not been established. Overall, our findings provide the strongest evidence to date supporting the long-standing hypothesis that natural selection for human-specific traits has increased the likelihood of certain disorders.

Although our results strongly suggest natural selection for down-regulation of ASD-linked genes, the reason why this conferred fitness benefits to our ancestors remains an open question. Answering this question is difficult in part because we do not know what human-specific features of cognition, brain anatomy, and neuronal wiring gave our ancestors a fitness advantage, but we can speculate about two general classes of evolutionary scenarios. First, down-regulation of ASD-linked genes may have led to uniquely human phenotypes. For example, haploinsufficiency of many ASD-linked genes is associated with developmental delay^47^, so their down-regulation could have contributed to the slower postnatal brain development in humans compared to chimpanzees. Alternatively, capacity for speech production and comprehension are unique to or greatly expanded in humans and often impacted in ASD and SCZ^53,91^. If down-regulation of ASD-linked genes conferred a fitness advantage by slowing postnatal brain development or increasing the capacity for language, that could result in the signal of positive selection we observed.

On the other hand, the down-regulation we observed may have been compensatory and reduced the negative effects of some other human-specific trait or traits. For example, the ratio of excitatory and inhibitory synapses on pyramidal neurons is fairly constant between humans and rodents despite massive differences in brain and neuron size^92^. In addition, excitatory-inhibitory imbalance is a leading hypothesis for the circuit basis of ASD^93^. If human brain expansion, changes in metabolism, or any other factor shifted this balance away from the fitness optimum, down-regulation of ASD-linked genes could potentially compensate. Overall, more work is needed to understand how natural selection acting on the expression of ASD-linked genes in the human lineage may have shaped human phenotypes.

Our results come with important caveats. As with most correlations, causality is not implied. Our initial hypothesis was that cell type proportions may affect evolutionary rates via more severe fitness effects of expression changes in more abundant cell types, leading to greater evolutionary constraint than in rare cell types (Fig 1A). While this is a plausible explanation for our results, there also may be unknown correlates of cell type proportion that are causal. We leave explicit testing of this model to future work.

Along with establishing a mechanism underlying these correlations, another exciting future direction will be to explore this phenomenon in other tissues and brain regions. While cross-species atlases from other brain regions exist, they generally lack a sufficient number of cells profiled^94,95^ or fail to meet our inclusion criteria in other ways (see Methods). However, this will become increasingly feasible as additional large cross-species snRNA-seq studies are published. An especially interesting question will be whether rare but vital neuron cell types (e.g. serotonergic or dopaminergic neurons^96,97^) follow the same pattern we have observed for neocortical neurons; this will help distinguish between cell type abundance vs. importance as the driving factor underlying the relationship we have observed. It will also be interesting to explore what factors are associated with the rate of cell type-specific gene expression divergence in contexts that lack stable cell type proportions (e.g. during development or in the immune system).

Considering that many ASD-linked genes are extremely sensitive to perturbations in their expression, our findings raise the important question of how significant reductions in the expression of so many dosage-sensitive genes were tolerated in the human lineage. As haploinsufficiency of many of these genes has severe fitness consequences in both humans and mice^47^, it is unlikely that these changes occurred through single mutations of large effect. In addition, our analysis of allele-specific expression suggests that *cis*-regulatory changes underlie many of the gene expression changes we observe. Therefore, we favor a model in which many *cis*-acting mutations of small effect fixed over time, eventually leading to the large-scale down-regulation of ASD-linked genes in the human lineage. It will be interesting to use deep learning predictions of variant effects combined with experimental validation to identify the genetic differences underlying changes in the expression of ASD-linked genes in the human lineage.

It is also possible that the down-regulation of many ASD-linked genes is less deleterious than the down-regulation of a single gene. As an analogy, whole-genome duplications can be well-tolerated in vertebrates, even though duplication of some individual genes—including many of those linked to ASD—can be far more deleterious. An intuitive explanation for this counter-intuitive observation is that relative expression levels, or stoichiometry, could impact fitness even more than absolute expression levels^98^. Under this model, the key idea is that the down-regulation of many ASD-linked genes would have less impact on their relative levels than a change in the expression of a single gene. Excitingly, CRISPR-based methods to precisely manipulate the expression levels of many genes at once may soon allow us to more directly test this hypothesis. Overall, it will be important to develop a deeper understanding of how cell types and genes implicated in ASD and SCZ have evolved in the human lineage as this will improve our understanding of uniquely human traits and neuropsychiatric disorders.

## Methods

### Quantifying cell type-specific gene expression divergence between species

We analyzed three main datasets in this study, which we refer to by the cortical area sampled (MTG, DLPFC, M1). These were the only studies meeting both of our inclusion criteria: multiple species profiled in the same study using the same snRNA-seq protocols for each species within a study, and at least 10 orthologous cell types having 250 or more cells per species. The following are examples of studies that did not meet these inclusion criteria:

- A multi-species study of the retina used different protocols for different species and not all species were sampled as part of the same original study. For example, different antibodies were used to enrich for subpopulations of cells in different species and some species did not have a sufficient number of cells profiled without enrichment to accurately estimate cell type proportions^99^.
- A multi-species study of substantia nigra dopaminergic neurons did not have a sufficient number of cells profiled per species^94^.
- A multi-species study of the lateral geniculate nucleus did not profile enough cells per species and their dissection scheme was incompatible with estimating neuronal cell type proportions^95^.

All statistical tests and analyses were performed in python using scipy v1.10.1^100^ except for the DESeq2 analysis. For the M1 and MTG data, we converted from RDS files to h5 files using Seurat and Seurat Disk^101^. We conducted all analyses within each dataset to avoid batch effects from comparing across datasets. We used the cell type annotations and counts matrices directly from the study that first reported the dataset in conjunction with scanpy v1.7.2^102^. The procedure outlined below was performed 100 times independently on each dataset unless otherwise noted. To quantify cell type-specific expression divergence without confounding with cell type proportion, we first down-sampled the number of cells in each cell type so that it was equal across all cell types and species. We down-sampled without replacement to 250 cells at the subclass level and 50 cells at the subtype level for the main analysis presented in the text. Only subclasses and subtypes with at least this many cells were included in downstream analysis.

We then restricted to 5-way one-to-one protein-coding non-mitochondrial orthologs (downloaded from ensembl biomart for hg38)^103^ between human, chimpanzee, gorilla, rhesus macaque, and marmoset for the MTG and DLPFC data and 3-way one-to-one orthologs for human, marmoset, and mouse for the M1 dataset. We then summed expression across all cells within a cell type to create a pseudobulked expression profile for that cell type.

For each possible pairwise comparison between species, we down-sampled the total counts in each cell type so that it was equal across all cell types for both species in the comparison. We then computed counts per million (CPM) in each cell type. After computing CPM, we filtered out genes with (1) fewer than 25 counts in both species or (2) fewer than 1 CPM in both species per cell type. As a result, if a gene passed the filtering criteria in one cell type but not another it would be included only for the cell type in which it passed the filtering criteria. We then computed the log_2_(CPM) and used the Spearman correlation distance to measure the gene expression divergence between species in each cell type.

Notably, this process involved several analysis decisions that could affect our results. To test how robust our results were to these choices, we tested all combinations of the following:

1. Down-sampling to 50, 100, 250, or 500 cells.
2. Filtering genes with fewer than 5, 10, 25, or 50 counts.
3. Filtering genes with fewer than 1 or 5 CPM.
4. Using log_2_(CPM) or not log transforming.
5. Using the Spearman correlation distance, Pearson correlation distance, Euclidean distance, or L1 distance metrics.

In general, our results were robust to any combination of these parameters (Supplemental Tables). When stratifying, we only used a subset of these combinations due to the greater number of computations required.

### Computing cell type proportions and correlation with gene expression divergence

All three datasets were generated with single-nucleus RNA-sequencing (snRNA-seq) and so likely accurately represent the true proportion of neuronal cell types in the neocortex^104^. To compute cell type proportions, we restricted to neuronal cells with greater than or equal to the number of cells we down-sampled to. We then computed cell type proportion separately for each species by dividing the number of cells of each type by the total number of cells profiled. For each interspecies comparison, we averaged the cell type proportion across both species. We then computed the Spearman correlation between the averaged cell type proportions and cell type-specific gene expression divergence computed as described above. As we did this across 100 independent down-samplings (numbered 1 to 100), we reported the median Spearman’s rho and p-value throughout the text and figures. If there was an individual down-sampling iteration that had the median Spearman’s rho and p-value, we made the scatterplots shown in Figures 1-4 using the first such iteration. If no iteration had the median rho and p-value, we showed the iteration closest to the median with the greatest number of iterations that had that rho and p-value. For example, if 22 iterations resulted in rho = -0.5 and 19 iterations resulted in rho = -0.6, both of which were closest to the median of -0.55, then an iteration with - 0.5 would be shown. If there was still a tie after this process, we showed the iteration with the lowest number. Because the Spearman correlation is a nonparametric rank-based test, it is unaffected by any rank-preserving transformation of the data; therefore our choice to show scatter plots with log-transformed cell type proportions was for visualization only and had no effect on the results.

To estimate divergence along the human branch, we used the formula:

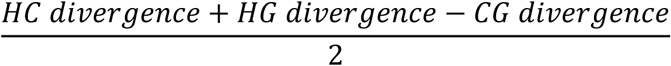

Here, HC stands for human-chimp, HG stands for human-gorilla, and CG stands for chimp-gorilla.

Similarly, to estimate divergence along the chimp branch, we used the formula:

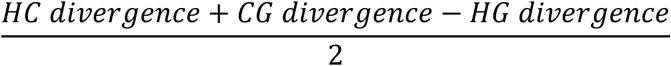

### Stratifying by expression level, cell type-specificity of expression, and constraint on expression

To stratify by expression level, we ranked genes by the average CPM between the two species being compared for each cell type separately. We then assigned the top third of genes with the highest expression to the highly expressed bin, the next third to the moderately expressed bin, and the remaining third to the lowly expressed bin. Whenever we stratified by expression level or another metric, we used the Euclidean distance to measure gene expression divergence because the limited dynamic range of expression for the moderately and lowly expressed bins led to unrealistically high correlation distances. Similarly, we ranked genes by Tau^62^, a measure of how cell type-specifically a gene is expressed, and split those genes into three bins. We computed Tau separately for both species across all subclasses or subtypes with a sufficient number of cells and then computed the average value for each gene. For constraint on expression, we considered all genes with heterozygous fitness effect^61^ s_het_ > 0.1 to be highly constrained, genes with s_het_ between 0.1 and 0.01 as moderately constrained, and the remaining genes with s_het_ < 0.01 to be lowly constrained. Because there was a different number of genes in each bin in this case, we down-sampled genes to reach an equal number in each bin.

When controlling for expression level and stratifying by Tau, we compared the high bin with the moderate and low bins separately. To control for expression, we first computed the log_2_ fold-change between all genes in the high bin and all genes in the moderate or low bin and restricted to pairs of genes with absolute log_2_ fold-change less than 0.05. We then split this list of gene pairs into those with a negative log_2_ fold-change, positive log_2_ fold-change, and zero log_2_ fold-change, shuffled the list, and removed duplicate genes. We kept all gene pairs with a log_2_ fold-change of zero and down-sampled the list of gene pairs with positive or negative log_2_ fold-change so that there were an equal number in each category. This resulted in a final set of genes in the high bin with matched expression to genes in the moderate or low bin which we used to compute cell type-specific gene expression divergence. When controlling for Tau, we applied the same strategy but required an absolute log_2_ fold-change less than 0.01.

### Comparing interindividual variability in gene expression and cell type proportion

To measure the within-human interindividual variation in cell type-specific gene expression, we used a uniformly processed dataset from the DLPFC^63^. We restricted to control samples from individuals of European ancestry with an age of death greater than or equal to 25. We selected thirteen neuronal subclasses for which the majority of individuals had greater than 50 nuclei profiled for further analysis and restricted to samples with greater than or equal to 50 nuclei for all thirteen subclasses. After this filtering process, 25 samples remained. Next, we down-sampled to 50 nuclei from each subclass in each dataset and computed pseudobulked counts. We then down-sampled counts so that there was an equal number of total counts across all subclasses for each individual. For each subclass, we removed genes with average counts across all individuals less than 25 and computed CPM. We then computed the Spearman correlation distance between each sample and the mean expression profile across all samples and took the mean of those 25 correlation distances as our measure of cell type-specific gene expression variation within humans. We performed this procedure across 100 independent down-samplings. To estimate cell type proportions, we computed the cell type proportions for the thirteen subclasses and averaged them together. We then computed the Spearman correlation between the subclass-specific interindividual variation and the cell type proportions across the 100 down-samplings. We report the median Spearman’s rho and p-value across the 100 down-samplings and show the first down-sampling with the median Spearman’s rho and p-value in Figure 5D.

### Analysis of ASD- and SCZ-linked genes in snRNA-seq data

We used the SFARI gene database of ASD-linked genes and considered any genes with a score of 1 to be “high-confidence” (233 total) and all genes regardless of score to be all ASD-linked genes (1176 genes)^66^. As we are not aware of a similar resource for SCZ, we used the 31 genes with FDR < 0.1 in a recent rare variant association study for SCZ^79^. Throughout, FDRs were corrected for multiple tests with the Benajmini-Hochberg method. To identify differentially expressed (DE) genes and compute log_2_ fold-changes between species, we ran DESeq2^105^ on the subclass-level pseudobulked counts and used apeglm^106^ to shrink the log_2_ fold-changes. To test for a bias toward lower expression of ASD- and SCZ-linked genes in each cell type, we restricted to genes with FDR < 0.05 in the human-chimpanzee comparison and use the binomial test comparing the number of genes with negative log_2_ fold-change (i.e. higher expression in chimpanzee) to the number of genes with positive log_2_ fold-change. We used the frequency of negative log_2_ fold-changes among all genes with FDR < 0.05 as the background probability in the binomial test. We repeated this for both high-confidence and all ASD-linked genes.

To determine whether the higher expression in chimpanzees relative to human was more likely due to changes on the chimpanzee branch or the human branch, we first filtered to only high-confidence ASD-linked genes that were differentially expressed between chimpanzees and gorillas in L2/3 IT neurons. Genes were assigned as having a significant human-derived or chimpanzee-derived expression change in the MTG dataset by comparison with the human-gorilla and chimpanzee-gorilla log_2_ fold-changes. First, if the absolute human-gorilla and chimpanzee-gorilla log_2_ fold-change were both greater than the absolute human-chimpanzee log_2_ fold-change, that gene was considered ambiguous. After removing ambiguous genes, a gene was considered as having a human-derived expression change if the absolute human-gorilla log_2_ fold-change was greater than the absolute human-chimpanzee log_2_ fold-change and vice versa for chimpanzee-derived. To generate Supplemental table 5, we used strict criteria to call genes as having a human-specific gene expression change in the MTG data, requiring that a gene be differentially expressed (i.e. FDR < 0.05) for each human-NHP comparison with the same direction of differential expression. We then added the SFARI score and whether a gene is considered syndromic and only include genes that are differentially expressed (FDR < 0.05) between human and chimpanzee.

### Analysis of ASD-linked genes in human-chimpanzee hybrid cortical organoid data

We used the previously described dataset from human-chimpanzee cortical organoids, reprocessed as previously described^89^. Briefly, reads were aligned to the human (hg38) and chimpanzee (PanTro6) genomes with STAR and corrected for mapping bias using Hornet^107^. Reads were assigned to the human or chimpanzee allele using a set of high-confidence human-chimp single nucleotide differences and collapsed to counts per gene with ASEr. DESeq2^105^ was used to identify genes with significant ASE with the hybrid line that each sample was from used as a covariate. DESeq2^105^ and apeglm^106^ were used to compute log_2_ fold-changes. For the below analyses, we used the chimpanzee-aligned data, which has a very slight bias toward higher expression from the human allele, to ensure that our analyses were conservative.

To test for a significant bias toward down or up-regulation from the human allele for ASD- or SCZ-linked genes, we restricted to genes with FDR < 0.05 in the cortical organoid data and intersected those genes with the list of ASD- or SCZ-linked genes. We then used the binomial test comparing the number of genes with negative log_2_ fold-change (i.e. higher expression in chimpanzee) to the number of genes with positive log_2_ fold-change. We used the frequency of negative log_2_ fold-changes among all genes with FDR < 0.05 as the background probability in the binomial test. We repeated this for both high-confidence and all ASD-linked genes. To investigate whether these *cis*-regulatory changes likely occurred in the human or chimpanzee lineage, we used the assignments as human- or chimpanzee-derived from L2/3 IT neurons in the MTG dataset described above. For genes that had matching human-chimpanzee log_2_ fold-change sign in both the MTG and cortical organoid datasets, we created a 2×2 table of human/chimp-derived and down/up-regulated from the human allele and applied Fisher’s exact test.

### Analysis of constraint on ASD-linked genes in humans and chimpanzees

We used previously published dN/dS estimates^73^ and restricted only to genes with at least one synonymous and nonsynonymous difference on both the human and chimpanzee branches. We compared dN/dS values for ASD-linked genes with a paired t-test. To compute the number of genetic differences within 5 kilobases of the transcription start site (TSS) for each lineage, we used our previously described set of high-confidence human-chimpanzee single nucleotide genetic differences^89^. Briefly, this was created by identifying all single nucleotide differences between PanTro6 and hg38 and the filtering out sites that were not homozygous for the reference allele in 3 humans and 3 chimpanzees. We then intersected this with a previously described list of human-chimpanzee orthologous TSS expanded by 2.5 kilobases on either side and restricted to only TSS for ASD-linked genes^90^. To correct for the slightly larger number of human-derived sites across all genes, we down-sampled the human-derived variants near the TSS of ASD-linked genes, keeping a fraction of sites equal to the total number of chimp-derived genetic differences divided by the total number of human-derived genetic differences. We then used a paired t-test to compare the two distributions.

To compare the within-species variance for humans and chimpanzees in expression of ASD-linked genes, we computed the variance in pseudobulked CPM from L2/3 IT neurons across individuals in the DLPFC and MTG separately. As the mean expression level and batch effects can have a major impact on expression variance, we normalized the variance to the variance of the 100 genes with closest mean expression to each ASD-linked gene. To do this, we computed the fraction of those 100 genes with smaller variance than the focal ASD-linked gene in each species and dataset separately. We then compared the values in human and chimpanzee with a paired t-test.

### Comparing different phenotypes and gene categories to ASD-linked genes

To compare down-regulation of high-confidence ASD-linked genes to genes associated with other phenotypes, we used the human phenotype ontology (HPO) restricting to phenotypes with at least 100 genes. We tested all these gene sets in addition to the high-confidence ASD-linked genes and computed fold-enrichment as described above for ASD-linked genes. We controlled for gene expression as described in the “Stratifying by expression level, cell type-specificity of expression, and constraint on expression” section, filtering out all gene pairs with an absolute log fold-change greater than 0.1.

To subset ASD-linked genes, we used all genes present in the SynGo database^108^ as our list of synaptic genes, all genes classified as “1 Monomer or homomultimer”, “2 Obligate heteromer”, “3 Low specificity DNA-binding protein” from Lambert et al.^109^ as our list of transcription factors and chromatin remodelers, and all genes with pLI > 0.9 from gnomad version 4.1^74^ as our list of haploinsufficient genes. We intersected these with the set of ASD-linked genes with these lists and removed all ASD-linked genes from those lists to define genes as “ASD-linked and in a category” or “ASD-linked and not in a category” respectively. We also removed genes in a category from the list of ASD-linked genes to define the list of genes that are ASD-linked and, for example, not synaptic. When working with the MTG data, we always subsequently restricted to high-confidence ASD-linked genes. With these categories in hand, we then computed the proportion of genes in each category that are down-regulated. We used the binomtest function from scipy with p set to the proportion of genes in a category not linked to ASD that are down-regulated to test whether ASD-linked genes within a particular category were more down-regulated than genes in the category that are not linked to ASD.

### Analysis of postsynaptic proteomics data

We plotted PSD-95 protein abundances from the supplemental materials of Wang et al^77^. We used the t-test to compare levels between species.

## Acknowledgements

We thank Liqun Luo and other Luo lab members for helpful discussion. We also thank Leslie Magtanong and other members of the Fraser Lab for helpful discussions and feedback on the manuscript. Some subfigures were made with biorender.

## Funding

Funding was provided by NIH R01HG012285 (awarded to HBF). ALS was supported by a fellowship under grant number FA9550-21-F-0003.

## Authors contributions

ALS performed all bioinformatic analysis, visualization, validation, and writing of software with guidance from HBF. ALS and HBF wrote the manuscript and ALS created the figures with input from HBF. HBF provided funding for the study.

## Competing interests

All authors declare no competing interests.

## Data availability

The MTG data are available from https://labshare.cshl.edu/shares/gillislab/resource/Primate_MTG_coexp/Great_Ape_Data/. The metadata for the MTG study are available from https://github.com/AllenInstitute/Great_Ape_MTG/blob/master/data/ (files ending in “for_plots_and_sharing_12_16_21.RDS”). The DLPFC data are available from https://data.nemoarchive.org/biccn/grant/u01_sestan/sestan/transcriptome/sncell/10x_v3/. The M1 data are available from https://data.nemoarchive.org/publication_release/Lein_2020_M1_study_analysis/Transcriptomic s/sncell/10X/. The constraint metric s_het_ was downloaded from the supplemental materials of https://www.biorxiv.org/content/10.1101/2023.05.19.541520v1. The constraint metric pLI was downloaded from https://gnomad.broadinstitute.org/downloads#v4-constraint. The SFARI ASD-linked genes were downloaded from https://gene.sfari.org/. The SCZ-linked genes were downloaded from https://www.nature.com/articles/s41586-022-04556-w. Protein abundance measurements in the post-synaptic density of humans, rhesus macaques, and mice were obtained from the supplemental materials of https://www.nature.com/articles/s41586-023-06542-2. The human population DLPFC single nucleus RNA-seq data used to compute within human cell type-specific gene expression variation were downloaded from https://brainscope.gersteinlab.org/output-sample-annotated-matrix.html. The human-chimpanzee hybrid cortical organoid data were downloaded from https://www.ncbi.nlm.nih.gov/geo/query/acc.cgi?acc=GSE144825. dN/dS estimates for the human and chimp lineages were downloaded from https://doi.org/10.1186/1471-2164-15-599. All code needed to reproduce the analyses described in this study is available at https://github.com/astarr97/Cell_Type_Evolution.

## Materials and correspondence

Correspondence should be addressed to HBF at and ALS at. Requests for materials will be fulfilled by the corresponding authors.

## List of supplementary materials

Supplemental Figures 1-44

Supplemental Tables 1-5

Supplemental Table 1 contains the median Spearman’s rho and p-value for the correlation between cell type divergence and proportion across a variety of parameter combinations.

Supplemental Table 2 contains the median Spearman’s rho and p-value for the correlation between cell type divergence and proportion stratifying by expression level across a variety of parameter combinations.

Supplemental Table 3 contains the median Spearman’s rho and p-value for the correlation between cell type divergence and proportion stratifying by evolutionary constraint across a variety of parameter combinations.

Supplemental Table 4 contains the median Spearman’s rho and p-value for the correlation between cell type divergence and proportion stratifying by cell type-specificity of expression across a variety of parameter combinations.

Supplemental Table 5 contains the log_2_ fold-changes and FDR-adjusted p-values for all human/NHP comparisons in the MTG dataset, whether those changes were classified as human-derived or human-specific according to the criteria outlined in the Methods, and the SFARI gene scores as well as whether a gene is considered syndromic in SFARI.

## Notes

### Competing Interest Statement

The authors have declared no competing interest.

### Summary of Updates

Added analysis of whether subsets of ASD-linked genes show a greater bias, comparing downregulation of ASD-linked genes to downregulation of other sets of genes linked to different phenotypes, clarifications throughout the text and figures.

