## Supplemental Figures for "A general principle of neuronal evolution reveals a human-accelerated neuron type potentially underlying the high prevalence of autism in humans"

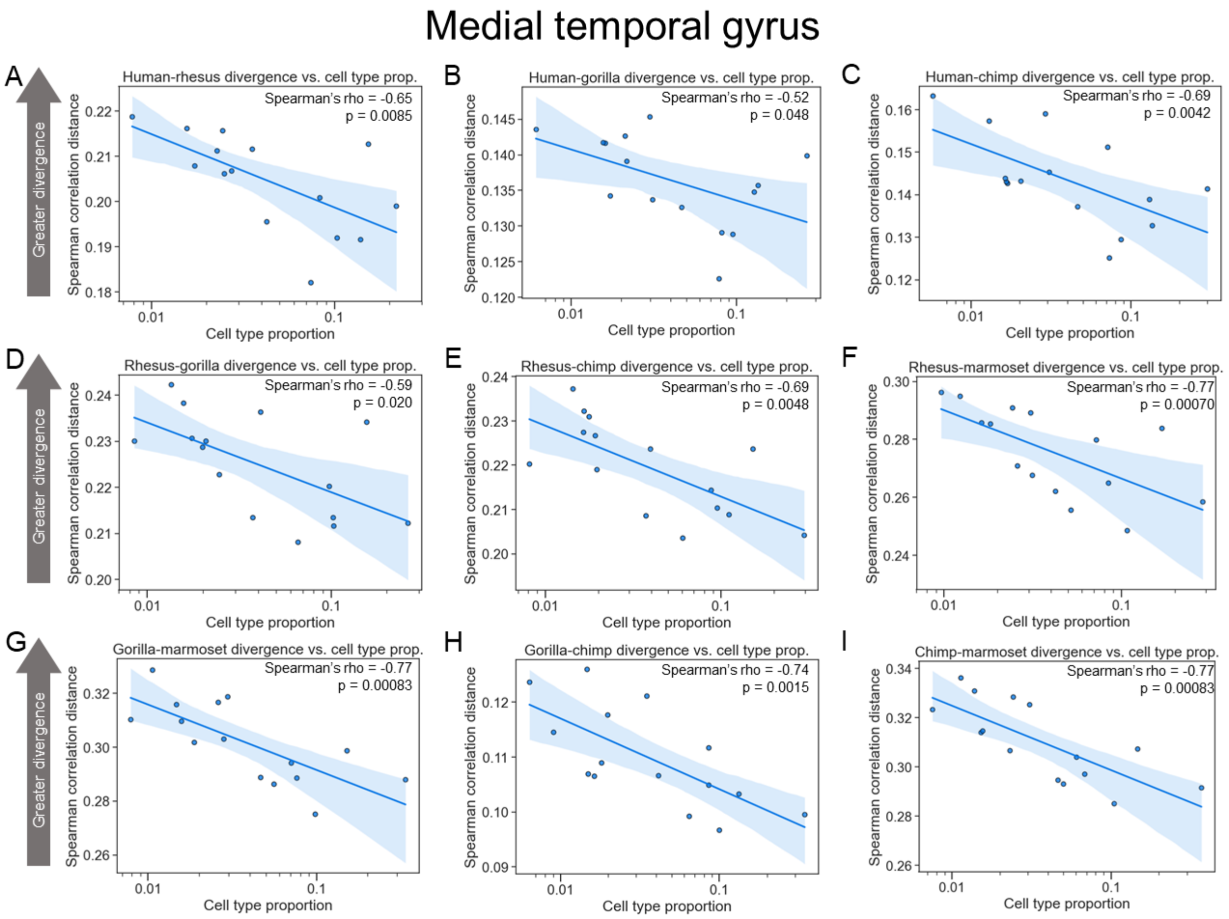


**Supplemental Figure 1: More common neuronal subclasses evolve more slowly than rarer subclasses in the MTG. A)** Plot showing the correlation between neuronal subclass proportion ($log_{10}$ scale on the x-axis) and subclass-specific divergence between human and rhesus in the MTG for all neurons. **B)** Same as in (A) but for human and gorilla. **C)** Same as in (A) but for human and chimp. **D)** Same as in (A) but for rhesus and gorilla. **E)** Same as in (A) but for rhesus and chimp. **F)** Same as in (A) but for rhesus and marmoset. **G)** Same as in (A) but for gorilla and marmoset. **H)** Same as in (A) but for gorilla and chimp. **I)** Same as in (A) but for chimp and marmoset.

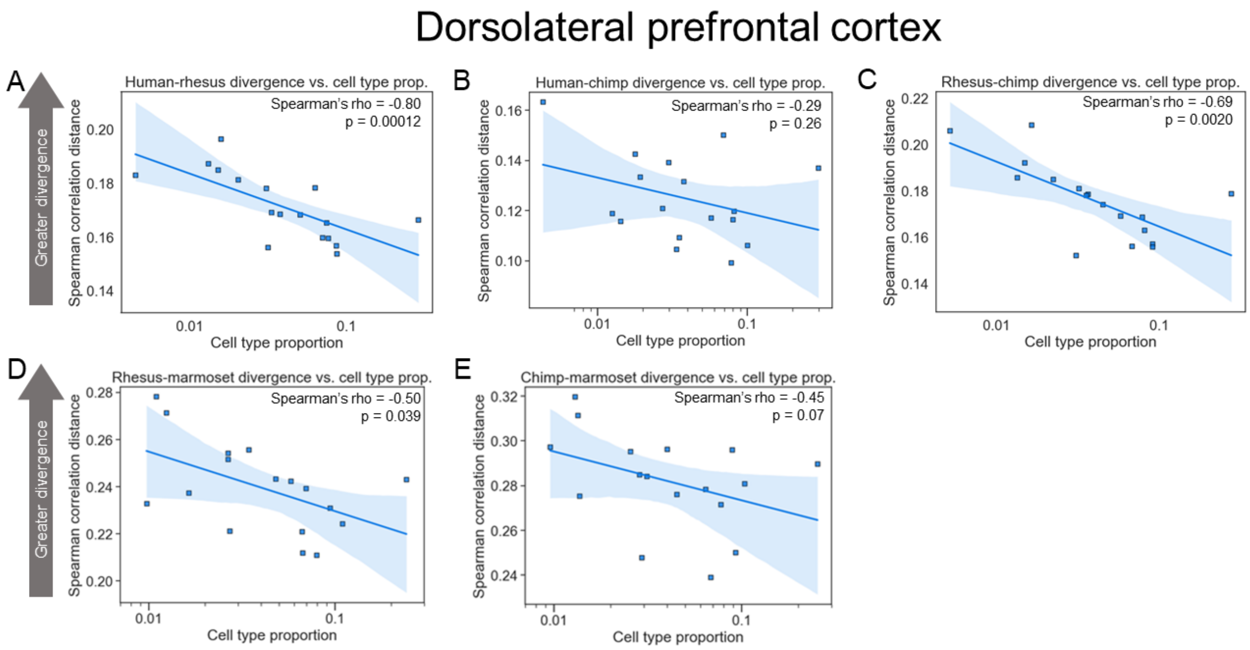


**Supplemental Figure 2: More common neuronal subclasses evolve more slowly than rarer subclasses in the DLPFC. A)** Plot showing the correlation between neuronal subclass proportion ($\log_{10}$ scale on the x-axis) and subclass-specific divergence between human and rhesus in the DLPFC for all neurons. **B)** Same as in (A) but for human and chimp. **C)** Same as in (A) but for rhesus and chimp. **D)** Same as in (A) but for rhesus and marmoset. **E)** Same as in (A) but for chimp and marmoset.

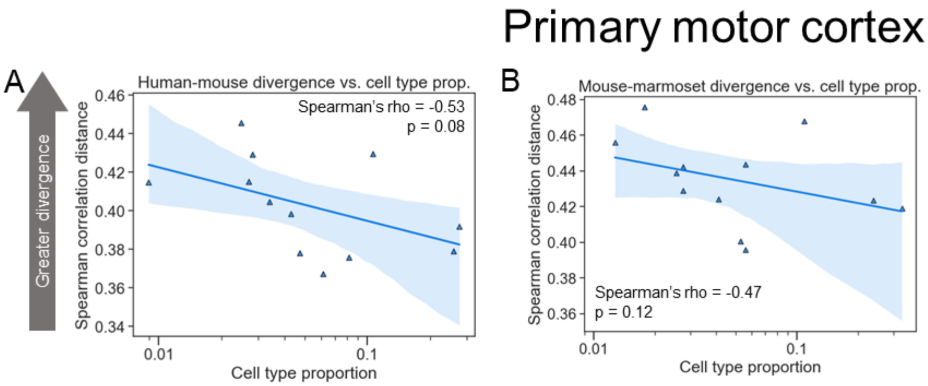


**Supplemental Figure 3: More common neuronal subclasses evolve more slowly than rarer subclasses in M1. A)** Plot showing the correlation between neuronal subclass proportion ($log_{10}$ scale on the x-axis) and subclass-specific divergence between human and mouse in M1 for all neurons. **B)** Same as in (A) but for mouse and marmoset.

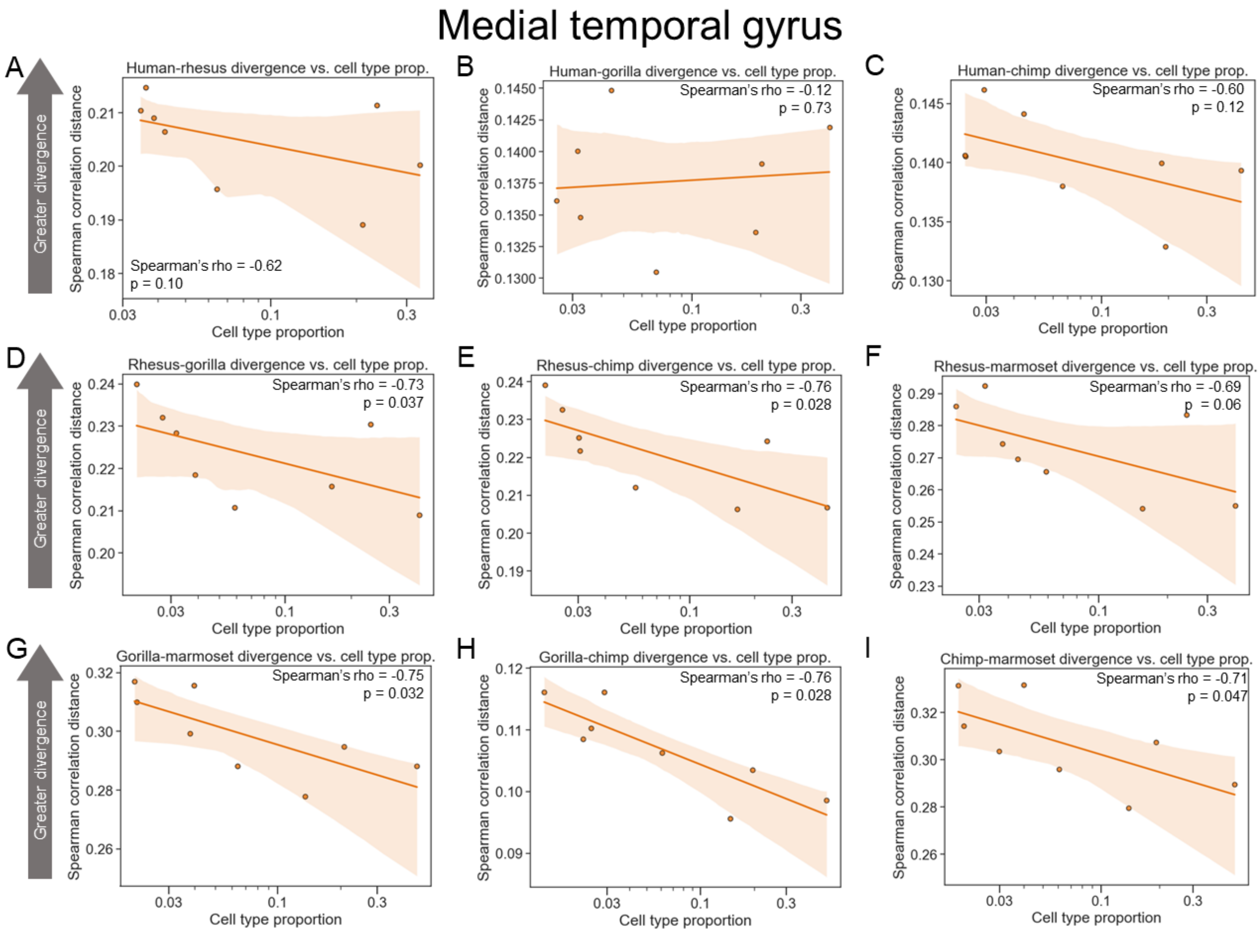


**Supplemental Figure 4: More common excitatory neuronal subclasses evolve more slowly than rarer subclasses in the MTG. A)** Plot showing the correlation between neuronal subclass proportion ($\log_{10}$ scale on the x-axis) and subclass-specific divergence between human and rhesus in the MTG for excitatory neurons. **B)** Same as in (A) but for human and gorilla. **C)** Same as in (A) but for human and chimp. **D)** Same as in (A) but for rhesus and gorilla. **E)** Same as in (A) but for rhesus and chimp. **F)** Same as in (A) but for rhesus and marmoset. **G)** Same as in (A) but for gorilla and marmoset. **H)** Same as in (A) but for gorilla and chimp. **I)** Same as in (A) but for chimp and marmoset.

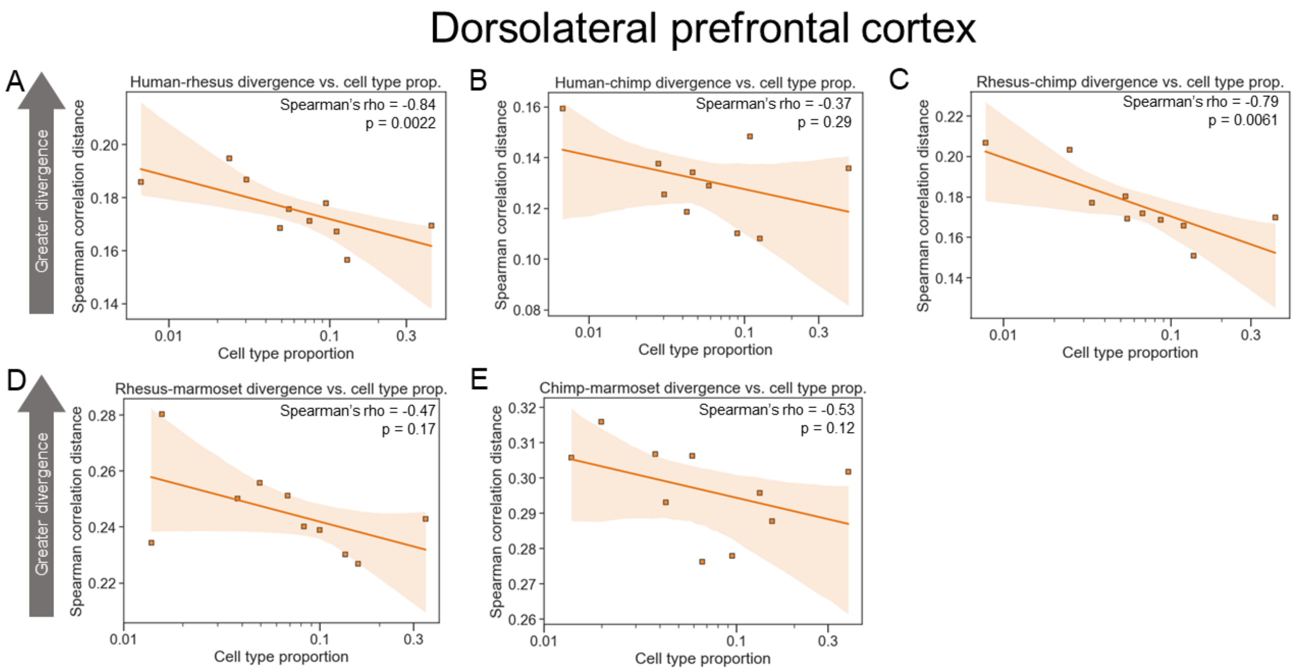


**Supplemental Figure 5: More common excitatory neuronal subclasses evolve more slowly than rarer subclasses in the DLPFC. A)** Plot showing the correlation between neuronal subclass proportion ($log_{10}$ scale on the x-axis) and subclass-specific divergence between human and rhesus in the DLPFC for excitatory neurons. **B)** Same as in (A) but for human and chimp. **C)** Same as in (A) but for rhesus and chimp. **D)** Same as in (A) but for rhesus and marmoset. **E)** Same as in (A) but for chimp and marmoset.

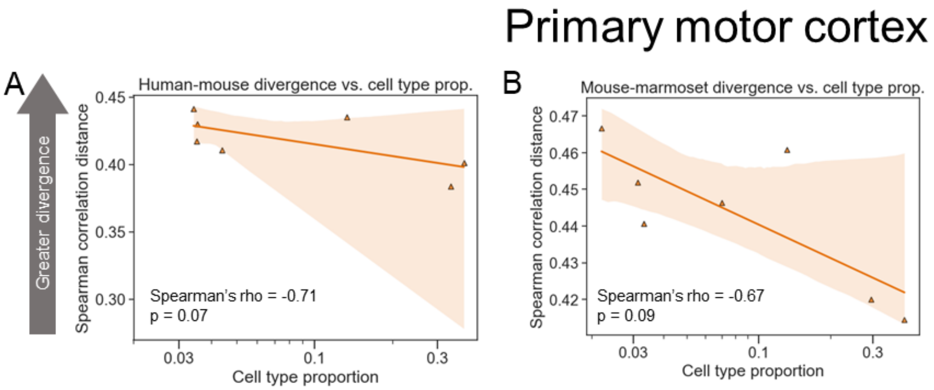


**Supplemental Figure 6: More common excitatory neuronal subclasses evolve more slowly than rarer subclasses in M1. A)** Plot showing the correlation between neuronal subclass proportion ($\log_{10}$ scale on the x-axis) and subclass-specific divergence between human and mouse in M1 for excitatory neurons. **B)** Same as in (A) but for mouse and marmoset.

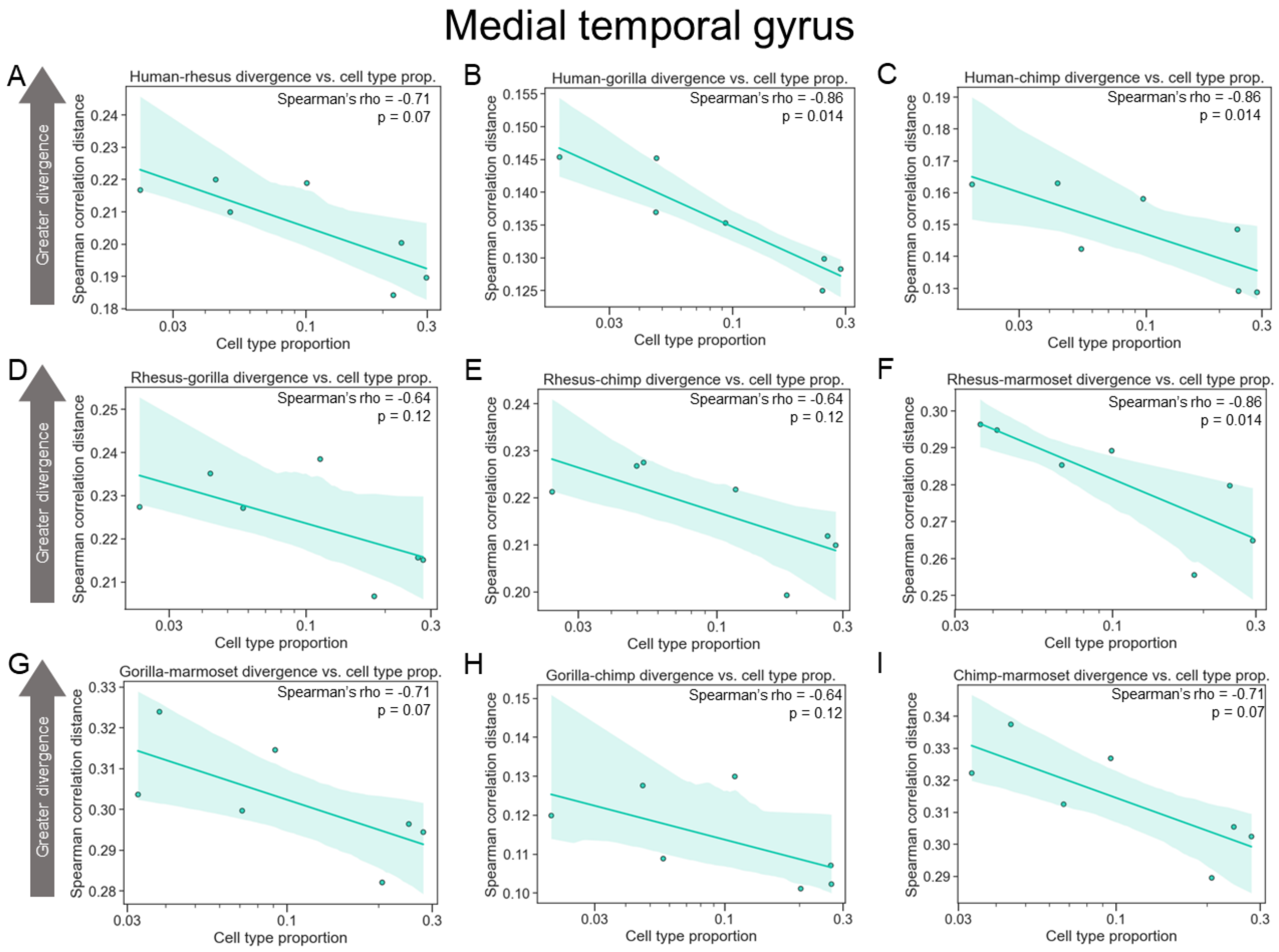


**Supplemental Figure 7: More common inhibitory neuronal subclasses evolve more slowly than rarer subclasses in the MTG. A)** Plot showing the correlation between neuronal subclass proportion ($log_{10}$ scale on the x-axis) and subclass-specific divergence between human and rhesus in the MTG for inhibitory neurons. **B)** Same as in (A) but for human and gorilla. **C)** Same as in (A) but for human and chimp. **D)** Same as in (A) but for rhesus and gorilla. **E)** Same as in (A) but for rhesus and chimp. **F)** Same as in (A) but for rhesus and marmoset. **G)** Same as in (A) but for gorilla and marmoset. **H)** Same as in (A) but for gorilla and chimp. **I)** Same as in (A) but for chimp and marmoset.

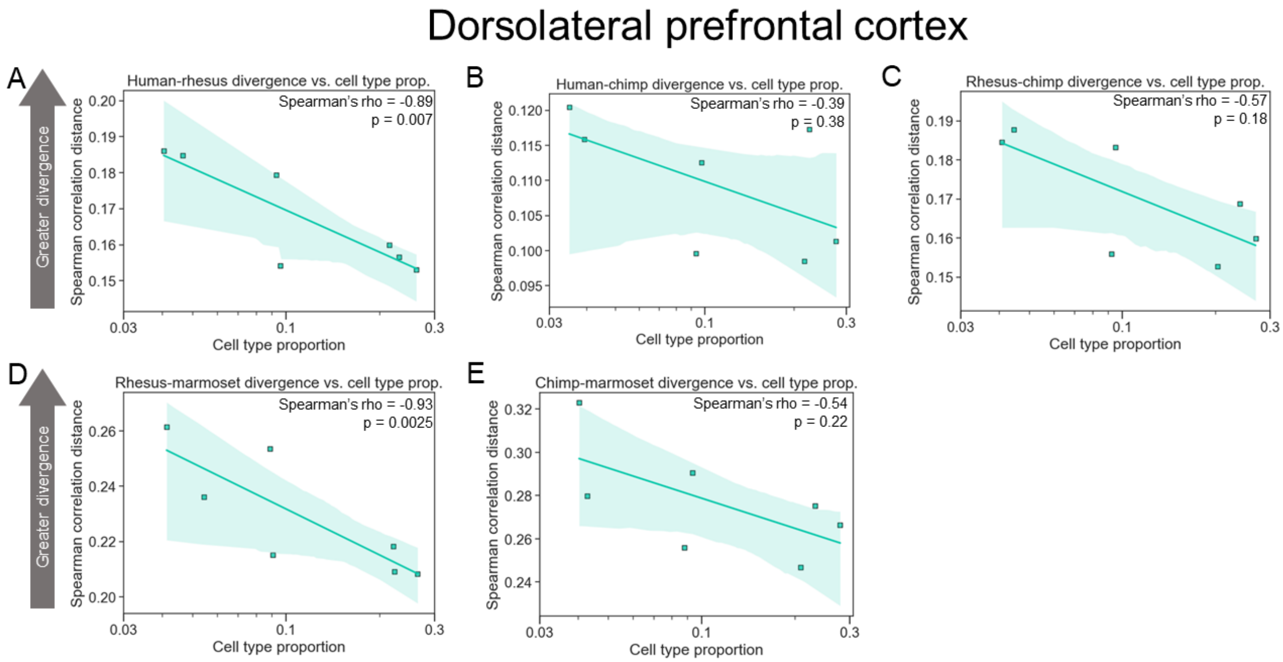


**Supplemental Figure 8: More common inhibitory neuronal subclasses evolve more slowly than rarer subclasses in the DLPFC. A)** Plot showing the correlation between neuronal subclass proportion ($\log_{10}$ scale on the x-axis) and subclass-specific divergence between human and rhesus in the DLPFC for inhibitory neurons. **B)** Same as in (A) but for human and chimp. **C)** Same as in (A) but for rhesus and chimp. **D)** Same as in (A) but for rhesus and marmoset. **E)** Same as in (A) but for chimp and marmoset.

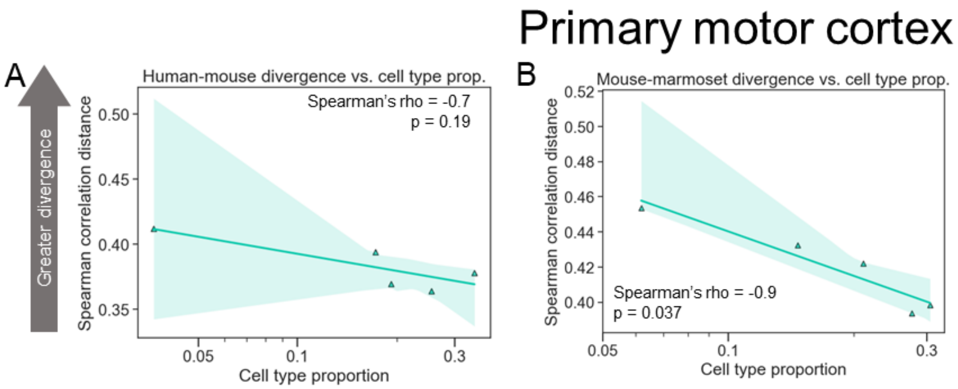


**Supplemental Figure 9: More common inhibitory neuronal subclasses evolve more slowly than rarer subclasses in M1. A)** Plot showing the correlation between neuronal subclass proportion ($log_{10}$ scale on the x-axis) and subclass-specific divergence between human and mouse in M1 for inhibitory neurons. **B)** Same as in (A) but for mouse and marmoset.

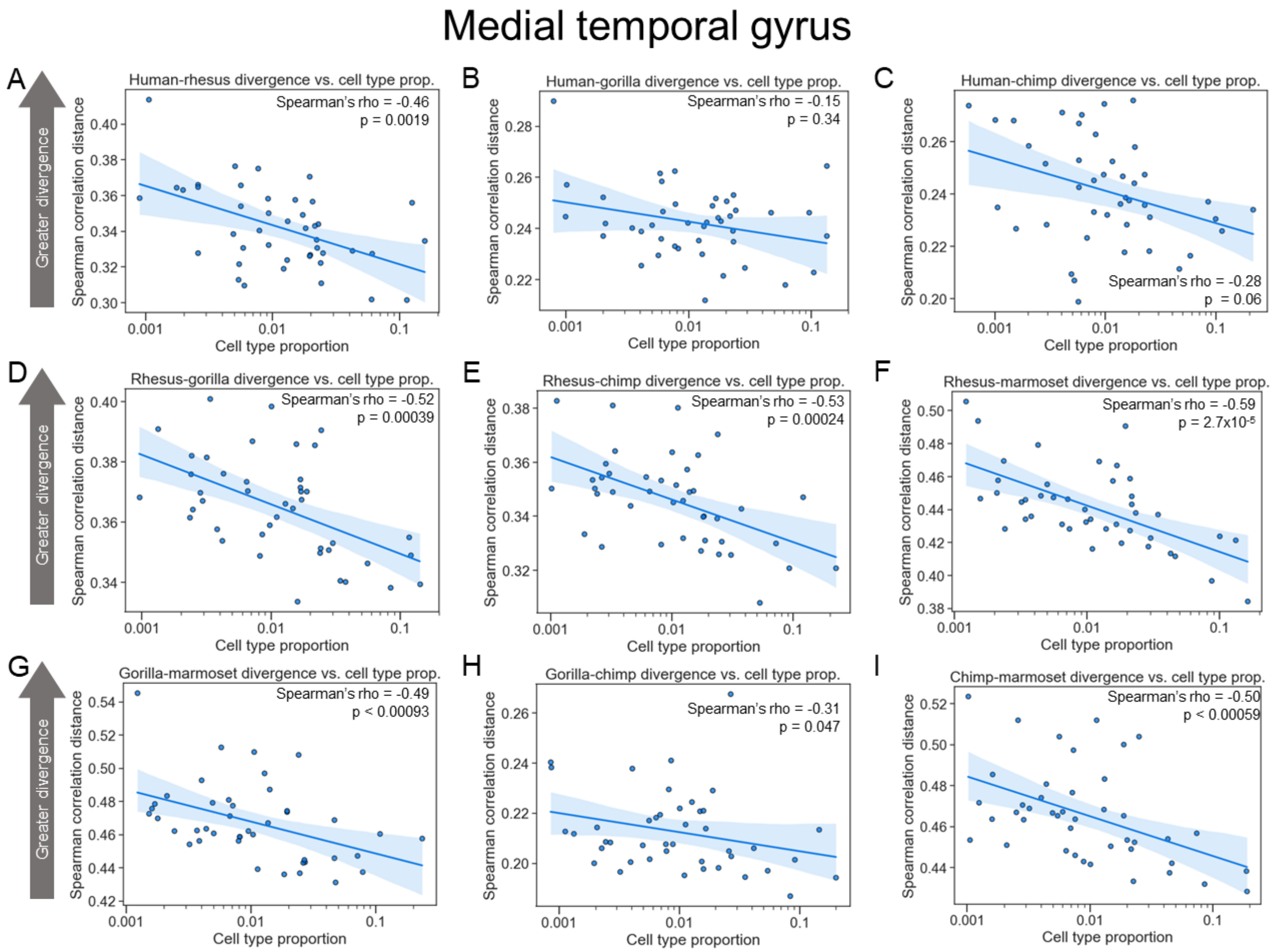


**Supplemental Figure 10: More common neuronal subtypes evolve more slowly than rarer subtypes in the MTG. A)** Plot showing the correlation between neuronal subtype proportion ($log_{10}$ scale on the x-axis) and subtype-specific divergence between human and rhesus in the MTG for all neurons. **B)** Same as in (A) but for human and gorilla. **C)** Same as in (A) but for human and chimp. **D)** Same as in (A) but for rhesus and gorilla. **E)** Same as in (A) but for rhesus and chimp. **F)** Same as in (A) but for rhesus and marmoset. **G)** Same as in (A) but for gorilla and marmoset. **H)** Same as in (A) but for gorilla and chimp. **I)** Same as in (A) but for chimp and marmoset.

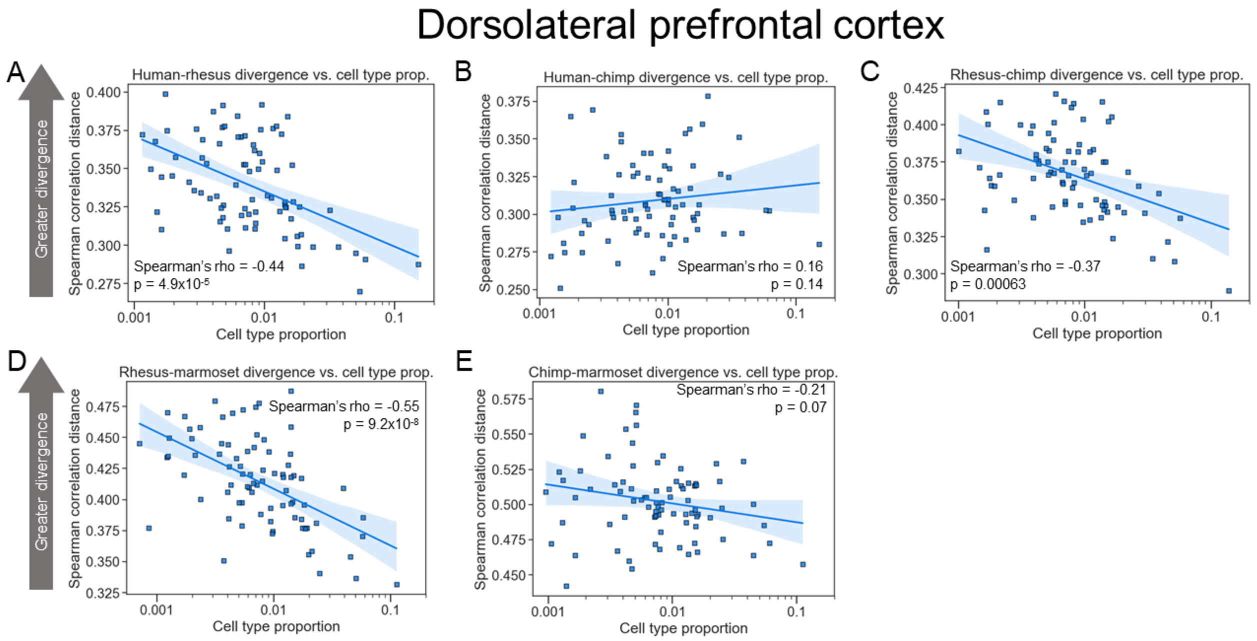


**Supplemental Figure 11: More common neuronal subtypes evolve more slowly than rarer subtypes in the DLPFC. A)** Plot showing the correlation between neuronal subtype proportion ($\log_{10}$ scale on the x-axis) and subtype-specific divergence between human and rhesus in the DLPFC for all neurons. **B)** Same as in (A) but for human and chimp. **C)** Same as in (A) but for rhesus and chimp. **D)** Same as in (A) but for rhesus and marmoset. **E)** Same as in (A) but for chimp and marmoset.

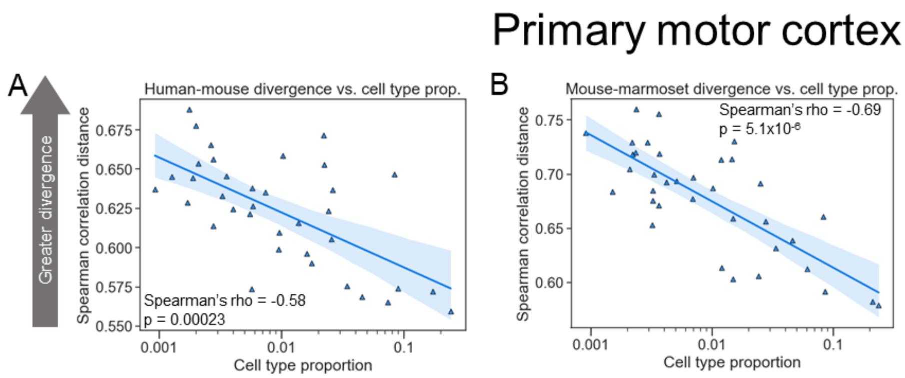


**Supplemental Figure 12: More common neuronal subtypes evolve more slowly than rarer subtypes in M1. A)** Plot showing the correlation between neuronal subtype proportion ($\log_{10}$ scale on the x-axis) and subtype-specific divergence between human and mouse in M1 for all neurons. **B)** Same as in (A) but for mouse and marmoset.

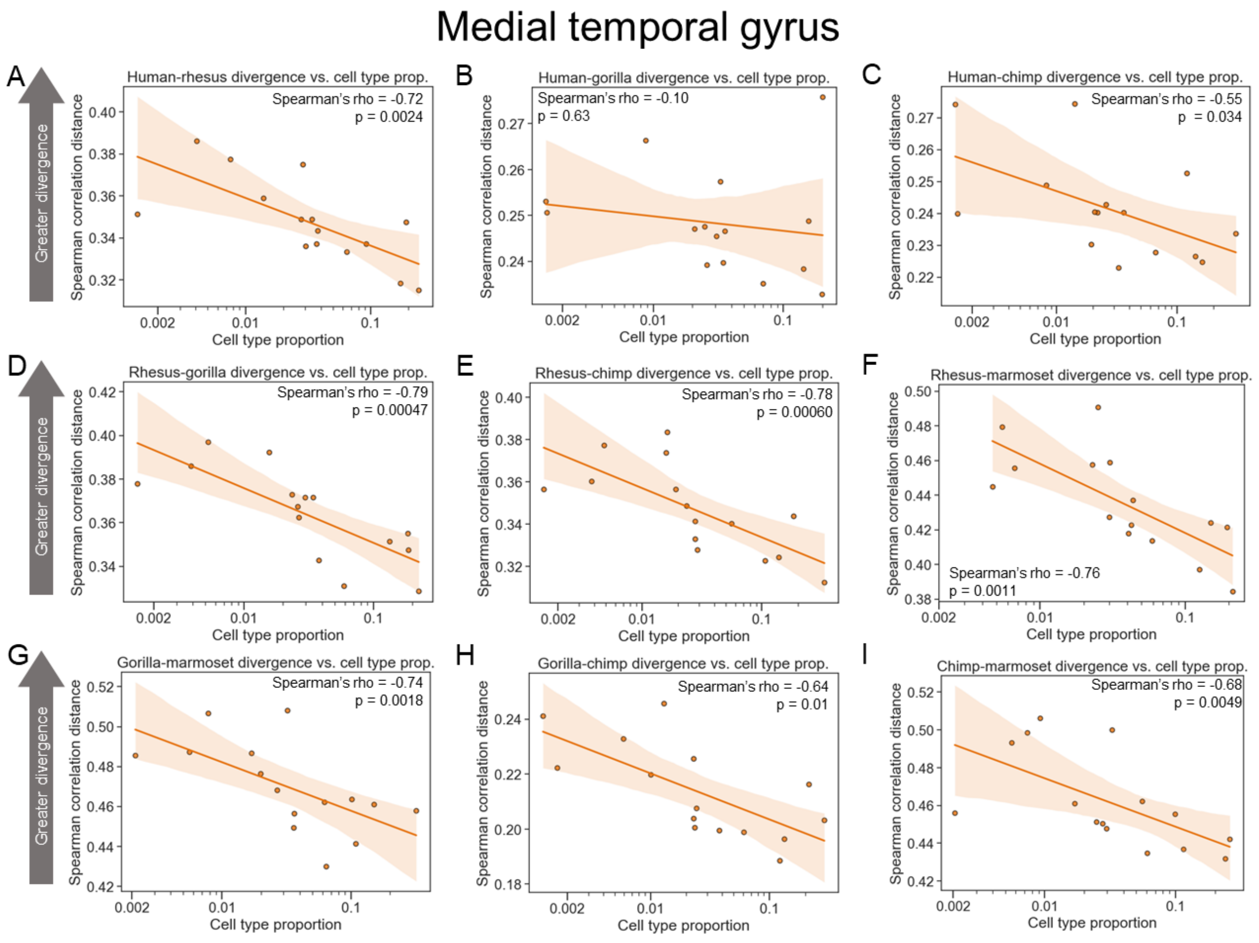


**Supplemental Figure 13: More common excitatory neuronal subtypes evolve more slowly than rarer subtypes in the MTG. A)** Plot showing the correlation between neuronal subtype proportion ($\log_{10}$ scale on the x-axis) and subtype-specific divergence between human and rhesus in the MTG for excitatory neurons. **B)** Same as in (A) but for human and gorilla. **C)** Same as in (A) but for human and chimp. **D)** Same as in (A) but for rhesus and gorilla. **E)** Same as in (A) but for rhesus and chimp. **F)** Same as in (A) but for rhesus and marmoset. **G)** Same as in (A) but for gorilla and marmoset. **H)** Same as in (A) but for gorilla and chimp. **I)** Same as in (A) but for chimp and marmoset.

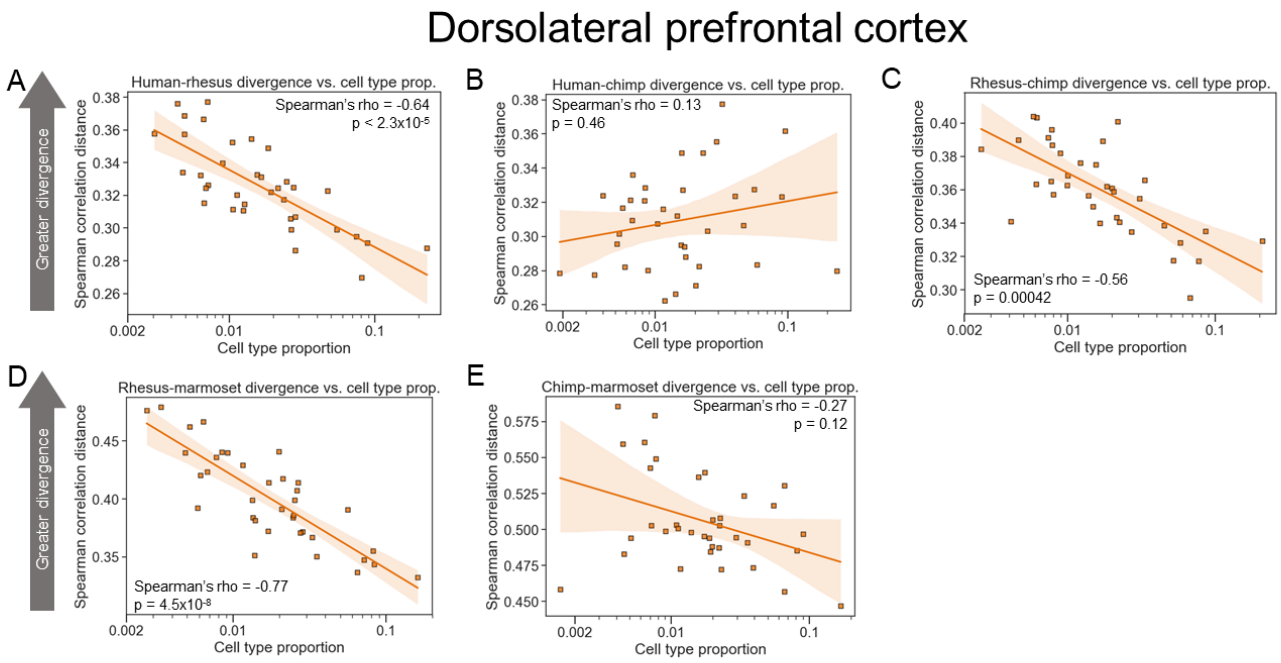


**Supplemental Figure 14: More common excitatory neuronal subtypes evolve more slowly than rarer subtypes in the DLPFC. A)** Plot showing the correlation between neuronal subtype proportion ($log_{10}$ scale on the x-axis) and subtype-specific divergence between human and rhesus in the DLPFC for excitatory neurons. **B)** Same as in (A) but for human and chimp. **C)** Same as in (A) but for rhesus and chimp. **D)** Same as in (A) but for rhesus and marmoset. **E)** Same as in (A) but for chimp and marmoset.

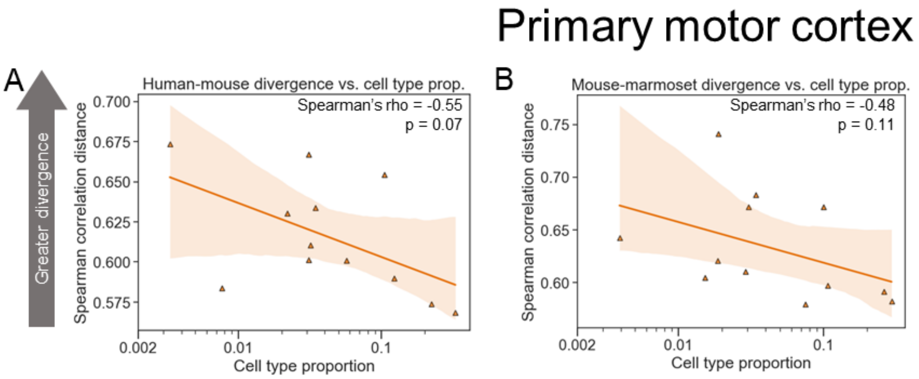


**Supplemental Figure 15: More common excitatory neuronal subtypes evolve more slowly than rarer subtypes in M1. A)** Plot showing the correlation between neuronal subtype proportion ($log_{10}$ scale on the x-axis) and subtype-specific divergence between human and mouse in M1 for excitatory neurons. **B)** Same as in (A) but for mouse and marmoset.

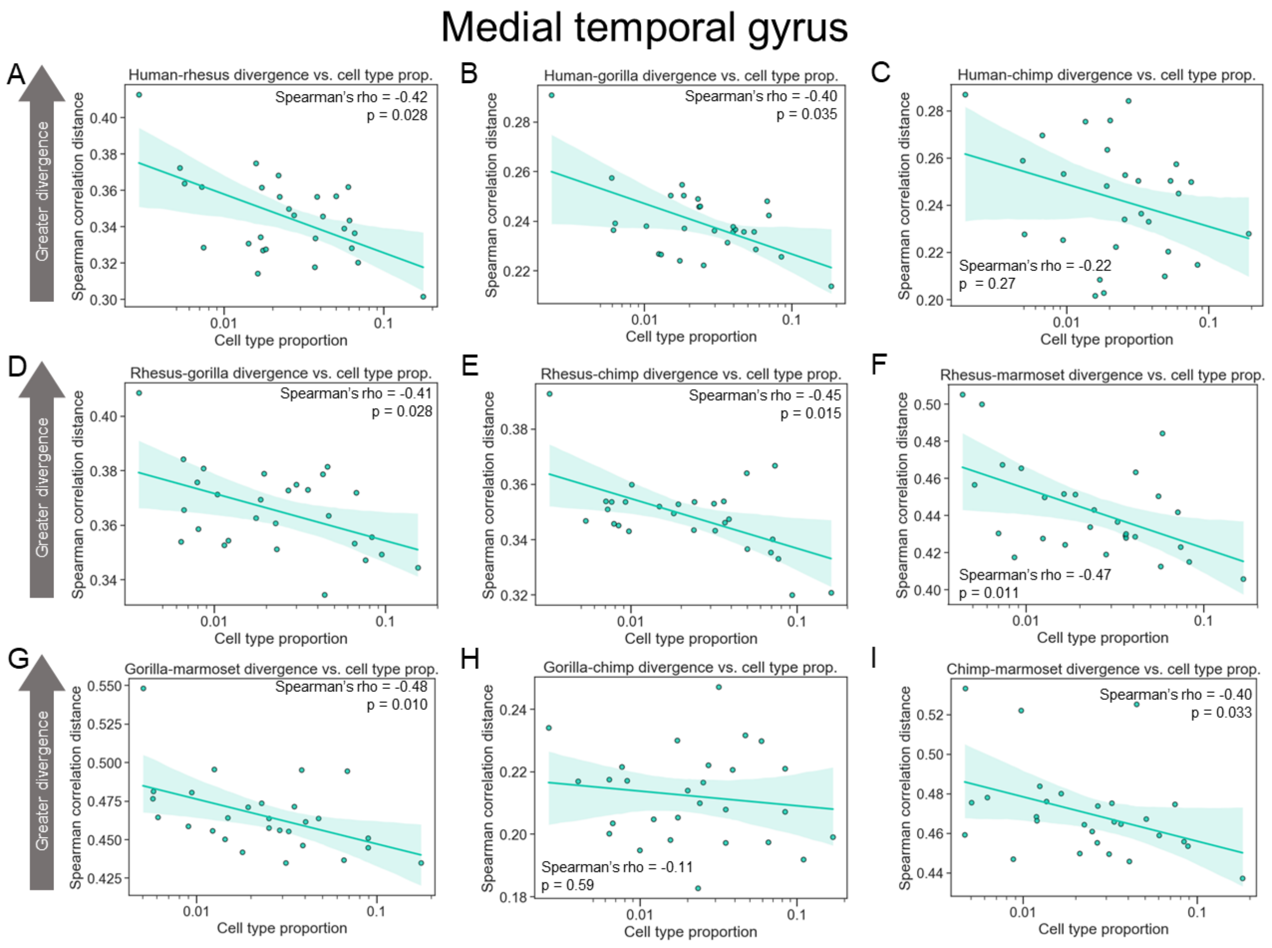


**Supplemental Figure 16: More common inhibitory neuronal subtypes evolve more slowly than rarer subtypes in the MTG. A)** Plot showing the correlation between neuronal subtype proportion ($\log_{10}$ scale on the x-axis) and subtype-specific divergence between human and rhesus in the MTG for inhibitory neurons. **B)** Same as in (A) but for human and gorilla. **C)** Same as in (A) but for human and chimp. **D)** Same as in (A) but for rhesus and gorilla. **E)** Same as in (A) but for rhesus and chimp. **F)** Same as in (A) but for rhesus and marmoset. **G)** Same as in (A) but for gorilla and marmoset. **H)** Same as in (A) but for gorilla and chimp. **I)** Same as in (A) but for chimp and marmoset.

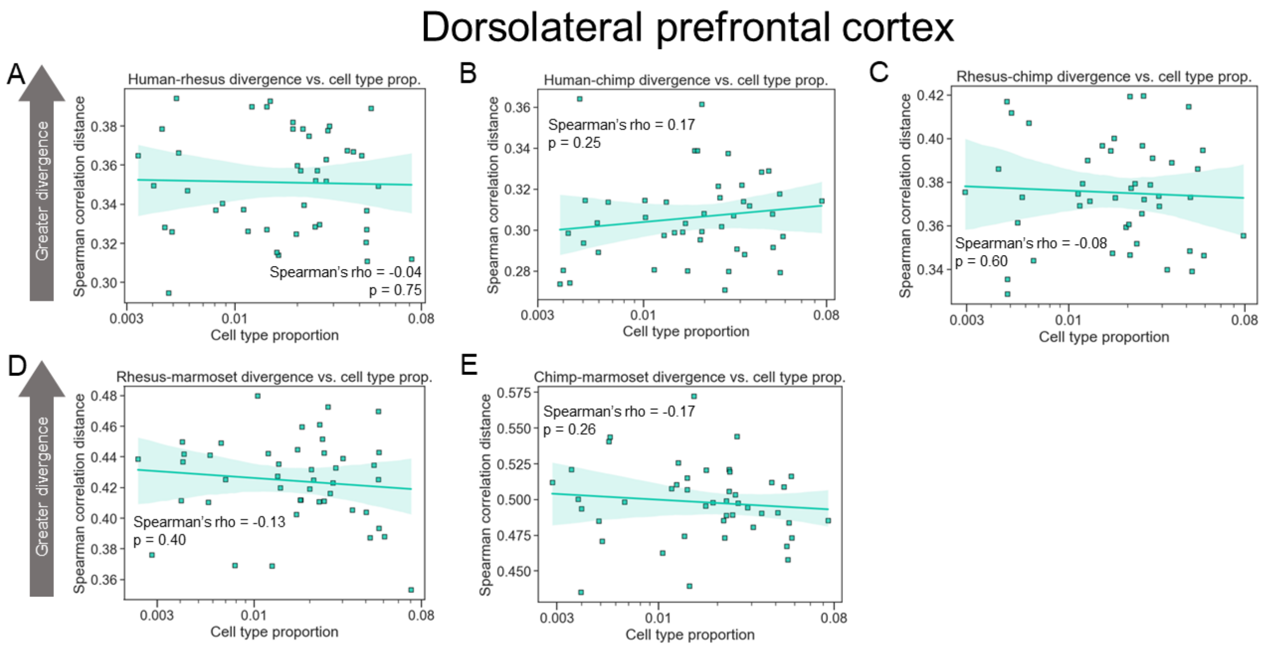


**Supplemental Figure 17: Weak relationship between inhibitory neuron subtype proportion and divergence in the DLPFC. A)** Plot showing the correlation between neuronal subtype proportion ($\log_{10}$ scale on the x-axis) and subtype-specific divergence between human and rhesus in the DLPFC for inhibitory neurons. **B)** Same as in (A) but for human and chimp. **C)** Same as in (A) but for rhesus and chimp. **D)** Same as in (A) but for rhesus and marmoset. **E)** Same as in (A) but for chimp and marmoset.

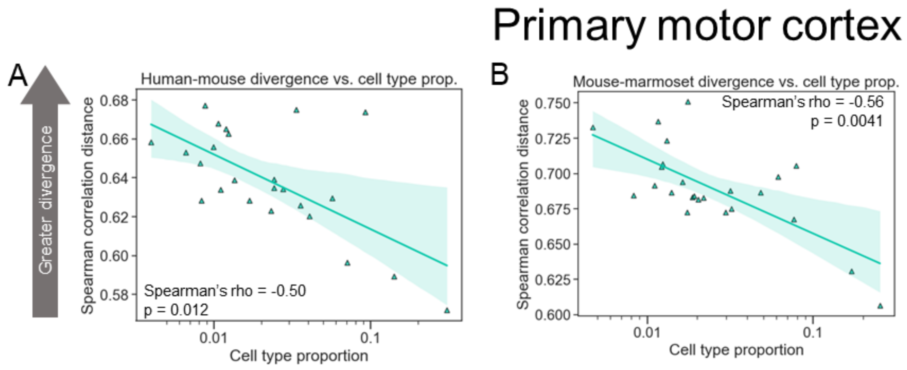


**Supplemental Figure 18: More common inhibitory neuronal subtypes evolve more slowly than rarer subtypes in M1. A)** Plot showing the correlation between neuronal subtype proportion ($log_{10}$ scale on the x-axis) and subtype-specific divergence between human and mouse in M1 for inhibitory neurons. **B)** Same as in (A) but for mouse and marmoset.

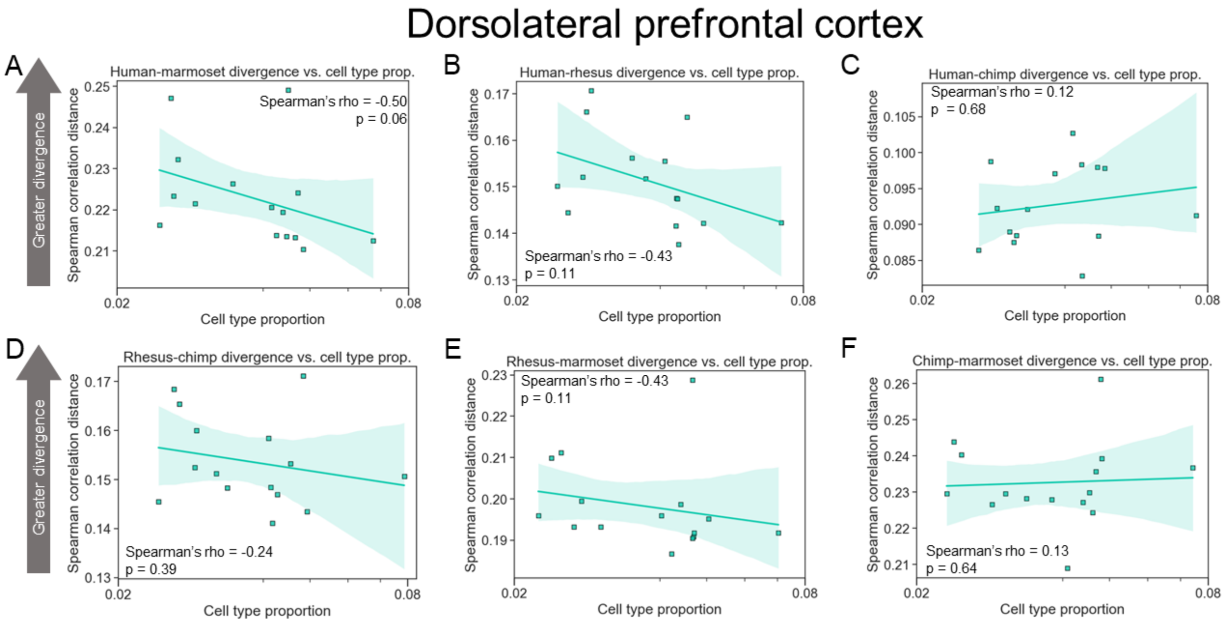


**Supplemental Figure 19: Stronger relationship between inhibitory neuron subtype proportion and divergence in the DLPFC for more abundant subtypes. A)** Plot showing the correlation between neuronal subtype proportion ($log_{10}$ scale on the x-axis) and subtype-specific divergence between human and rhesus in the DLPFC for inhibitory neurons. **B)** Same as in (A) but for human and chimp. **C)** Same as in (A) but for rhesus and chimp. **D)** Same as in (A) but for rhesus and marmoset. **E)** Same as in (A) but for chimp and marmoset.

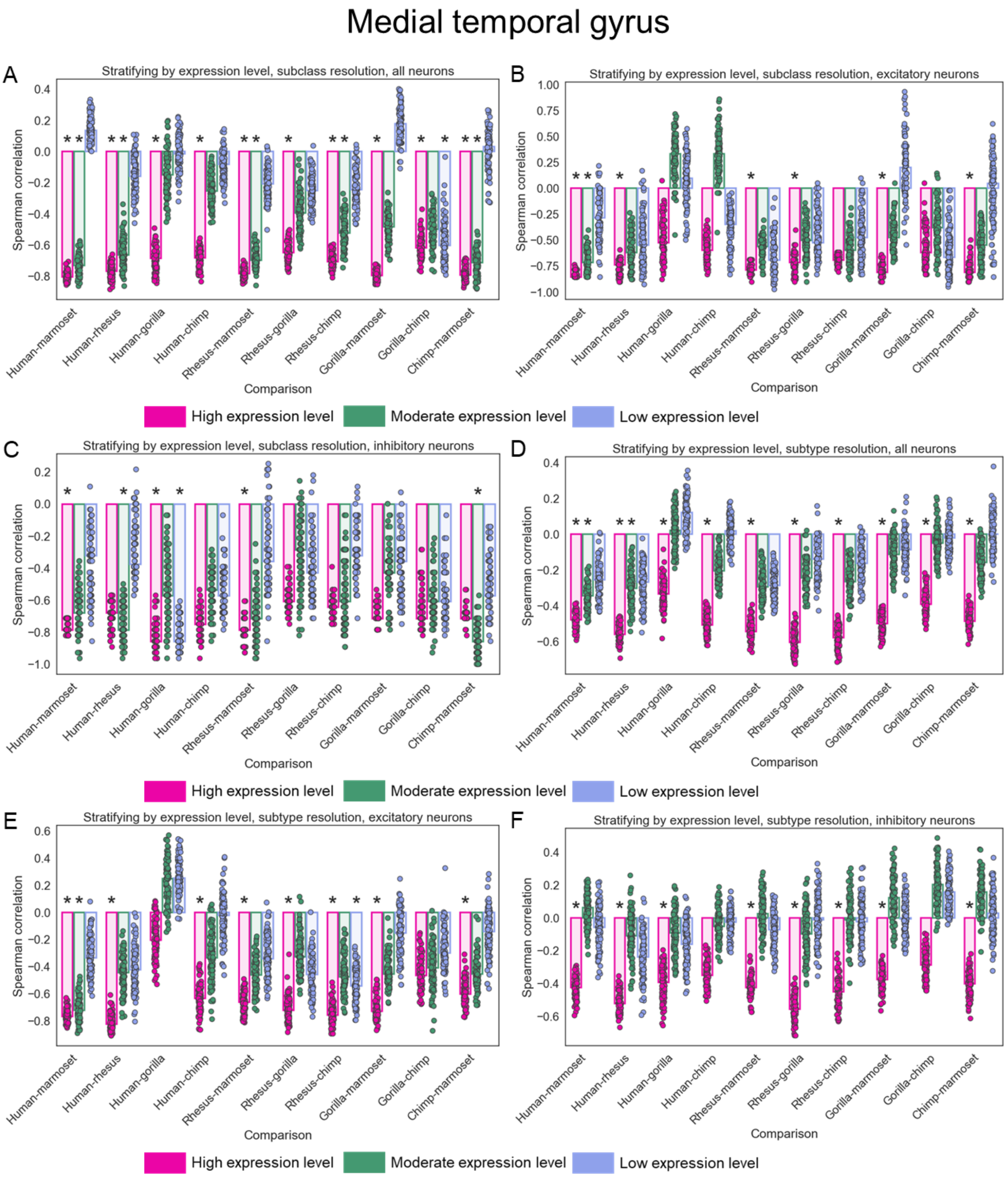


**Supplemental Figure 20: Correlation between cell type proportion and evolutionary divergence stratifying by expression level in the MTG. A)** Barplot showing the median Spearman correlation across 100 independent down-samplings for highly, moderately, and lowly expressed genes at the subclass level in the MTG dataset. Each species comparison is shown separately on the x-axis and colors correspond to the expression level bin. Each point represents the Spearman's rho for one down-sampling. Asterisks indicate if the median p-value across the 100 down-samplings was less than 0.05. **B)** Same as in (A) but for only excitatory neurons. **C)** Same as in (A) but for only inhibitory neurons. **D)** Same as in (A) but at the subtype level. **E)** Same as in (B) but at the subtype level. **F)** Same as in (C) but at the subtype level.

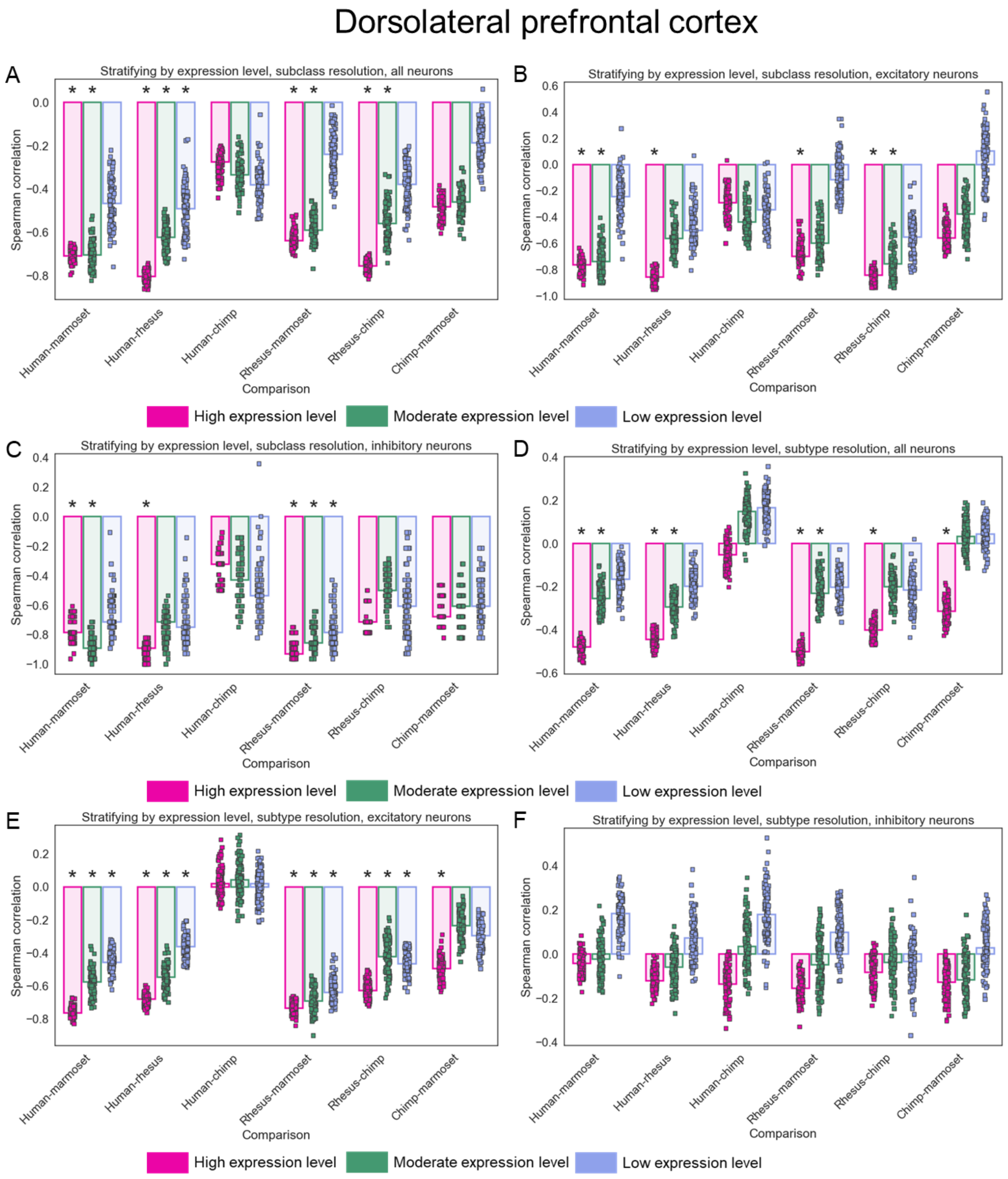


**Supplemental Figure 21: Correlation between cell type proportion and evolutionary divergence stratifying by expression level in the DLPFC. A)** Barplot showing the median Spearman correlation across 100 independent down-samplings for highly, moderately, and lowly expressed genes at the subclass level in the DLPFC dataset. Each species comparison is shown separately on the x-axis and colors correspond to the expression level bin. Each point represents the Spearman's rho for one down-sampling. Asterisks indicate if the median p-value across the 100 down-samplings was less than 0.05. **B)** Same as in (A) but for only excitatory neurons. **C)** Same as in (A) but for only inhibitory neurons. **D)** Same as in (A) but at the subtype level. **E)** Same as in (B) but at the subtype level. **F)** Same as in (C) but at the subtype level.

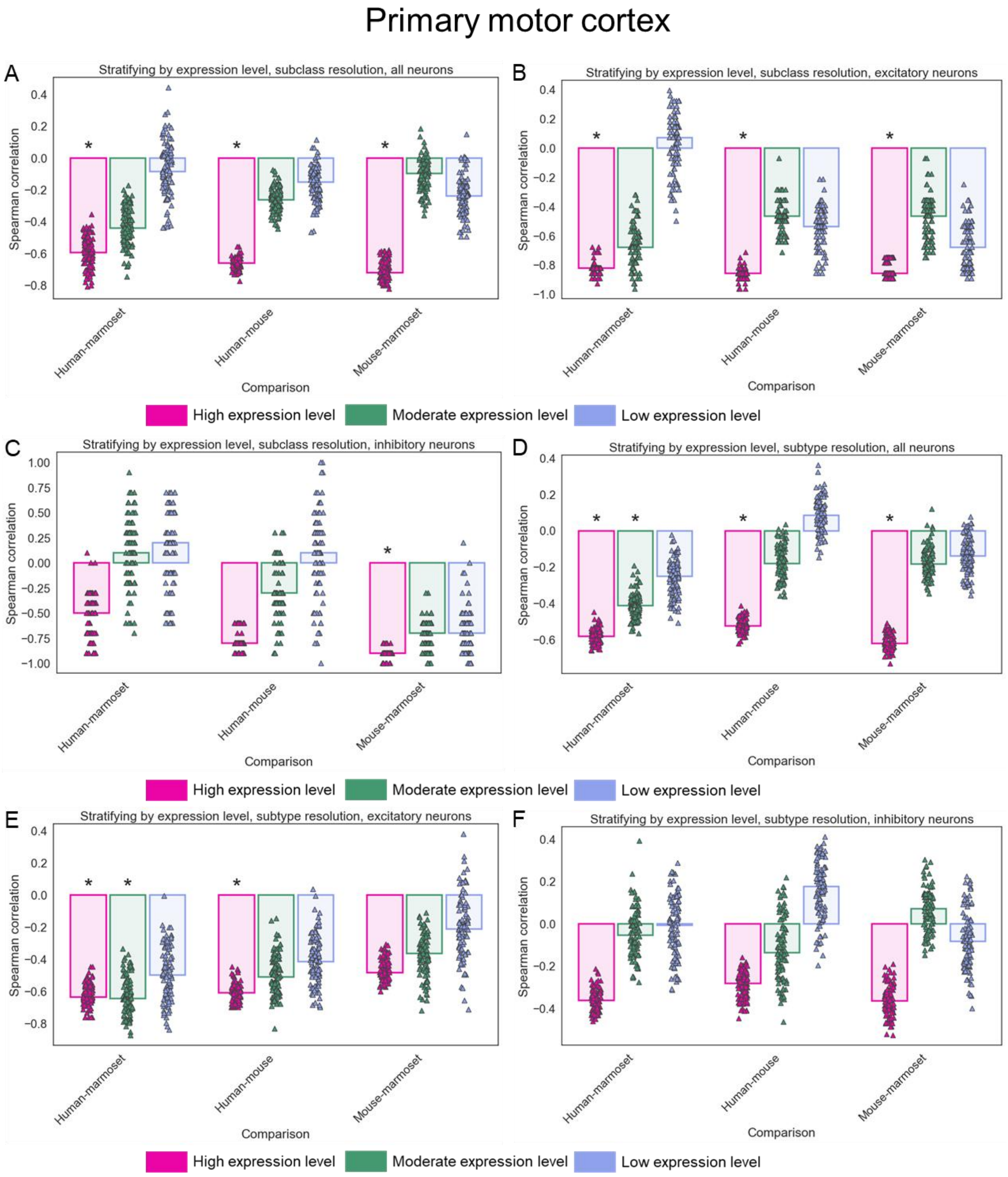


**Supplemental Figure 22: Correlation between cell type proportion and evolutionary divergence stratifying by expression level in M1. A)** Barplot showing the median Spearman correlation across 100 independent down-samplings for highly, moderately, and lowly expressed genes at the subclass level in the M1 dataset. Each species comparison is shown separately on the x-axis and colors correspond to the expression level bin. Each point represents the Spearman's rho for one down-sampling. Asterisks indicate if the median p-value across the 100 down-samplings was less than 0.05. **B)** Same as in (A) but for only excitatory neurons. **C)** Same as in (A) but for only inhibitory neurons. **D)** Same as in (A) but at the subtype level. **E)** Same as in (B) but at the subtype level. **F)** Same as in (C) but at the subtype level.

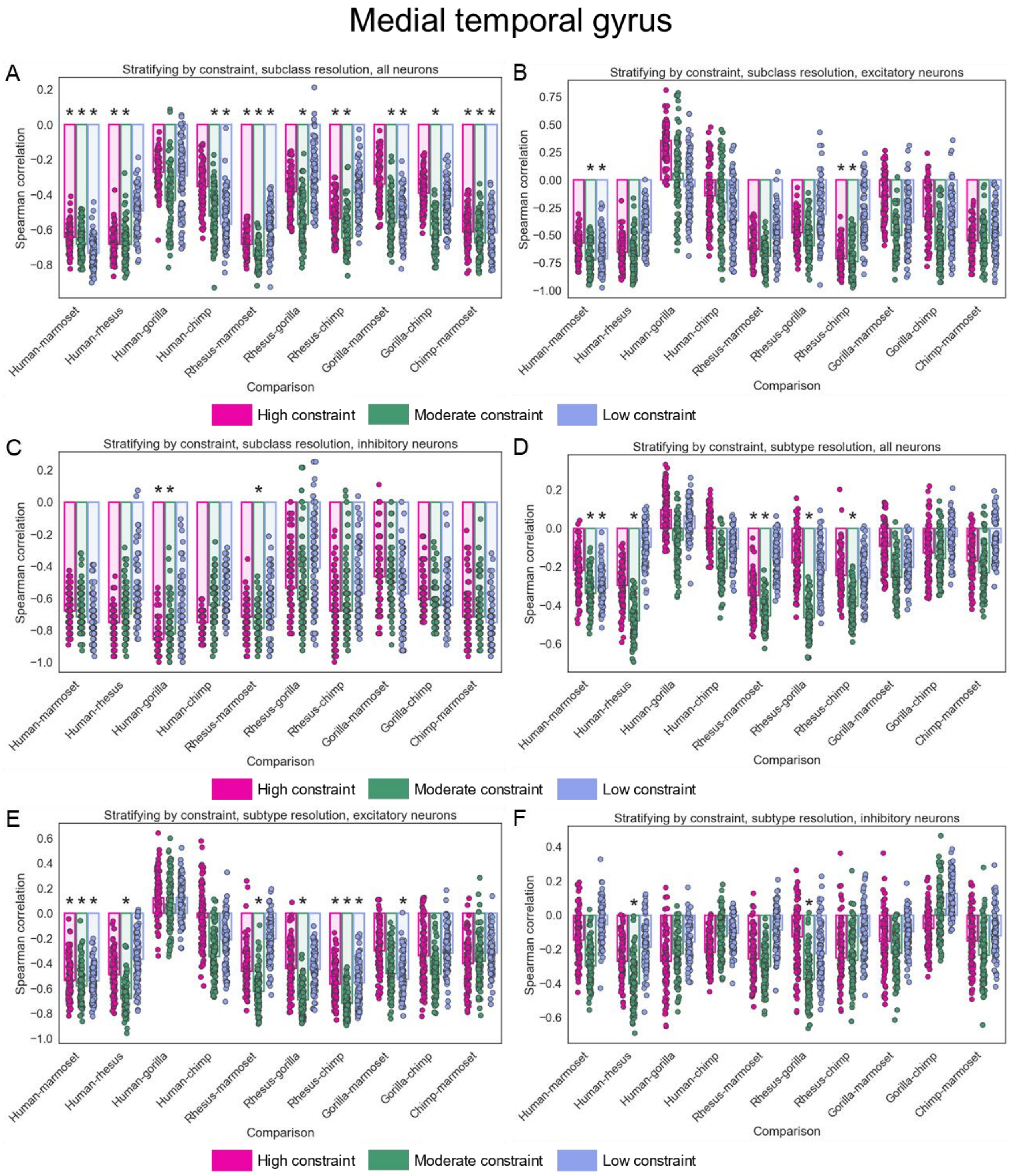


**Supplemental Figure 23: Correlation between cell type proportion and evolutionary divergence stratifying by constraint in the MTG. A)** Barplot showing the median Spearman correlation across 100 independent down-samplings for highly, moderately, and lowly evolutionarily constrained genes at the subclass level in the MTG dataset. Each species comparison is shown separately on the x-axis and colors correspond to the constraint bin. Each point represents the Spearman's rho for one down-sampling. Asterisks indicate if the median p-value across the 100 down-samplings was less than 0.05. **B)** Same as in (A) but for only excitatory neurons. **C)** Same as in (A) but for only inhibitory neurons. **D)** Same as in (A) but at the subtype level. **E)** Same as in (B) but at the subtype level. **F)** Same as in (C) but at the subtype level.

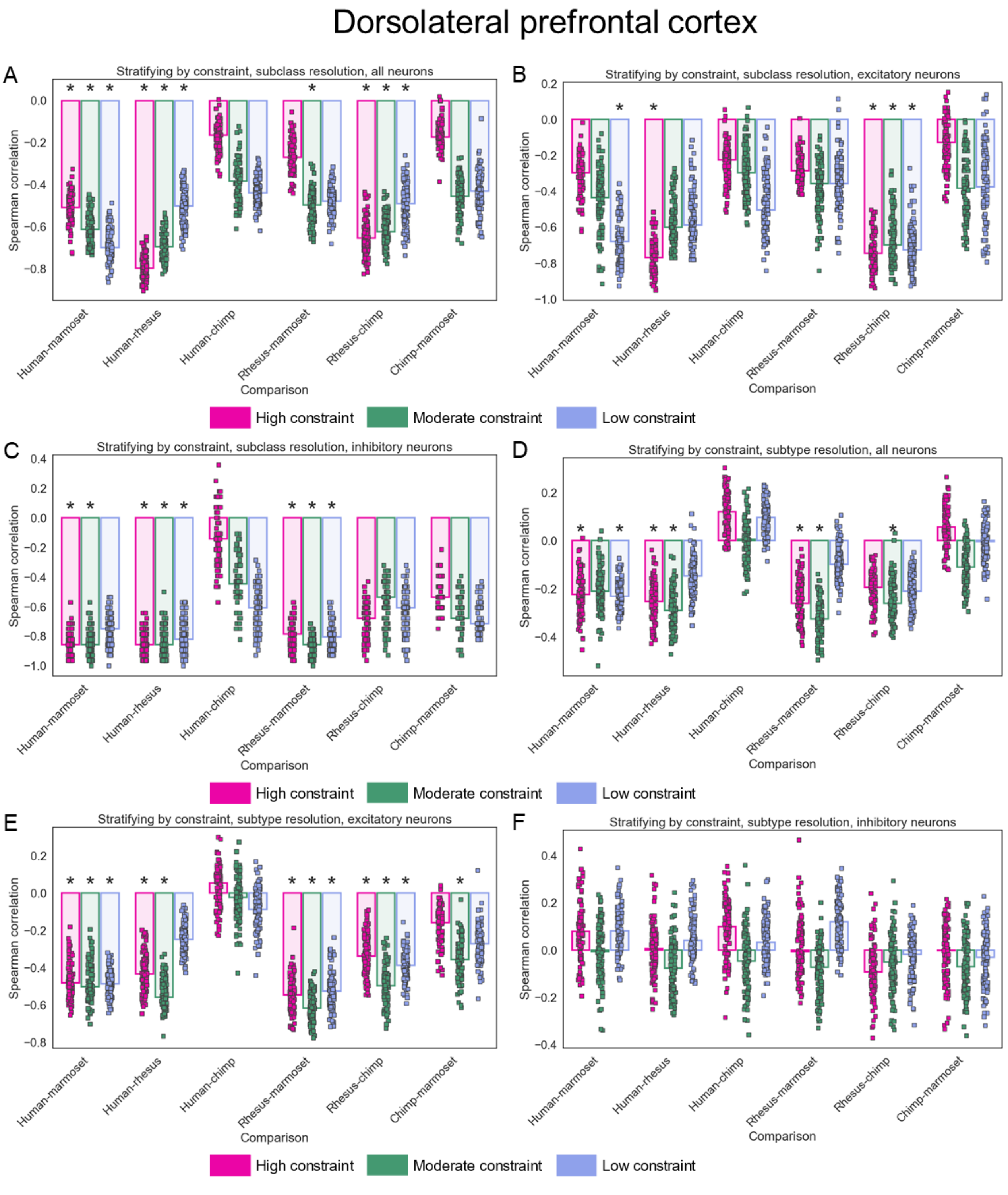


**Supplemental Figure 24: Correlation between cell type proportion and evolutionary divergence stratifying by constraint in the DLPFC. A)** Barplot showing the median Spearman correlation across 100 independent down-samplings for highly, moderately, and lowly evolutionarily constrained genes at the subclass level in the DLPFC dataset. Each species comparison is shown separately on the x-axis and colors correspond to the constraint bin. Each point represents the Spearman's rho for one down-sampling. Asterisks indicate if the median p-value across the 100 down-samplings was less than 0.05. **B)** Same as in (A) but for only excitatory neurons. **C)** Same as in (A) but for only inhibitory neurons. **D)** Same as in (A) but at the subtype level. **E)** Same as in (B) but at the subtype level. **F)** Same as in (C) but at the subtype level.

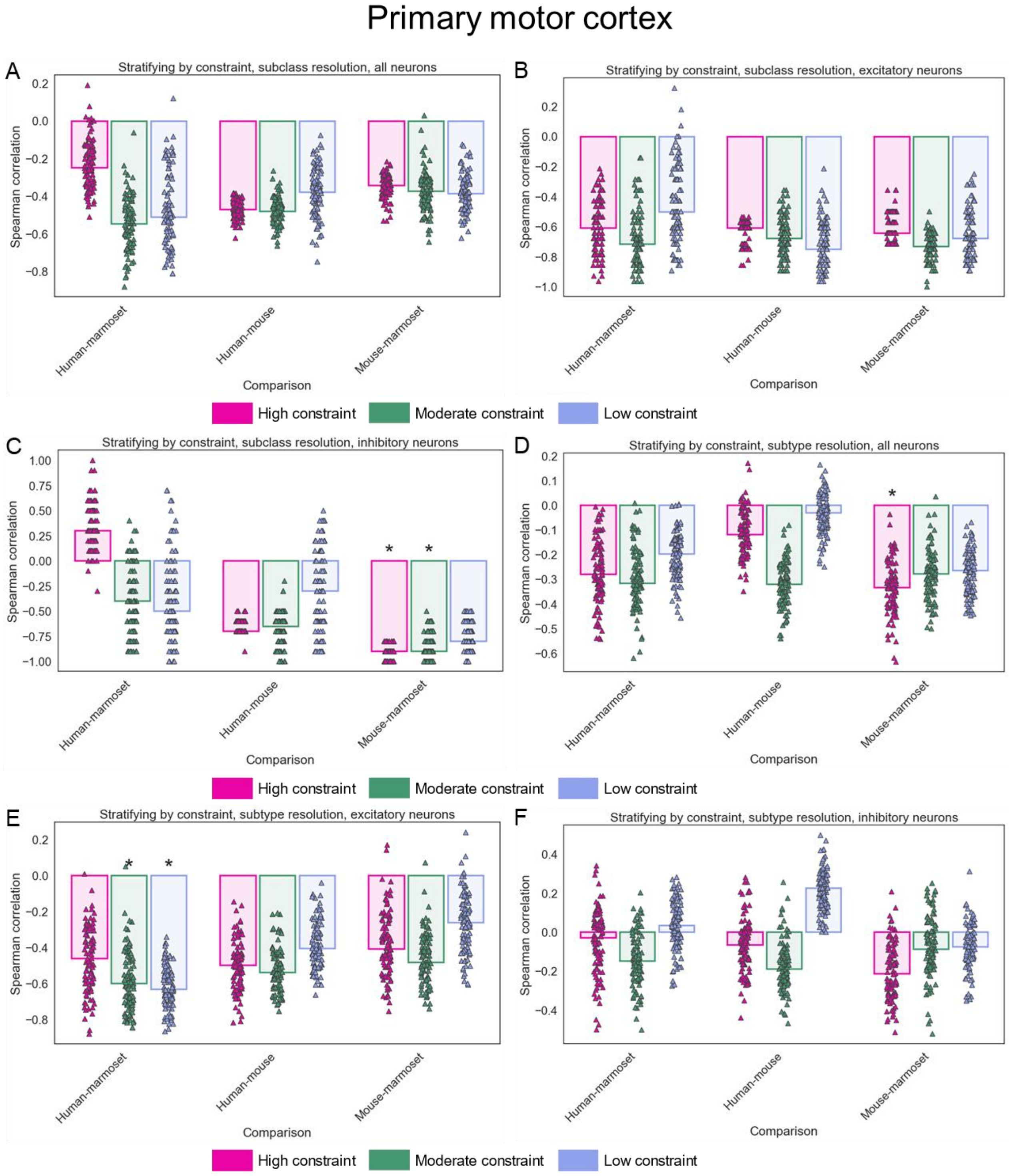


**Supplemental Figure 25: Correlation between cell type proportion and evolutionary divergence stratifying by constraint in M1. A)** Barplot showing the median Spearman correlation across 100 independent down-samplings for highly, moderately, and lowly evolutionarily constrained genes at the subclass level in the M1 dataset. Each species comparison is shown separately on the x-axis and colors correspond to the constraint bin. Each point represents the Spearman's rho for one down-sampling. Asterisks indicate if the median p-value across the 100 down-samplings was less than 0.05. **B)** Same as in (A) but for only excitatory neurons. **C)** Same as in (A) but for only inhibitory neurons. **D)** Same as in (A) but at the subtype level. **E)** Same as in (B) but at the subtype level. **F)** Same as in (C) but at the subtype level.

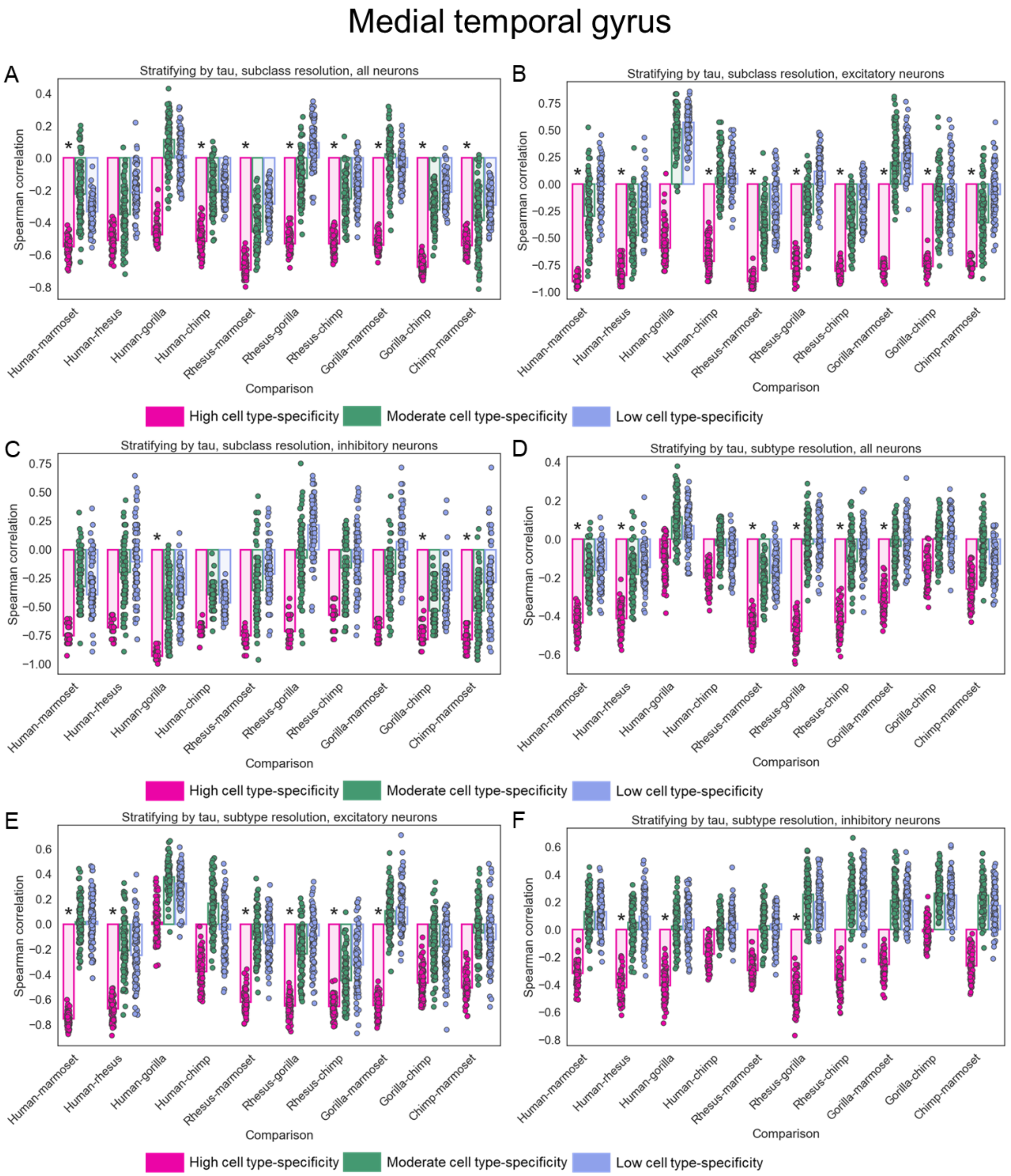


**Supplemental Figure 26: Correlation between cell type proportion and evolutionary divergence stratifying by cell type-specificity of expression in the MTG. A)** Barplot showing the median Spearman correlation across 100 independent down-samplings for lowly, moderately, and highly broadly expressed genes at the subclass level in the MTG dataset. Each species comparison is shown separately on the x-axis and colors correspond to the cell type-specificity bin. Each point represents the Spearman's rho for one down-sampling. Asterisks indicate if the median p-value across the 100 down-samplings was less than 0.05. **B)** Same as in (A) but for only excitatory neurons. **C)** Same as in (A) but for only inhibitory neurons. **D)** Same as in (A) but at the subtype level. **E)** Same as in (B) but at the subtype level. **F)** Same as in (C) but at the subtype level.

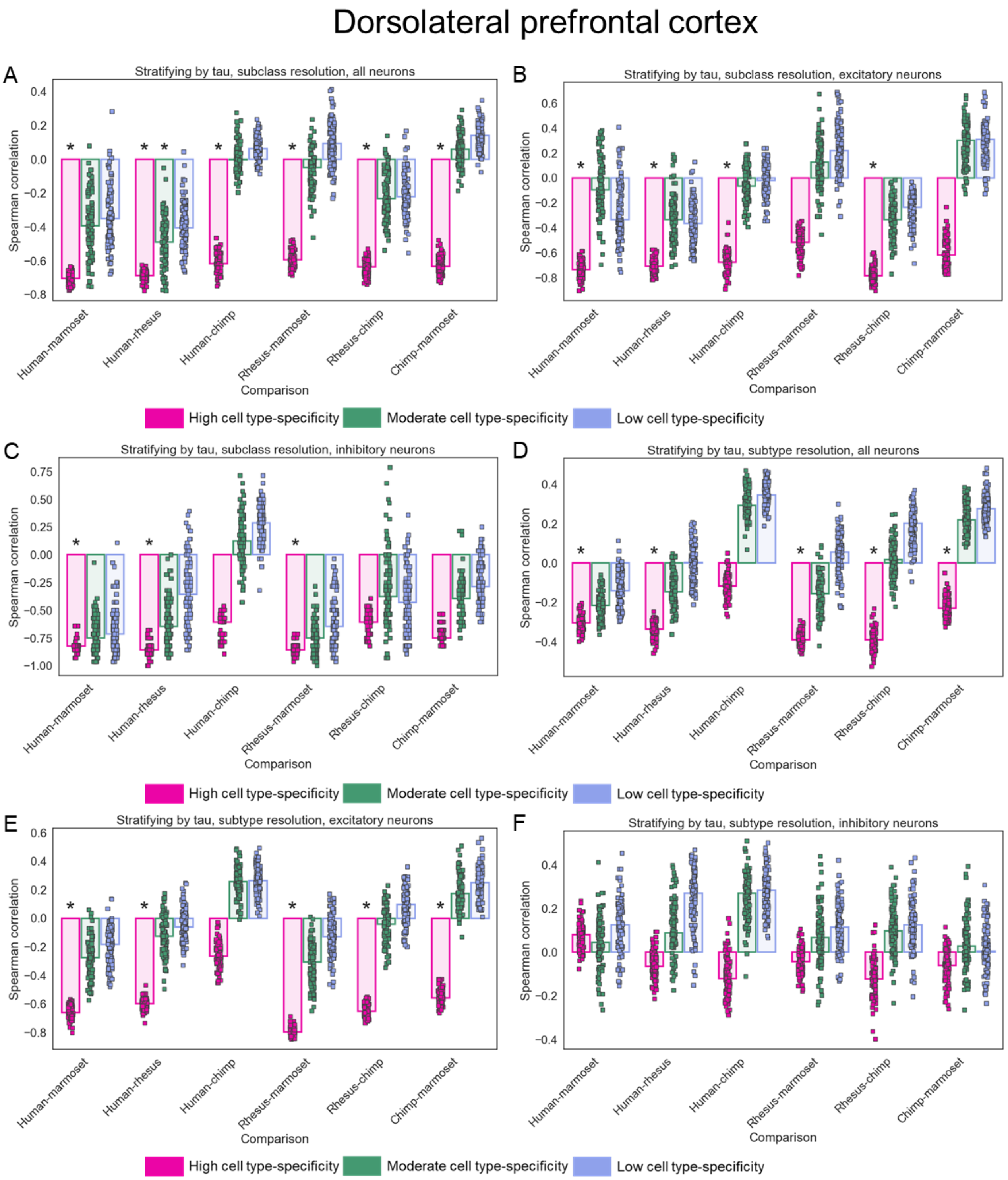


**Supplemental Figure 27: Correlation between cell type proportion and evolutionary divergence stratifying by cell type-specificity of expression in the DLPFC. A)** Barplot showing the median Spearman correlation across 100 independent down-samplings for lowly, moderately, and highly broadly expressed genes at the subclass level in the DLPFC dataset. Each species comparison is shown separately on the x-axis and colors correspond to the cell type-specificity bin. Each point represents the Spearman's rho for one down-sampling. Asterisks indicate if the median p-value across the 100 down-samplings was less than 0.05. **B)** Same as in (A) but for only excitatory neurons. **C)** Same as in (A) but for only inhibitory neurons. **D)** Same as in (A) but at the subtype level. **E)** Same as in (B) but at the subtype level. **F)** Same as in (C) but at the subtype level.

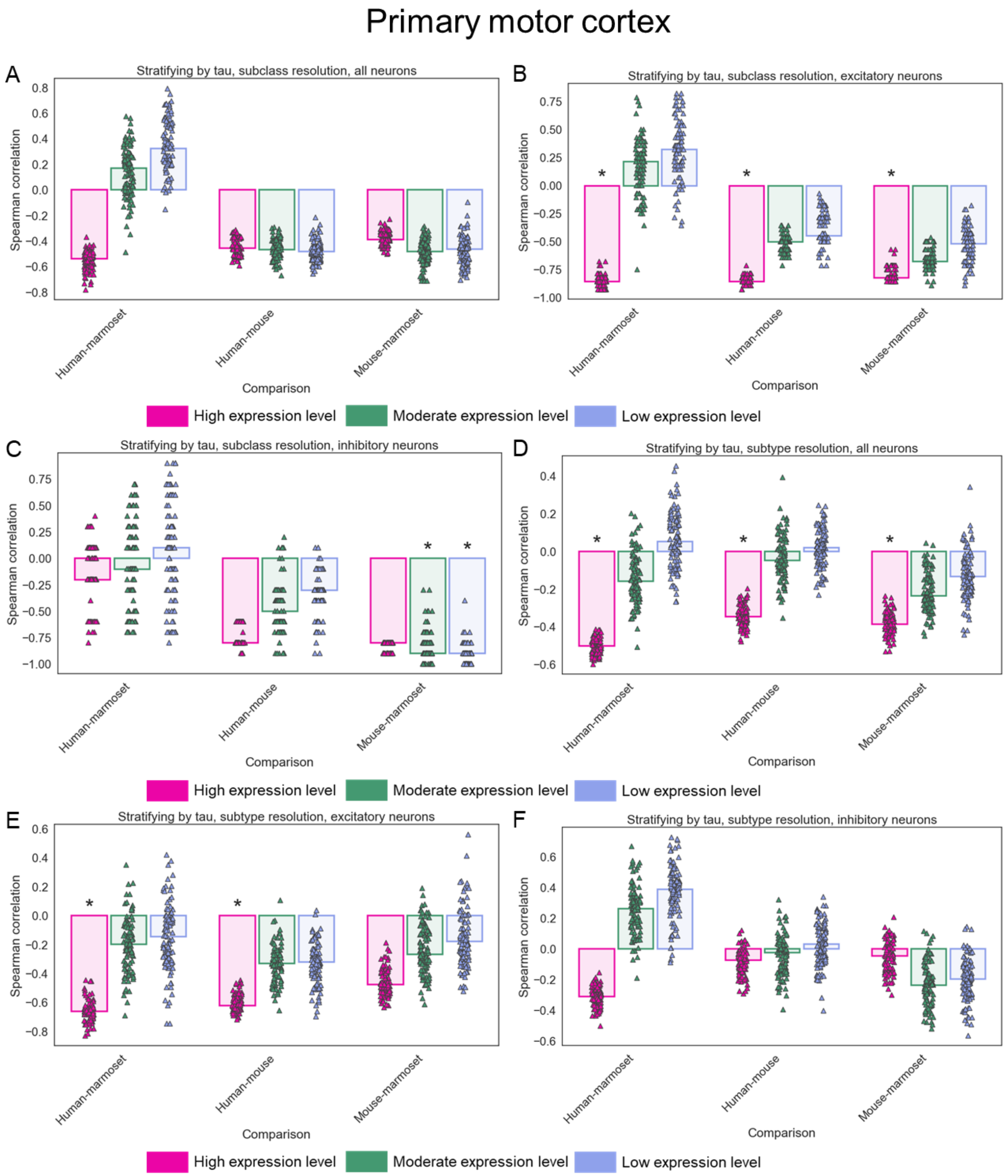


**Supplemental Figure 28: Correlation between cell type proportion and evolutionary divergence stratifying by cell type-specificity of expression in M1. A)** Barplot showing the median Spearman correlation across 100 independent down-samplings for lowly, moderately, and highly broadly expressed genes at the subclass level in the M1 dataset. Each species comparison is shown separately on the x-axis and colors correspond to the cell type-specificity bin. Each point represents the Spearman's rho for one down-sampling. Asterisks indicate if the median p-value across the 100 down-samplings was less than 0.05. **B)** Same as in (A) but for only excitatory neurons. **C)** Same as in (A) but for only inhibitory neurons. **D)** Same as in (A) but at the subtype level. **E)** Same as in (B) but at the subtype level. **F)** Same as in (C) but at the subtype level.

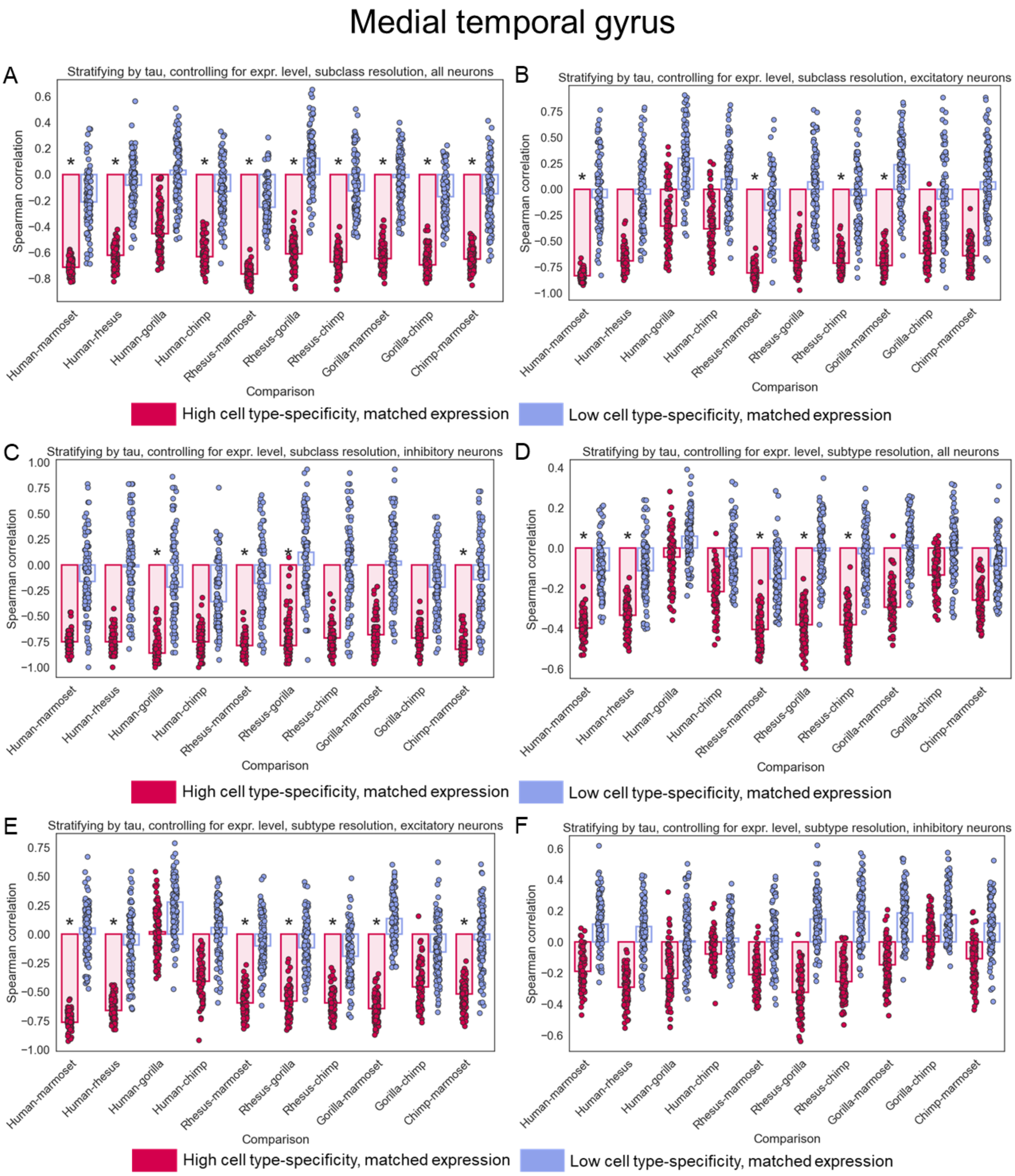


**Supplemental Figure 29: Correlation between cell type proportion and evolutionary divergence stratifying by cell type-specificity of expression and controlling for expression level in the MTG. A)** Barplot showing the median Spearman correlation across 100

independent down-samplings for genes with high cell type-specificity and low cell type-specificity matched for expression level at the subclass level in the MTG dataset. Each species comparison is shown separately on the x-axis and colors correspond to the cell type-specificity bin. Each point represents the Spearman’s rho for one down-sampling. Asterisks indicate if the median p-value across the 100 down-samplings was less than 0.05. **B)** Same as in (A) but for only excitatory neurons. **C)** Same as in (A) but for only inhibitory neurons. **D)** Same as in (A) but at the subtype level. **E)** Same as in (B) but at the subtype level. **F)** Same as in (C) but at the subtype level.

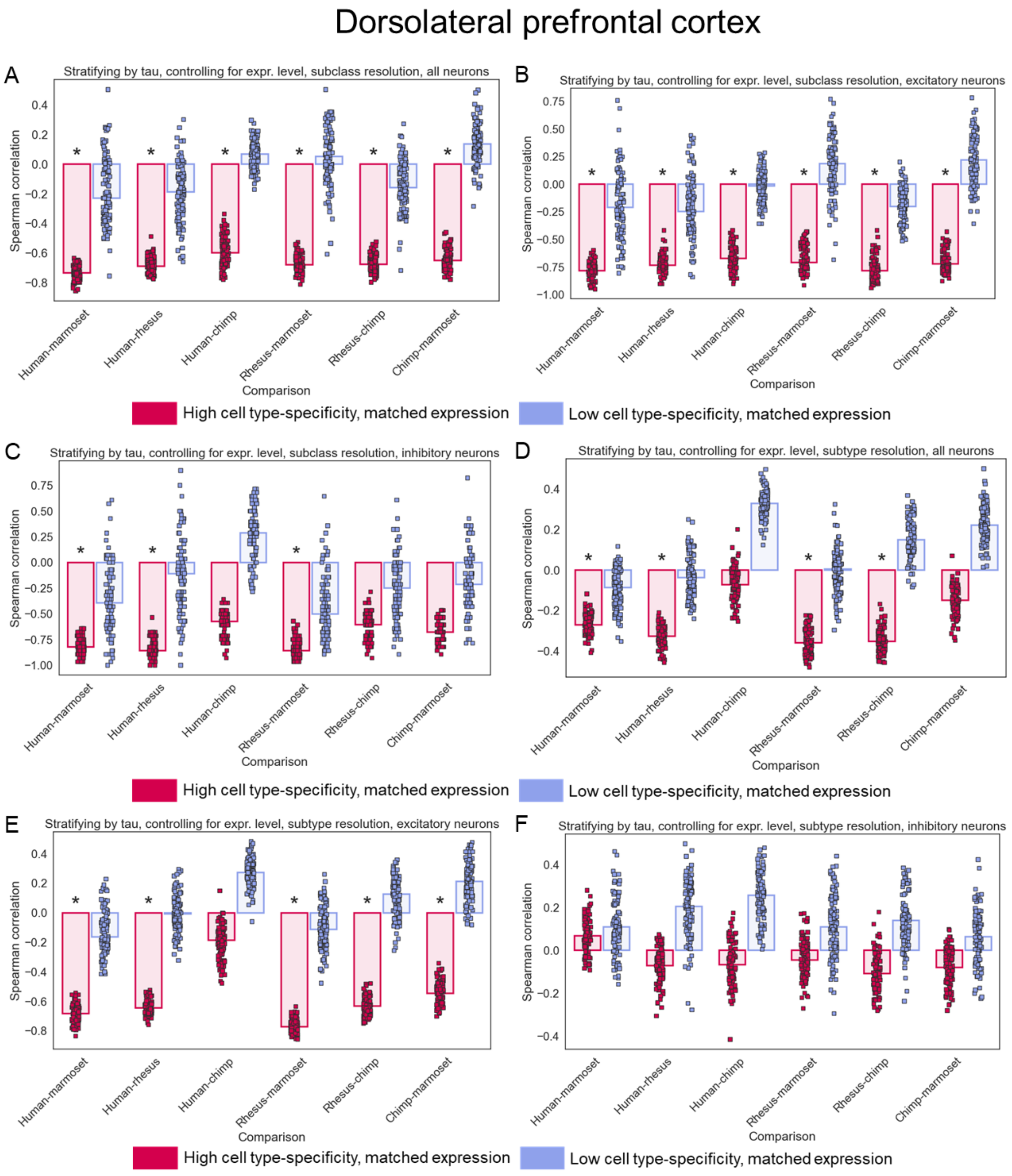


**Supplemental Figure 30: Correlation between cell type proportion and evolutionary divergence stratifying by cell type-specificity of expression and controlling for expression level in the DLPFC. A)** Barplot showing the median Spearman correlation across

100 independent down-samplings for genes with high cell type-specificity and low cell type-specificity matched for expression level at the subclass level in the DLPFC dataset. Each species comparison is shown separately on the x-axis and colors correspond to the cell type-specificity bin. Each point represents the Spearman's rho for one down-sampling. Asterisks indicate if the median p-value across the 100 down-samplings was less than 0.05. **B)** Same as in (A) but for only excitatory neurons. **C)** Same as in (A) but for only inhibitory neurons. **D)** Same as in (A) but at the subtype level. **E)** Same as in (B) but at the subtype level. **F)** Same as in (C) but at the subtype level.

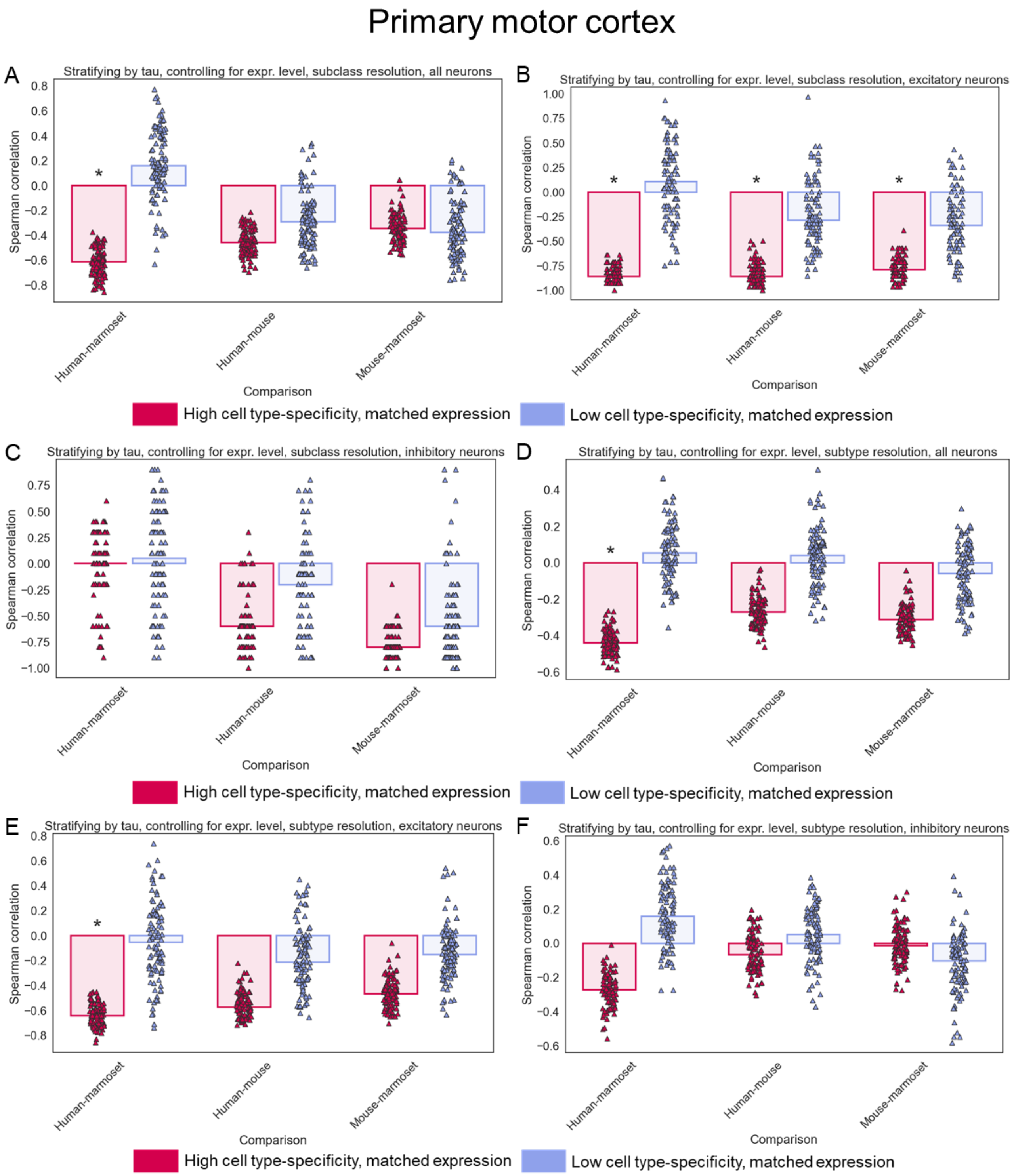


**Supplemental Figure 31: Correlation between cell type proportion and evolutionary divergence stratifying by cell type-specificity of expression and controlling for expression level in M1. A)** Barplot showing the median Spearman correlation across 100

independent down-samplings for genes with high cell type-specificity and low cell type-specificity matched for expression level at the subclass level in the M1 dataset. Each species comparison is shown separately on the x-axis and colors correspond to the cell type-specificity bin. Each point represents the Spearman's rho for one down-sampling. Asterisks indicate if the median p-value across the 100 down-samplings was less than 0.05. **B)** Same as in (A) but for only excitatory neurons. **C)** Same as in (A) but for only inhibitory neurons. **D)** Same as in (A) but at the subtype level. **E)** Same as in (B) but at the subtype level. **F)** Same as in (C) but at the subtype level.

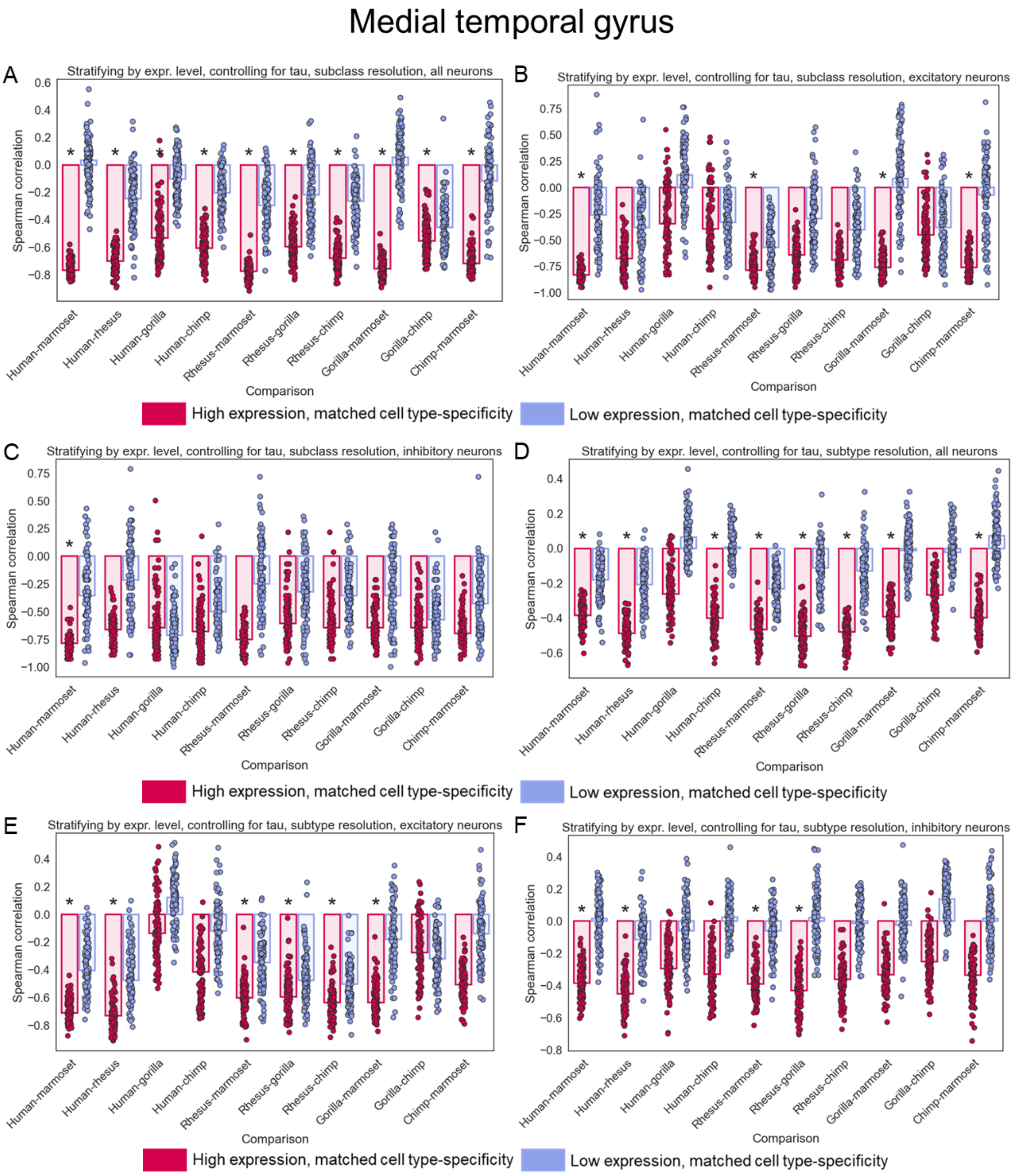


**Supplemental Figure 32: Correlation between cell type proportion and evolutionary divergence stratifying by expression level and controlling for cell type-specificity of expression in the MTG. A)** Barplot showing the median Spearman correlation across 100

independent down-samplings for highly and lowly expressed genes matched for cell type-specificity of expression at the subclass level in the MTG dataset. Each species comparison is shown separately on the x-axis and colors correspond to the expression level bin. Each point represents the Spearman's rho for one down-sampling. Asterisks indicate if the median p-value across the 100 down-samplings was less than 0.05. **B)** Same as in (A) but for only excitatory neurons. **C)** Same as in (A) but for only inhibitory neurons. **D)** Same as in (A) but at the subtype level. **E)** Same as in (B) but at the subtype level. **F)** Same as in (C) but at the subtype level.

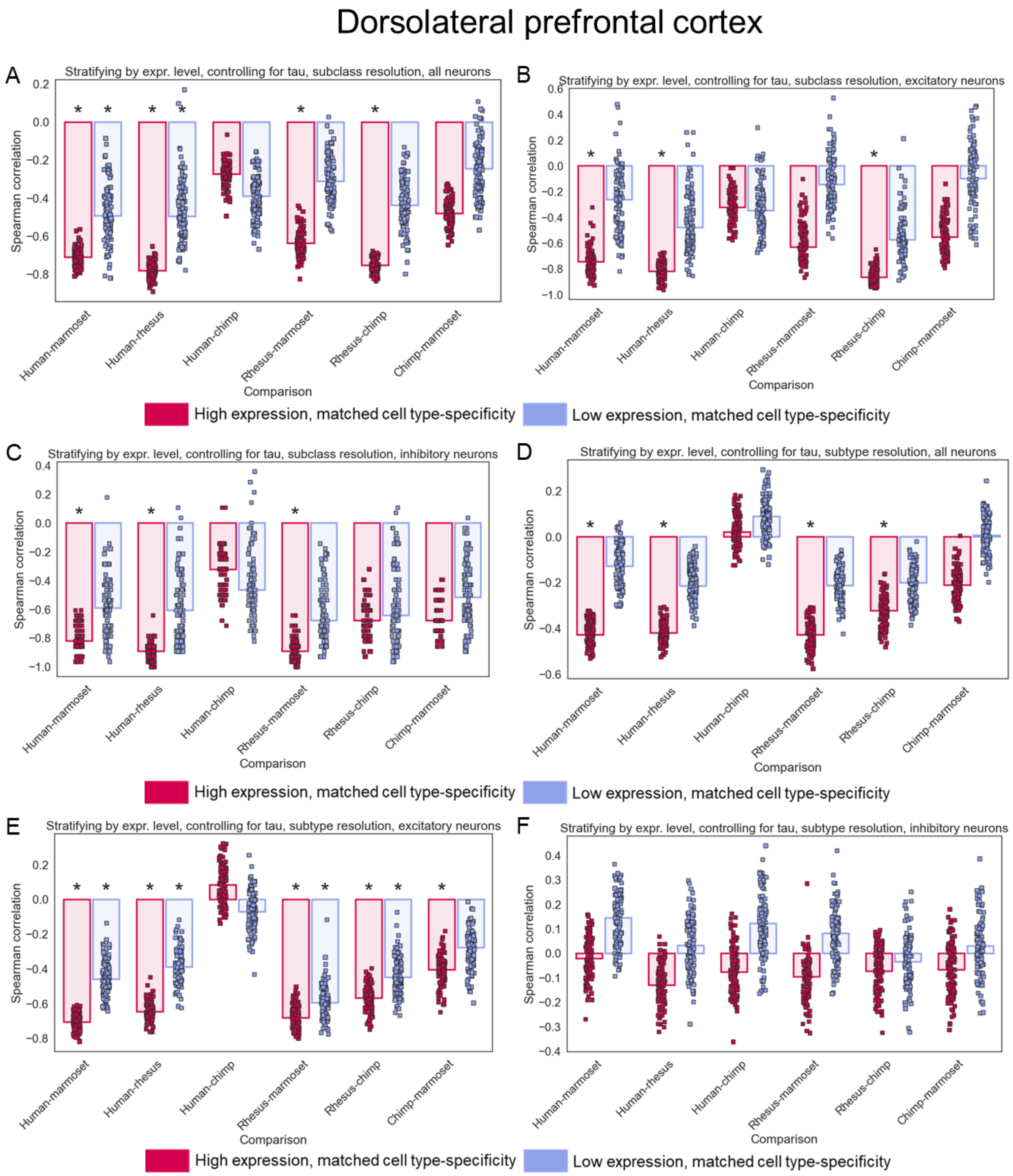


**Supplemental Figure 33: Correlation between cell type proportion and evolutionary divergence stratifying by expression level and controlling for cell type-specificity of expression in the DLPFC. A)** Barplot showing the median Spearman correlation across 100

independent down-samplings for highly and lowly expressed genes matched for cell type-specificity of expression at the subclass level in the DLPFC dataset. Each species comparison is shown separately on the x-axis and colors correspond to the expression level bin. Each point represents the Spearman's rho for one down-sampling. Asterisks indicate if the median p-value across the 100 down-samplings was less than 0.05. **B)** Same as in (A) but for only excitatory neurons. **C)** Same as in (A) but for only inhibitory neurons. **D)** Same as in (A) but at the subtype level. **E)** Same as in (B) but at the subtype level. **F)** Same as in (C) but at the subtype level.

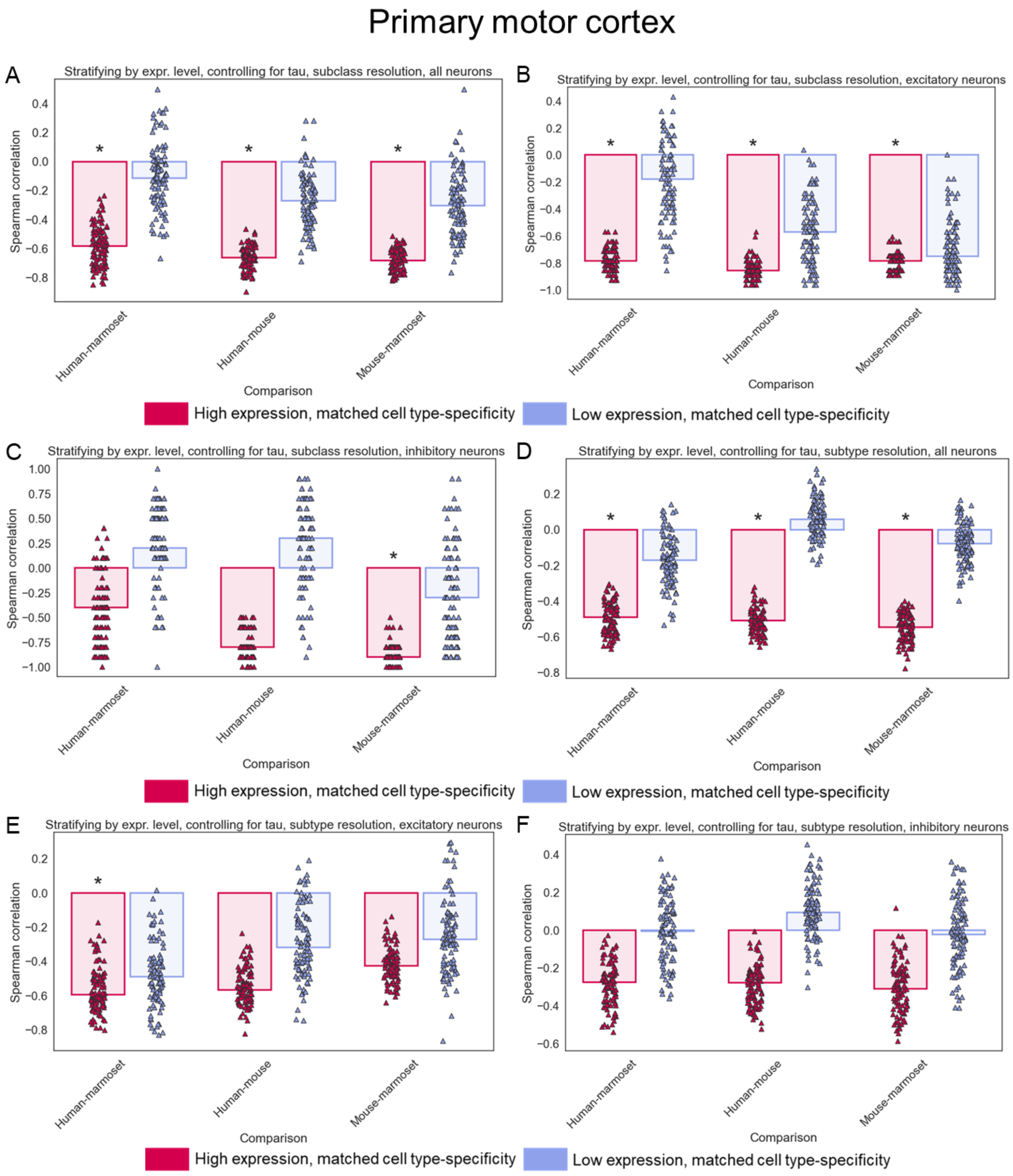


**Supplemental Figure 34: Correlation between cell type proportion and evolutionary divergence stratifying by expression level and controlling for cell type-specificity of expression in M1. A)** Barplot showing the median Spearman correlation across 100

independent down-samplings for highly and lowly expressed genes matched for cell type-specificity of expression at the subclass level in the M1 dataset. Each species comparison is shown separately on the x-axis and colors correspond to the expression level bin. Each point represents the Spearman's rho for one down-sampling. Asterisks indicate if the median p-value across the 100 down-samplings was less than 0.05. **B)** Same as in (A) but for only excitatory neurons. **C)** Same as in (A) but for only inhibitory neurons. **D)** Same as in (A) but at the subtype level. **E)** Same as in (B) but at the subtype level. **F)** Same as in (C) but at the subtype level.

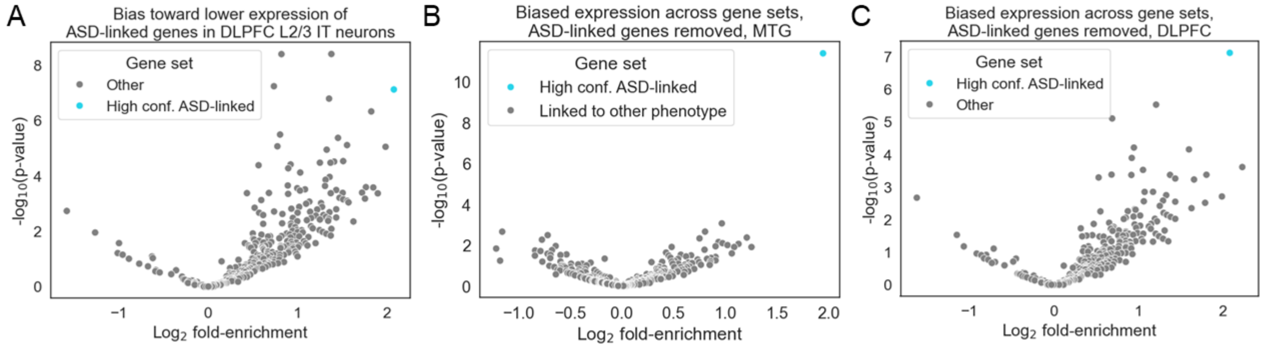


**Supplemental Figure 35: Bias toward down-regulation in ASD-linked genes. A)** Volcano plot showing the $\log_2$ fold-enrichment for down-regulation in humans (x-axis) and the $-\log_{10}$ binomial p-value (y-axis). SFARI high-confidence ASD-linked genes are shown in blue, all other categories of genes (taken from the Human Phenotype Onotology) are shown in grey. Data are from DLPFC L2/3 IT neurons. **B)** Same as in (A) but in the MTG and removing all ASD-linked genes from HPO gene sets. **C)** Same as in (B) but for DLPFC.

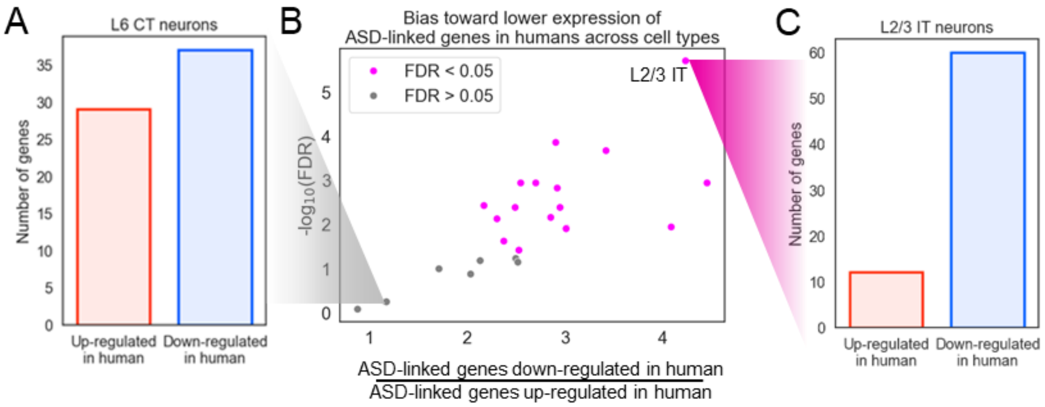


**Supplemental Figure 36: Down-regulation of ASD-linked genes in humans compared to chimpanzees in the DLPFC. A)** Barplot showing the number of high-confidence ASD-linked genes that are up-regulated vs. down-regulated in human relative to chimpanzee in DLPFC L6 CT neurons. **B)** Volcano plot showing the fold-enrichment for down-regulation in humans (x-axis) and the -$\log_{10}$ binomial FDR (y-axis). Subclasses with FDR < 0.05 are shown in red; only subclasses with at least 500 DE genes up-regulated in human and 500 DE genes down-regulated in human are shown. Data is from DLPFC and only high-confidence ASD-linked genes are used. **C)** Barplot showing the number of high-confidence ASD-linked genes that are up-regulated in human and number of genes that are down-regulated in human relative to chimp in DLPFC L2/3 IT neurons.

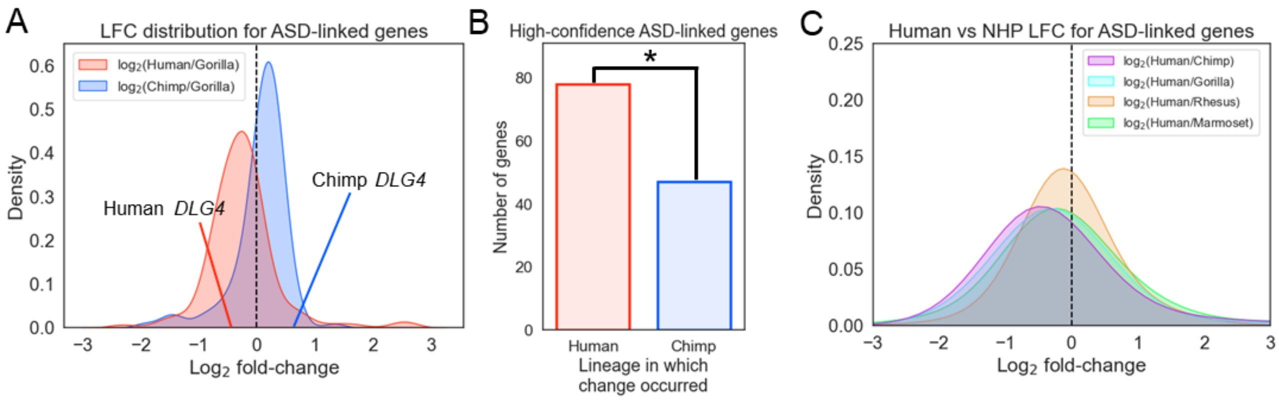


**Supplemental Figure 37: Lower expression of high-confidence ASD-linked genes compared to non-human primates. A)** Distribution of $\log_2$ fold-changes (x-axis) comparing human or chimpanzee to gorilla in L2/3 IT neurons for high-confidence ASD-linked genes with FDR < 0.05 when comparing human and chimpanzee. Only genes with absolute $\log_2$ fold-change less than 3 are shown. The magnitude of the [human/gorilla] divergence was generally greater than the magnitude of the [chimp/gorilla] divergence, suggesting that there has been greater divergence in the human lineage ([human/gorilla] median absolute log2 fold-change = 0.45, [chimp/gorilla] = 0.28, t-test p = 0.00036). **B)** Barplot showing the number of significantly DE high-confidence ASD-linked genes that are human-derived (red) and chimp-derived (blue) in MTG L2/3 IT neurons. * indicates binomial $p < 0.05$. **C)** Distribution of $\log_2$ fold-changes (x-axis) comparing human to chimpanzee, gorilla, macaque, or marmoset in MTG L2/3 IT neurons for high-confidence ASD-linked genes with FDR < 0.05 when comparing human and chimpanzee. Only genes with absolute $\log_2$ fold-change less than 3 are shown.

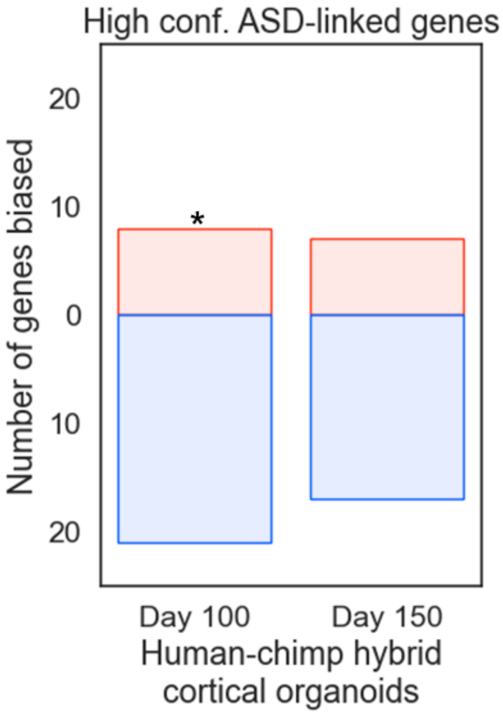


**Supplemental Figure 38: Lower expression of high-confidence ASD-linked genes from the human allele in cortical organoids.** Barplot showing the number of significantly DE high-confidence ASD-linked genes with higher expression from the human allele (red) and higher expression from the chimp allele (blue) in cortical organoids. * indicates binomial p = 0.01.

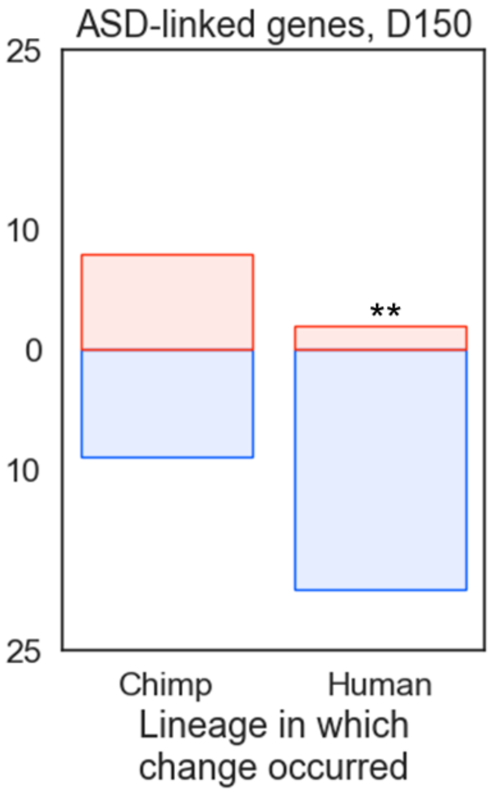


**Supplemental Figure 39: Lower expression of ASD-linked genes from the human allele in cortical organoids.** Barplot showing the number of significantly DE ASD-linked genes with higher expression from the human allele (red) and higher expression from the chimp allele (blue) in day 150 cortical organoids for human-derived and chimp-derived genes separately. ** indicates Fisher's exact test $p < 0.01$. All ASD-linked genes, regardless of SFARI gene score, are used.

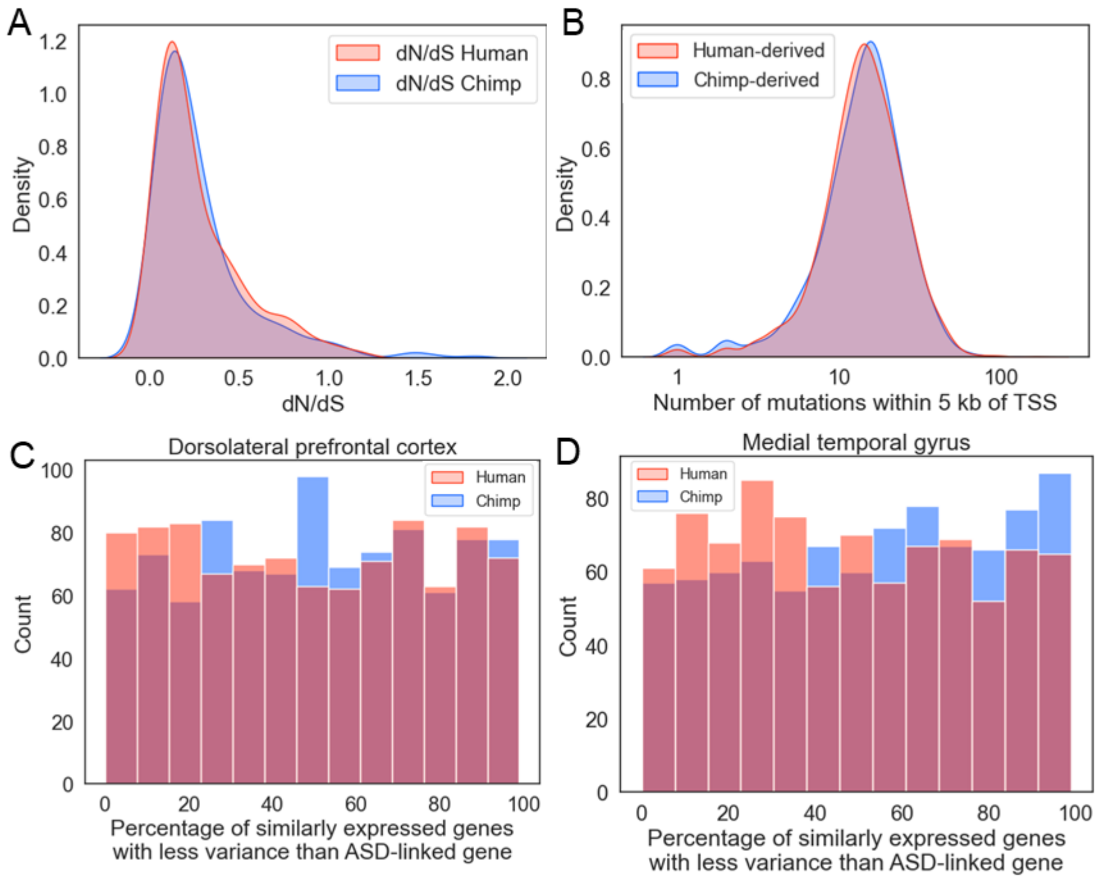


**Supplemental Figure 40: No evidence for loss of constraint on ASD-linked genes in the human lineage. A)** Distribution of per-gene dN/dS in humans (red) and chimpanzees (blue) for all ASD-linked genes with at least one nonsynonymous and one synonymous change in each lineage. **B)** Distribution of number of mutations within five kilobases of the TSS per-gene for all ASD-linked genes for human (red) and chimp (blue). The total number of human-derived genetic differences across all genes was down-sampled to match the total number of chimp-derived changes. **C)** Histogram showing the distribution of the proportion of the 100 genes with closest expression to each ASD-linked gene with lower interindividual expression variance in humans (red) and chimpanzees (blue). **D)** Same as in (C) but for the MTG dataset.

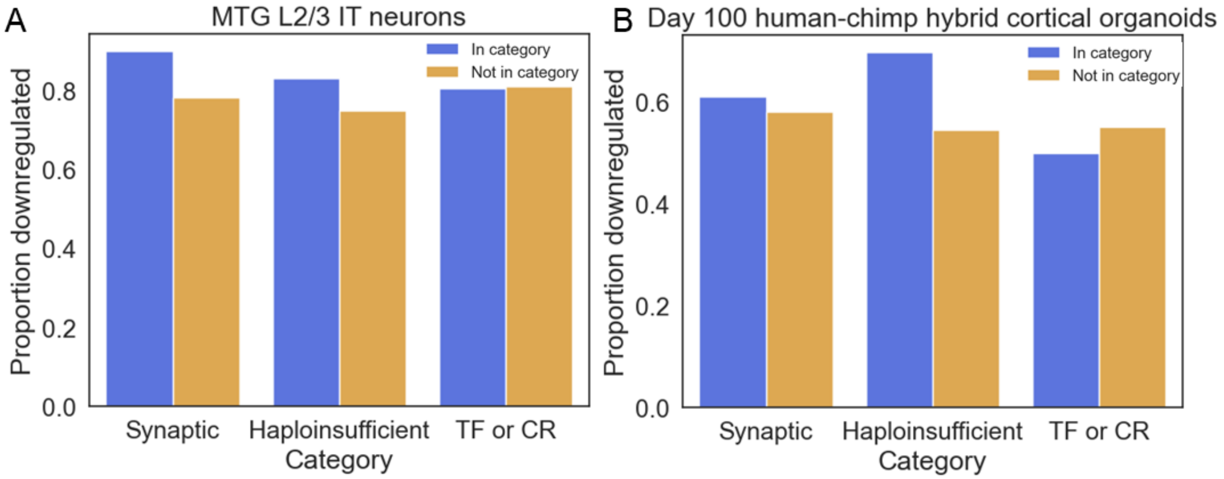


**Supplemental Figure 41: ASD-linked genes tend to be down-regulated regardless of functional category. A)** Barplot showing the proportion of high-confidence ASD-linked genes in a particular category (e.g. synaptic) that are down-regulated compared to the proportion of high-confidence ASD-linked genes not in that category that are down-regulated. The blue bar corresponds to genes in the category of interest and the orange bar to all high-confidence ASD-linked genes not in the category. **B)** Same as in (A) but for the day 100 human-chimp hybrid cortical organoid dataset and using all ASD-linked genes, regardless of confidence level, due to the lower number of differentially expressed genes in this dataset.

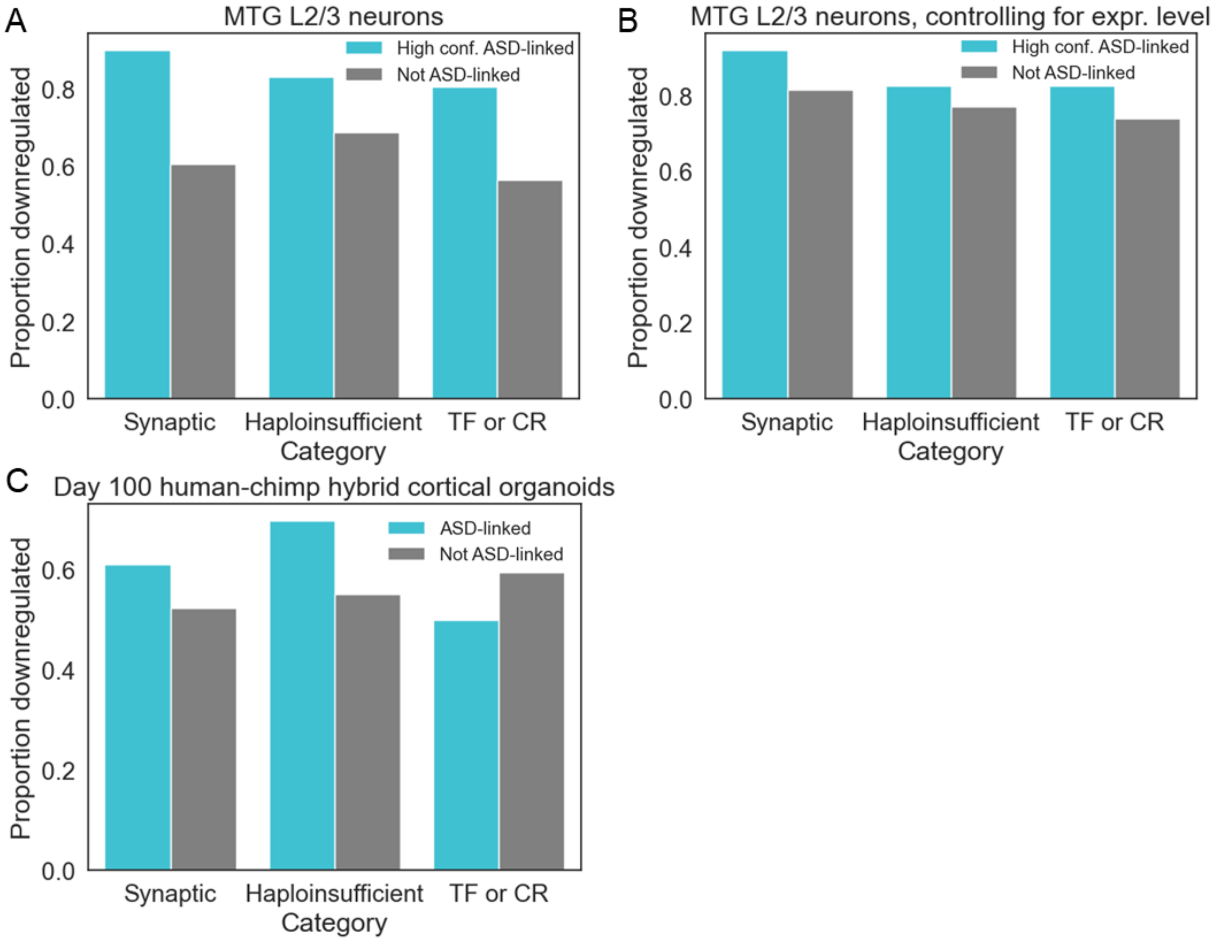


**Supplemental Figure 42: Genes with similar properties to ASD-linked genes tend to be down-regulated. A)** Barplot showing the proportion of high-confidence ASD-linked genes that are in a particular category (e.g synaptic) that are down-regulated compared to the proportion of genes that are not linked to ASD but are in that category that are down-regulated. The blue bar corresponds to high-confidence ASD-linked genes and the grey bar to all genes in the category that are not ASD-linked. Data are from MTG L2/3 neurons. All three comparisons are significantly different at binomial $p = 0.0006$ for synaptic, $p = 0.003$ for haploinsufficient, and $p = 0.004$ for TF or CR; binomial tests used the proportion of down-regulation in a category not linked to ASD as the null probability of down-regulation. **B)** Same as in (A) but controlling for gene expression (Methods). Binomial $p > 0.1$ for all three comparisons. **C)** Same as in (A) but for the day 100 human-chimp hybrid cortical organoid dataset and using all ASD-linked genes, regardless of confidence level, due to the lower number of differentially expressed genes in this dataset. Binomial $p = 0.27$ for synaptic, $p = 0.03$ for haploinsufficient, and $p = 0.48$ for TF or CR.

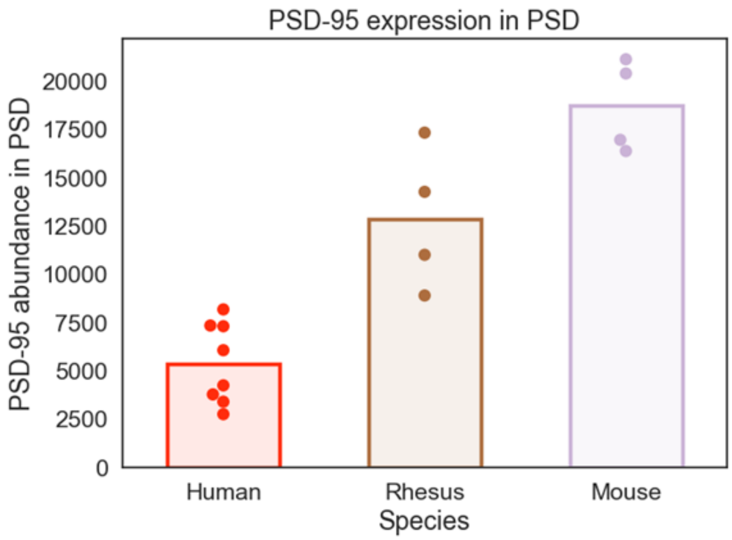


**Supplemental Figure 43: Reduced levels of PSD-95 at human postsynaptic densities.** Protein expression of PSD-95, the protein encoded by *DLG4*, in postsynaptic densities of humans, rhesus macaques, and mice[77].

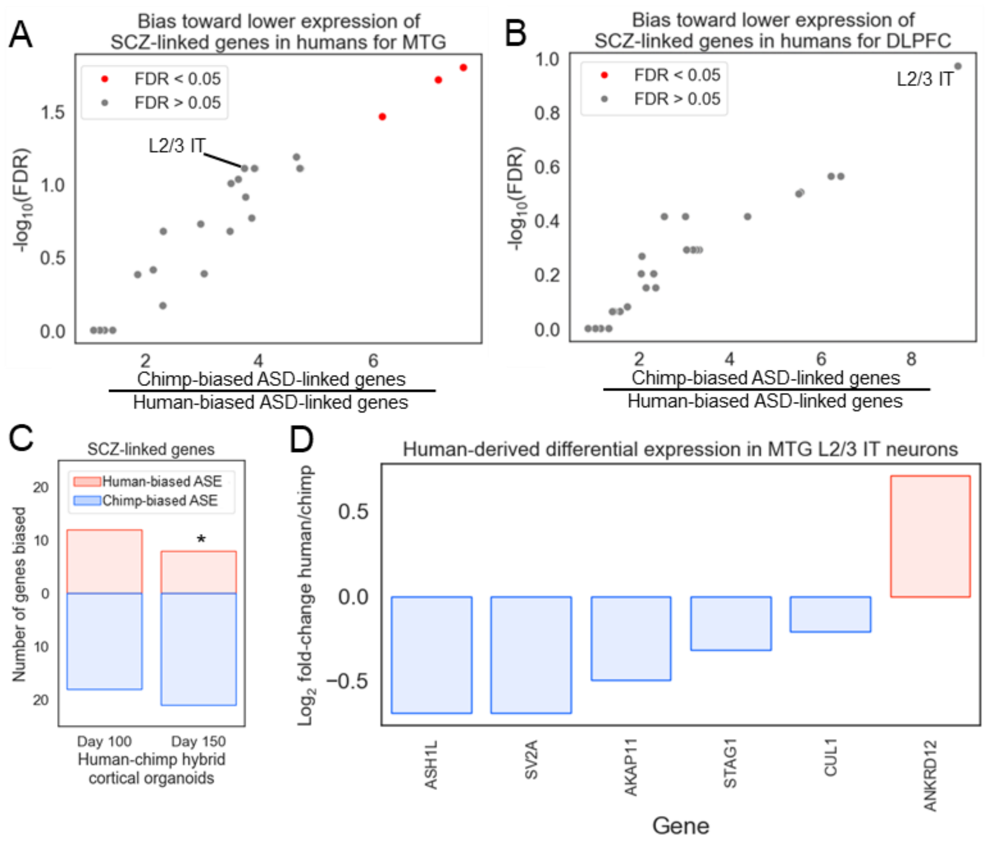


**Supplemental Figure 44: Down-regulation of SCZ-linked genes in humans compared to chimpanzees. A)** Volcano plot showing the fold-enrichment for down-regulation in humans (x-axis) and the -$\log_{10}$ binomial FDR (y-axis). Subclasses with FDR < 0.05 are shown in red; only subclasses with at least 500 DE genes up-regulated in human and 500 DE genes down-regulated in human are shown. Data is from MTG. **B)** Same as in (A) but for the DLPFC dataset. **C)** Barplot showing the number of SCZ-linked genes with higher expression from the human allele (red) or higher expression from the chimp allele (blue) in hybrid cortical organoids. * indicates binomial $p < 0.05$. **D)** Differential expression of SCZ-linked genes in MTG L2/3 IT neurons. Gene names (x-axis) and the $\log_2$ fold-change between human and chimpanzee expression (y-axis) are shown.
